# Genetic Encoding of a Highly Photostable, Long Lifetime Fluorescent Amino Acid for Imaging in Mammalian Cells

**DOI:** 10.1101/2021.04.05.438526

**Authors:** Chloe M. Jones, D. Miklos Robkis, Robert J. Blizzard, Mika Munari, Yarra Venkatesh, Tiberiu S. Mihaila, Alex J. Eddins, Ryan A. Mehl, William N. Zagotta, Sharona E. Gordon, E. James Petersson

## Abstract

Acridonylalanine (Acd) is a fluorescent amino acid that is highly photostable, with a high quantum yield and long fluorescence lifetime in water. These properties make it superior to existing genetically encodable fluorescent amino acids for monitoring protein interactions and conformational changes through fluorescence polarization or lifetime experiments, including fluorescence lifetime imaging microscopy (FLIM). Here, we report the genetic incorporation of Acd using engineered pyrrolysine tRNA synthetase (RS) mutants that allow for efficient Acd incorporation in both *E. coli* and mammalian cells. We compare protein yields and amino acid specificity for these Acd RSs to identify an optimal construct. We also demonstrate the use of Acd in FLIM, where its long lifetime provides strong contrast compared to endogenous fluorophores and engineered fluorescent proteins, which have lifetimes less than 5 ns.

## Introduction

Fluorescence spectroscopy provides a powerful method to monitor protein folding and dynamics in realtime under physiological conditions. One challenge for monitoring these dynamics is the ability to fluorescently label the protein of interest without disrupting its folding and function.^1, 2^ Although widely used, large fluorescent proteins tags such as green fluorescent protein (GFP)^3, 4^ can hinder native protein folding and function,^1, 5, 6^ and affinity labeling such as bioorthogonal (“click”) chemistry^7^ and SNAP^8^ or Halo^9^ tags can be limited by biocompatibility, labeling efficiency and solvent accessibility of the labeling location.^1, 10^ To avoid these issues, a fluorescent label can be incorporated co-translationally, eliminating the need for additional post-translational manipulation and allowing one to label proteins at interior sites.^11^ Incorporation of the labels in living cells is made possible by an engineered aminoacyl tRNA synthetase (RS) that selectively charges an orthogonal tRNA with a non-canonical amino acid (ncAA).^12^ The anticodon loop recognizes an amber stop codon that is introduced in the mRNA chain, leading to the insertion of the ncAA in the desired position of a protein of interest.

The Petersson and Mehl laboratories have previously incorporated acridonylalanine (Acd or δ) in *E. coli* using these amber codon suppression methods.^13-18^ Acd is a small fluorescent amino acid (222 Å^3^), with an emission maximum at 420-450 nm, an extinction coefficient of 5,700 M^-1^cm^-1^ and a quantum yield of 0.98 in aqueous buffer (ESI, Fig. S2 and Table S2) and high photostability (ESI, Fig. S1 and Table S1).^19, 20^ We varies with the polarity of the environment and has been used in fluorescence polarization (FP) assays to investigate the dynamics of protein-protein interactions.^17^ Compared to other visible wavelength fluorescent ncAAs used for *in cellulo* genetic incorporation,^22-25^ hydroxycoumarin ethylglycine (Hco), coumarinyl lysine (CoulLys), dansylalanine (DanAla), and acetylnaphthalenylaminoalanine (Anap), Acd has the highest quantum yield, longest fluorescence lifetime, and greatest photostability in water (ESI, Figs. S1-S9 and Tables S1-S6). The longer lifetime of Acd makes the fluorophore more responsive to molecular weight changes in FP experiments.^17, 26^ It also permits better resolution of multiple populations in lifetime experiments, including single photon counting experiments^27^ and in fluorescence lifetime imaging microscopy (FLIM, Fig. 1).^28^

**Fig. 1.**
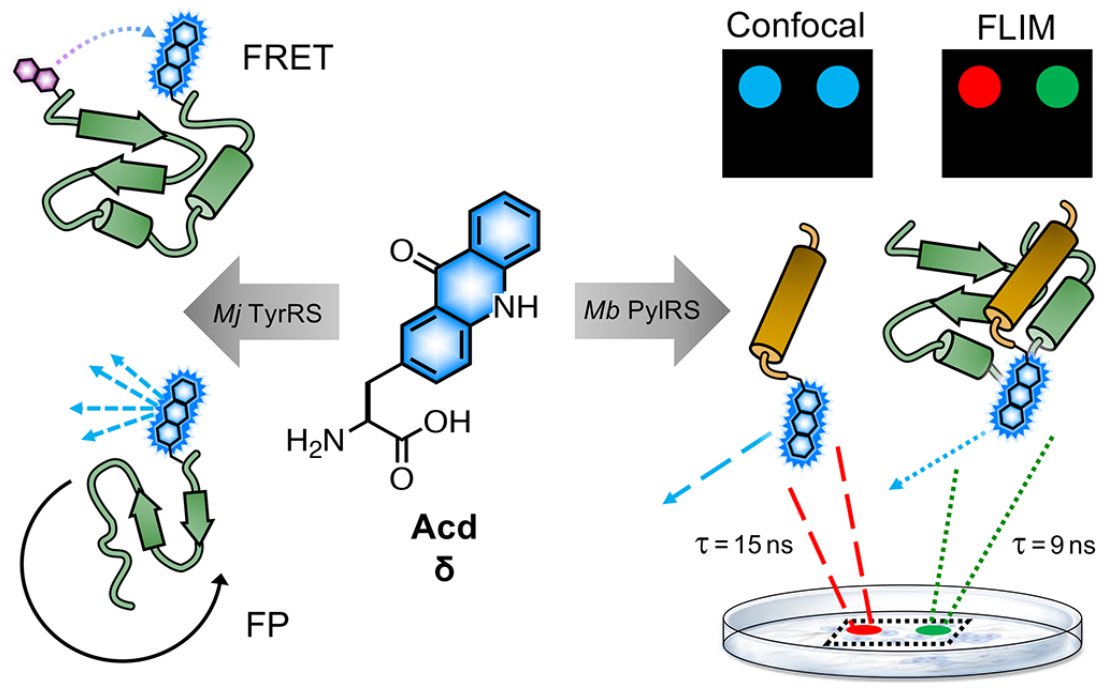
Acd Applications. Acd (or δ) has been previously used in Förster resonance energy transfer (FRET) and fluorescence polarization (FP) to monitor conformational changes and protein binding interactions among purified proteins expressed in *E. coli* using RSs based on *Mj* TyrRS. Development of a *Mb* PylRS-based system for mammalian cell incorporation enables confocal microscopy and fluorescence lifetime imaging (FLIM) studies to observe processes like protein association in live cells. FLIM pixels are coloured by lifetime, (red = 15 ns, green = 9 ns) which is related to the Acd local environment.

To date, all Acd studies relied on *E. coli* protein expression, as our engineered RSs were based on an *Methanocaldococcus jannachii* (*Mj*) tyrosyl RS and thus were only orthogonal to the translation machinery in prokaryotes. In order to use Acd in mammalian cell imaging experiments, including FLIM, we needed an RS that is orthogonal in mammalian cells.

Here, we report the development of an Acd incorporation method that relies on an engineered pyrrolysine (Pyl) RS for site specific protein labeling. PylRS variants from *Methanosarcina bakeri* (*Mb*),^29^ *Methanosarcina mazei* (*Mm*),^29^ and *Methanomethylophilus alvus* (*Ma*)^30, 31^ have been shown to be orthogonal in both *E. coli* and mammalian cells. Using engineered *Mb* PylRS variants that specifically and efficiently incorporate Acd in both *E. coli* and mammalian cells, we also demonstrate that this new labelling platform can be used for FLIM studies in mammalian cells.

## Results

### Acd Synthetase Selection

To identify a *Mb* PylRS/tRNA_CUA_ pair capable of site-specifically incorporating Acd into proteins in response to an amber stop codon (TAG) we screened a library of *Mb* PylRS variants in which five active-site residues were randomized to all 20 amino acids (N_311_, C_313_, V_366_, W_382_, G_387_) using a standard life/death selection in *E. coli*.^32, 33^ Following a round of positive selection in the presence of Acd and a round of negative selection against canonical amino acids, 96 colonies were assessed for their ability to suppress a TAG codon interrupted sfGFP gene (sfGFP-TAG_150_) in the presence of Acd and suppressor tRNA_CUA_. The top 25 performing clones were sequenced and 13 unique RS clones were identified with highly similar active site sequences (Fig. 2 Top and ESI, Table S7). These 13 clones were evaluated for (1) their efficiency (full-length protein yield in presence of Acd), (2) fidelity (level of canonical amino acid misincorporation in absence of Acd) and (3) their efficiency in producing full length protein using *N*-phenyl-amino phenylalanine (Npf), an undesired side product of Acd synthesis that was recognized by first-generation AcdRSs.^13, 14^ Three of these clones showed permissivity for Npf, and were not considered further.

**Fig. 2.**
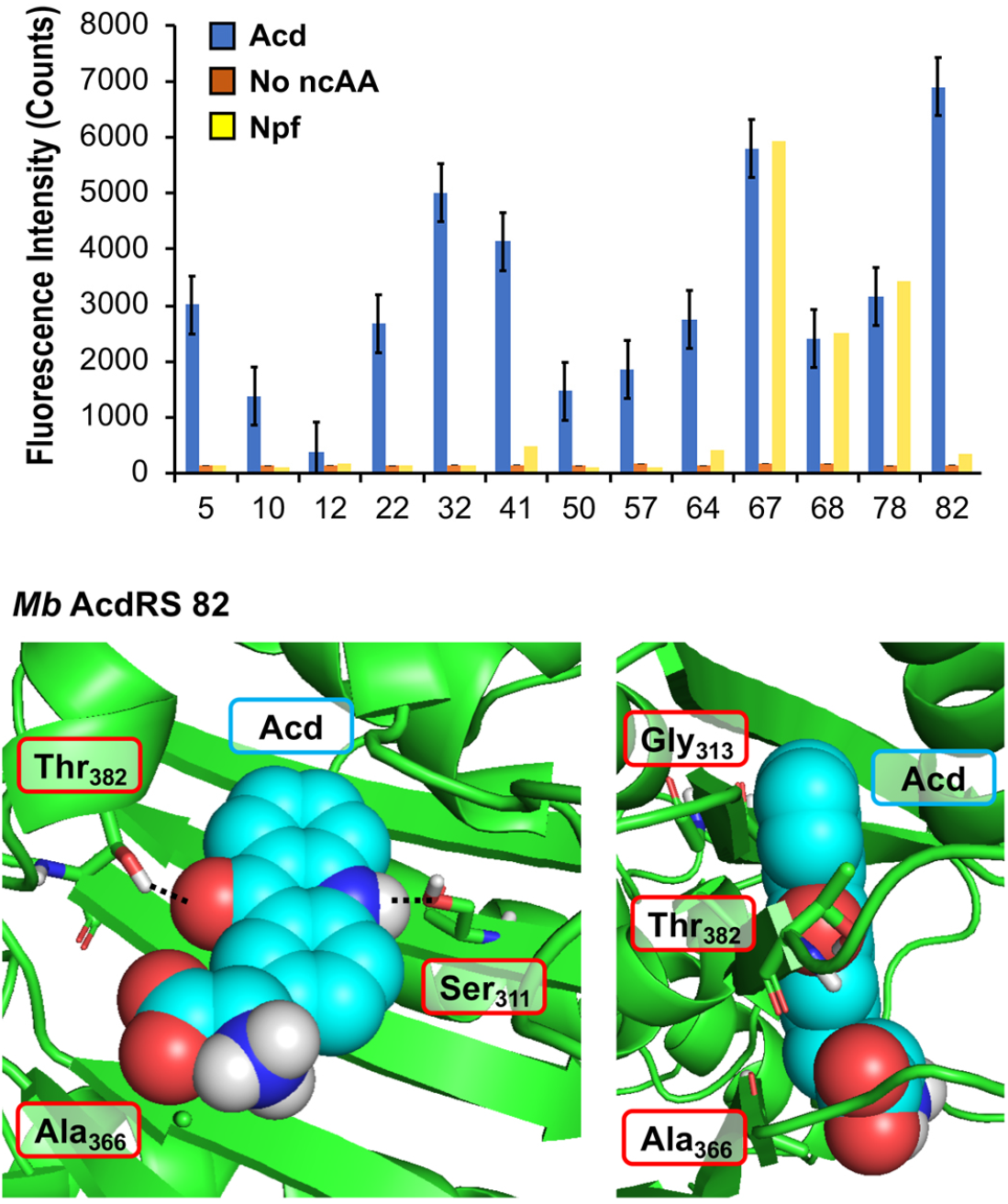
PylRS Mutants for Acd Incorporation. Top: Fluorescence measurements of sfGFP reporter to evaluate RS efficiency. Orange, blue, and yellow represent fluorescence from colonies induced in media containing no ncAA, 1 mM Acd, or 1 mM Npf, respectively. Cultures of 500 μL were grown for 48 hours before 4-fold dilution of suspended cells with buffer for fluorescence measurement at 528 nm (485 nm excitation) using a platereader. RSs enabling high sfGFP fluorescence in Acd conditions with low fluorescence in no ncAA or Npf conditions were selected for further characterization. Additional data showing sfGFP fluorescence for all clones are shown in ESI, Fig. S10. Bottom: A homology model of *Mb* AcdRS 82 based on the x-ray crystal structure of *Mm* PylRS (PDB ID: 2ZIN)^35^ with Acd docked in the active site using Rosetta. Key points of interaction with Acd are shown from two angles. Sites of mutation relative to parent *Mb* PylRS are labeled in red boxes. A model of *Mb* AcdRS 41 is shown in ESI, Fig. S11.

The most active and selective variants (32, 41, 82) had similar mutations in the active site (N_311_S and V_366_A, as well as W_382_T or W_382_V). *Mb* AcdRS 32 and 82 also had an additional active site mutation of C_313_A or C_313_G, respectively, and *Mb* AcdRS 82 also contained a non-active site mutation, L_155_V. AcdRS 41 and AcdRS 82 were selected for further evaluation as 82 was most active and 41 featured the same active site mutations, except it retained Cys_313_ of the parent RS. The two RSs were cloned into both pDULE2^35^ and pUltra^36^ plasmids, under respective lpp and tacI promoters, each also containing tRNA_CUA_. We did not retain the L_155_V mutation since other active sequences (e.g. 75) showed activity without the mutation. If necessary, Npf incorporation concerns can be eliminated by using an alternative Acd synthetic route.^13^

### Structural Modeling

In order to better understand the effects of the mutations in the selected mutants, we built a homology model of *Mb* AcdRS 82 based on a x-ray crystal structure of substrate-bound *Mm* PylRS (PDB ID: 2ZIN).^34^ With the exception of one loop region, there is very high active site sequence homology between the *Mm* and *Mb* PylRS (ESI, Fig. S12), and it has been demonstrated that the conservation of the active site allows mutations for ncAA-selective variants to be swapped between the two types of PylRS.^37-41^ Thus, we expect the homology model to be quite accurate. Acd was docked into the *Mb* AcdRS 82 active site in Rosetta, followed by relaxation of the structure (Fig. 2 Bottom) A model of *Mb* AcdRS 41 was generated by a G_313_C mutation of *Mb* AcdRS 82, followed by relaxation (ESI, Fig. S11).

One can rationalize the effects of the four mutations in AcdRS 82 as follows: Asn_311_ to Ser creates space in the active site and provides a hydrogen-bond acceptor for the Acd sidechain N-H, Trp_382_ to Thr removes a steric clash and provides a hydrogen-bond donor for the Acd ketone, Cys_313_ to Gly removes a steric clash with the distal ring of Acd and allows it to hydrogen-bond with both Ser_311_ and Thr_382_ together, and Val_366_ to Ala removes a steric clash. The existence of the hydrogen bond bridge in the AcdRS 82 model nicely explains its superior activity relative to AcdRS 41.

### Protein expression in E. coli and characterization of Acd incorporation

From the initial screen, *Mb* AcdRS 41 and *Mb* AcdRS 82 were subjected to further investigation of their incorporation efficiency and ability to mis-incorporate canonical amino acids at the TAG site. To this end we expressed TAG-bearing constructs of α-synuclein (αS) and calmodulin (CaM) to investigate the RSs in the context of an intrinsically disordered protein (αS) where the tertiary structure of the protein should not affect incorporation, and a structured protein (CaM) where destabilization by Acd incorporation could decrease protein expression.^18^ We have previously studied conformational changes and protein/protein interactions in both proteins using Acd.^13, 16^ Each protein construct was fused to a C-terminal GyrA intein containing a six histidine (His_6_) tag. This fusion construct provides a way to eliminate potential protein truncations, as the intein with the His_6_ tag is only produced for the full length protein and can be cleaved tracelessly using β-mercaptoethanol.^42^

To compare RS efficiency and specificity, we first expressed αS using a plasmid encoding αS with a TAG codon replacing Phe_94_ or Glu_114_ as well as a plasmid encoding either *Mb* AcdRS 41 *Mb* AcdRS 82 and the cognate tRNA_CUA_. We compared incorporation efficiency, based on isolated yields, and amino acid specificity, based on matrix-assisted laser desorption ionization (MALDI) mass spectrometry (MS) data. Phe_94_ and Glu_114_ were chosen as the incorporation sites as they have been shown to tolerate a variety of bulky fluorophore substitutions, including both ncAA incorporation and Cys modification.^16, 43-50^ As a positive control, we expressed the same protein construct using an *Mj* RS previously optimized for Acd incorporation in *E. coli (Mj* AcdRS A9).^14^ As an additional comparison of anticipated masses and yields, we also expressed the wild-type (WT) αS sequence fused to the His_6_ intein tag. This serves as an approximate reference mass for any mis-incorporation at the TAG site due to charging of canonical amino acids. Given the structural similarity of Acd to the aromatic amino acids Phe, Trp, and Tyr we anticipated that there was a potential for mis-incorporation of these amino acids. Indeed, Phe mis-incorporation has been previously observed in PylRS mutants selected for incorporation of aromatic unnatural amino acids,^52^ although the selectivity of engineered PylRSs has been reported to be generally good.^53^

Protein expressions in *E. coli* were performed with either pDULE2 or pUltra plasmids, using BL21 DE3 or C321.deltaA cells (release factor 1 removed and all 321 UAG codons changed to improve ncAA incorporation),^54^ respectively. Protein levels were monitored via gel throughout cell lysis, nickel purification and intein cleavage. SDS-PAGE analysis confirmed that full-length Acd-containing αS was produced for both *Mb* AcdRS 41 and *Mb* AcdRS 82 using either the pDULE2/BL21 and pUltra/C321 expression methods. However, MALDI MS analysis of purified whole protein constructs demonstrated that significant mis-incorporation of a canonical amino acid occurred with pUltra/C321 with either *Mb* AcdRS 41 and *Mb* AcdRS 82 (ESI, Fig. S19). In contrast, MALDI MS data confirmed that both *Mb* AcdRS 41 and *Mb* AcdRS 82 exclusively charge tRNA_CUA_ with Acd for pDULE2/BL21 expressions. (Fig. 3 and ESI, Figs. S17-S18). Whole protein masses matching the anticipated mass for Acd for incorporation at position 114 in αS are shown in Table 1. No evidence of Npf incorporation was observed.

**Fig. 3.**
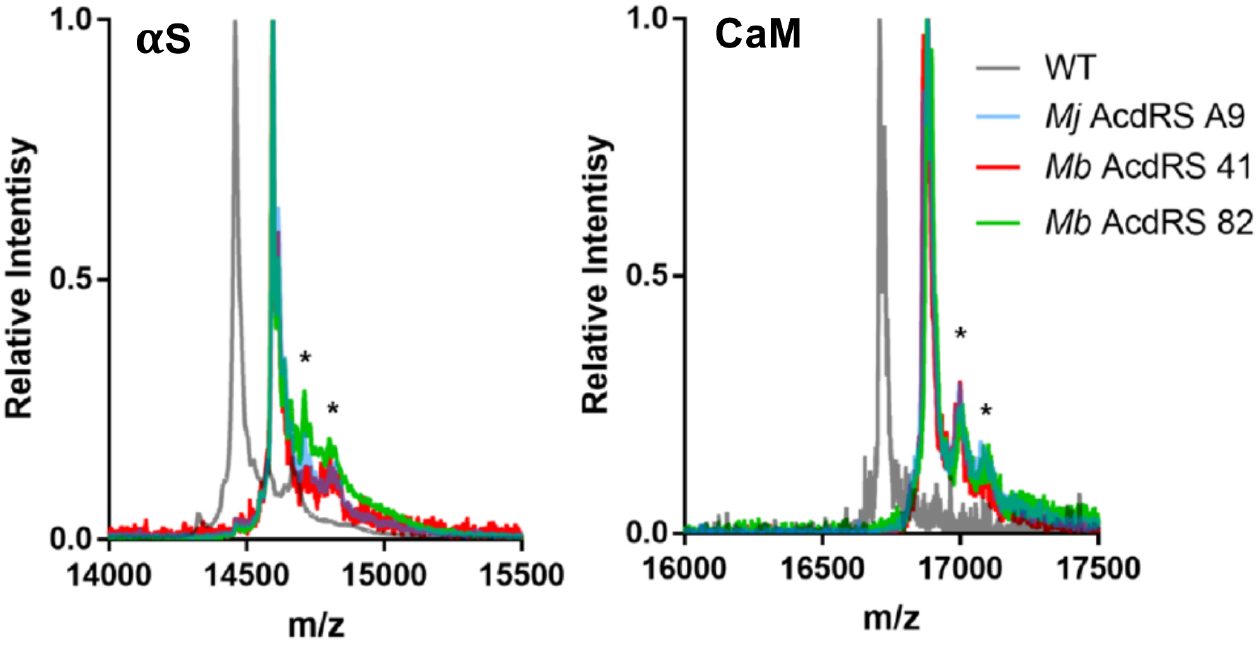
MALDI MS Analysis of Acd Incorporation. Expressions of αS-E_114_δ or CaM-L_113_δ using *Mj* AcdRS A9, *Mb* AcdRS 41, or *Mb* AcdRS 82 were performed in parallel. WT αS and CaM were also expressed for comparison. MALDI MS data showed masses corresponding to Acd incorporation with no peaks corresponding to mis-incorporation of a canonical amino acid (approximately equal to WT mass). *indicates a previously described MALDI matrix adduct.^53^

**Table 1.**
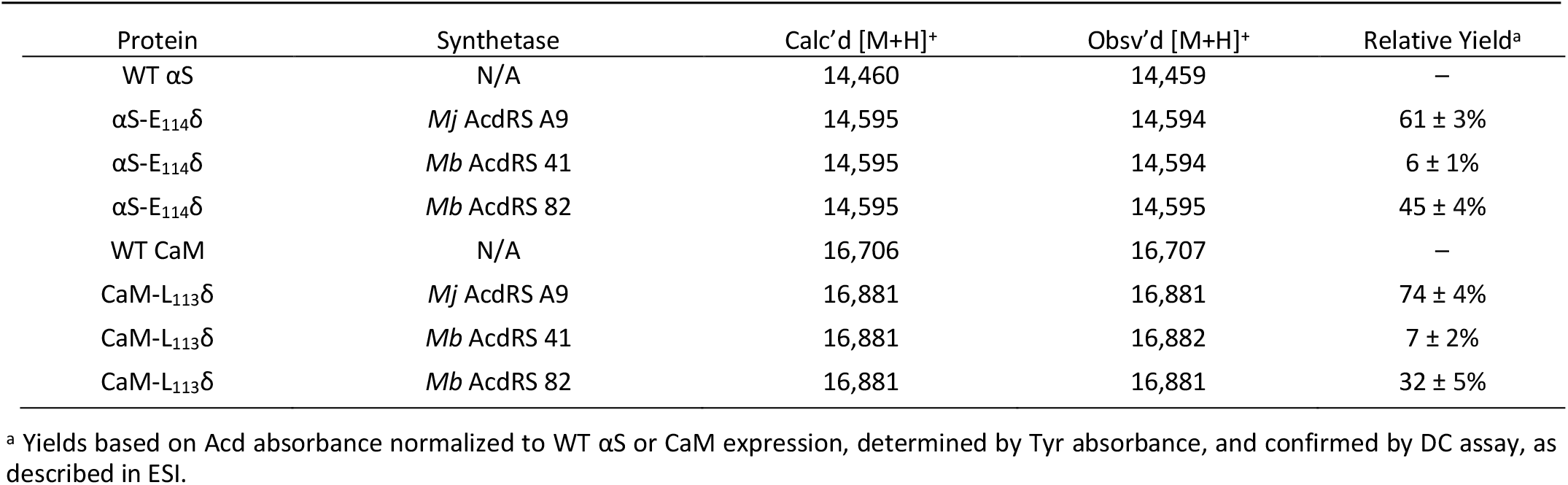
Masses and yields of proteins from three rounds of *E. coli* Acd expression trials.

| Protein | Synthetase | Calc'd [M+H]<sup>+</sup> | Obsv'd [M+H]<sup>+</sup> | Relative Yield<sup>a</sup> |
| --- | --- | --- | --- | --- |
| WT αS | N/A | 14,460 | 14,459 | — |
| αS-E<sub>114</sub>δ | *Mj* AcdRS A9 | 14,595 | 14,594 | 61 ± 3% |
| αS-E<sub>114</sub>δ | *Mb* AcdRS 41 | 14,595 | 14,594 | 6 ± 1% |
| αS-E<sub>114</sub>δ | *Mb* AcdRS 82 | 14,595 | 14,595 | 45 ± 4% |
| WT CaM | N/A | 16,706 | 16,707 | — |
| CaM-L<sub>113</sub>δ | *Mj* AcdRS A9 | 16,881 | 16,881 | 74 ± 4% |
| CaM-L<sub>113</sub>δ | *Mb* AcdRS 41 | 16,881 | 16,882 | 7 ± 2% |
| CaM-L<sub>113</sub>δ | *Mb* AcdRS 82 | 16,881 | 16,881 | 32 ± 5% |

<sup>a</sup> Yields based on Acd absorbance normalized to WT αS or CaM expression, determined by Tyr absorbance, and confirmed by DC assay, as described in ESI.

Final protein yields were calculated based on molar absorptivity and confirmed by detergent compatible (DC) protein quantification assay, showing that *Mb* AcdRS 82 incorporates Acd into αS more efficiently than *Mb* AcdRS 41 (Table 1 and ESI, Tables S8-S9, gel analysis in Fig. S13-S16). The misincorporation of a natural amino acid under pUltra/C321 expression conditions may be a function of differences in culture media or the increased permissivity of the translation system in the C321 cells.^54^ No evidence of mis-incorporation was observed in multiple expression trials with the pDULE2/BL21 method. CaM-L_113_δ yields in pDULE2/BL21 expression studies followed a similar trend for *Mb* AcdRS 41 and *Mb* AcdRS 82. Similar to αS expressed using pDULE2/BL21, no evidence of mis-incorporation was seen for either RS (Fig. 3). Protein yields calculated by molar absorptivity and confirmed by DC assay again showed that *Mb* AcdRS 82 is more active (Table 1 and ESI, Table S2). Together, the αS and CaM results show that both *Mb* AcdRS 41 and *Mb* AcdRS 82 can incorporate Acd into proteins with high amino acid specificity using the pDULE2/BL21 expression system. However, *Mb* AcdRS 82 does this more efficiently to produce roughly 5-fold more purified protein. Excited by these high yields, we moved forward to study Acd incorporation in mammalian cells using *Mb* AcdRS 82. In addition, we wished to test whether *Mb* AcdRS 82 is more active than *Mb* AcdRS 41 in mammalian cells as it is in *E. coli* cells.

### Mammalian cell expression screening

The incorporation efficiency of both of the *Mb* AcdRS variants was examined in mammalian cells. *Mb* AcdRS 41 and *Mb* AcdRS 82 were separately cloned into the pAcBac1 tR4-MbPyl vector, which contains two copies of a cognate *Mb* PylRS RNA_CUA_.^55^ These plasmids were transfected into HEK293T/17 cells together with a dominant negative eukaryotic release factor one construct (DN-eRF1) that enhances ncAA expression yields in mammalian cells^56^ as well as a vector encoding enhanced GFP (EGFP) with a TAG codon at position 40 (EGFP-Y_40_δ). RS efficiency and specificity were tested by measuring GFP fluorescence in the presence or absence of Acd for each *Mb* AcdRS. Transfections in the absence pAcBac1 tR4-AcdRS provided an additional negative control for read-through of the TAG codon in the absence of a cognate RS and tRNA_CUA_.

The trend of incorporation efficiency for the RSs observed in *E. coli* held in that GFP fluorescence for *Mb* AcdRS 82 was 3-fold higher than *Mb* AcdRS 41. Additionally, both RSs showed low GFP fluorescence in the absence of Acd, indicating little to no mis-incorporation of natural amino acids (Fig. 4). Based on these results and our studies in *E. coli*, we selected *Mb* AcdRS 82 for further mammalian cell experiments. We also purified EGFP using a C-terminal His_6_ tag and analysed incorporation fidelity using electrospray ionization (ESI) MS. We found no evidence of misincorporation of Npf with either RS, but trace incorporation of a natural amino acid using AcdRS41 (ESI, Fig. S26). Based on these results and our studies in *E. coli*, we selected *Mb* AcdRS 82 for further mammalian cell experiments.

**Fig. 4.**
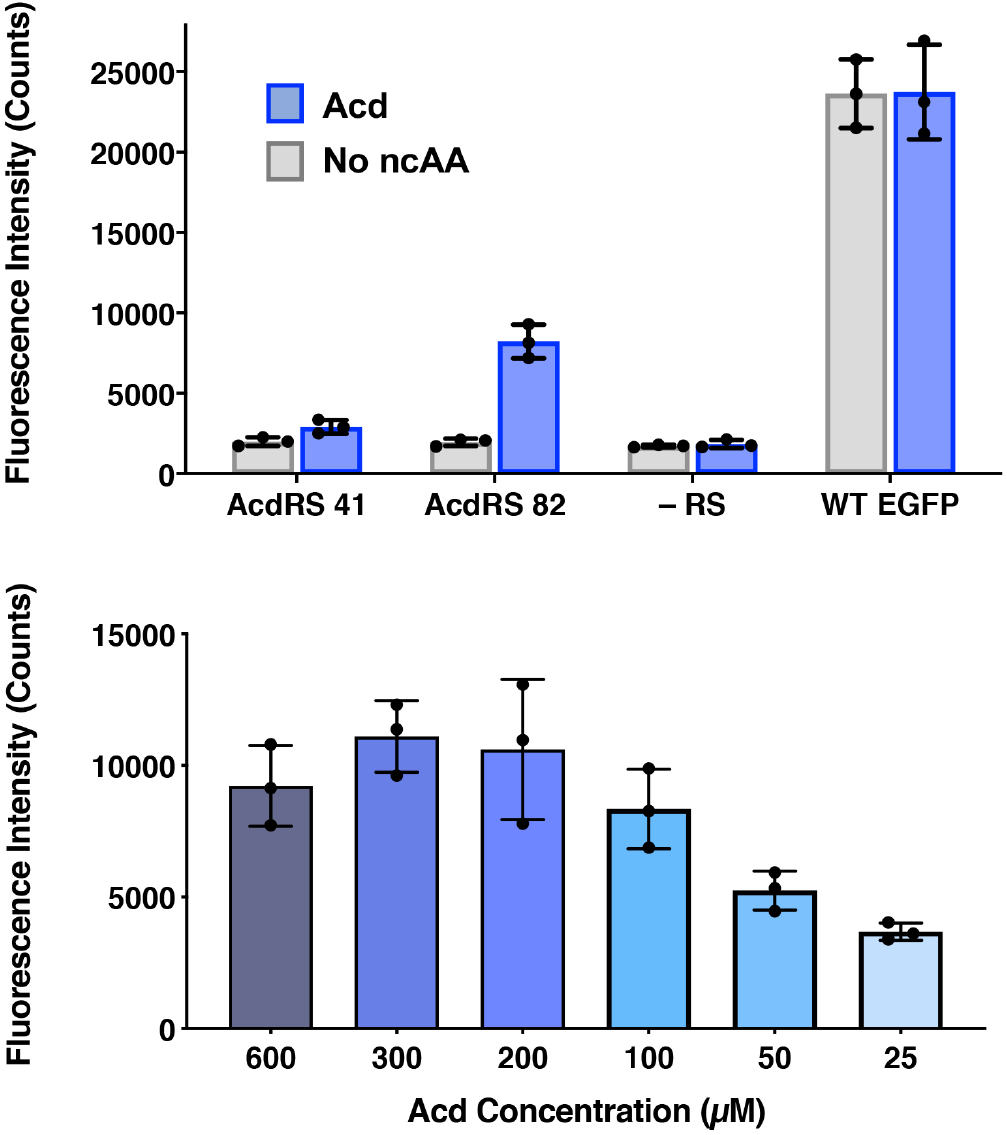
Lysate Fluorescence Analysis of Acd Incorporation in Mammalian Cells. Top: Fluorescence measured from expression of EGFP-Y_40_δ using *Mb* AcdRS 41 or 82 in HEK cells with 300 μM Acd or no ncAA added. As a negative control, the EGFP plasmid was transfected without the AcdRS and tRNA_CUA_ plasmid (-aaRS). WT EGFP was separately transfected as a positive control. In all conditions, DN-eRF1 was also transfected. n = 3. Bottom: Expression of EGFP-Y_40_δ using *Mb* AcdRS 82 was performed as above, with varying concentrations of Acd. n = 3. Cell images are shown in ESI, Figs. S21-S22.

To determine optimal Acd concentrations for imaging experiments, we expressed EGFP-Y_40_δ using AcdRS82 with varying concentrations of Acd. We found that maximal EGFP fluorescence was observed with 300 μM Acd (Fig. 4, Bottom and images in ESI, Fig. S22). We also performed a cell viability study with varying Acd concentrations and found that Acd was toxic to cells at concentrations above 600 μM, possibly explaining the decrease in EGFP expression at 600 μM Acd (ESI, Fig. S20). Cell imaging revealed that the cell morphology was normal at 300 μM (ESI, Fig. S25). Finally, we varied the concentrations of transfection reagents and plasmids and tested the effect of DN-eRF1 to find optimal expression conditions (ESI, Figs. S23-S24). Comparing to WT EFGP expression based on both imaging and lysate fluorescence measurements in these experiments demonstrated that suppression was 29% efficient under optimal conditions, using *Mb* AcdRS 82 with DN-eRF1, 300 μM Acd, and 1.6 μg DNA (for a 6 well dish) at a 1:5 DNA to Lipofectamine 2000 ratio, with imaging after 24-48 h.

### Mammalian cell fluorescence microscopy

To investigate whether Acd can be used as a fluorescence reporter for protein localization in mammalian cells, we incorporated Acd at TAG mutants in three fusion proteins of varying localization, insulin receptor fused to GFP (IR-K_676_δ-GFP), hyperpolarization-activated cyclic nucleotide–gated ion channel fused to yellow fluorescent protein (spHCN-W_355_δ-YFP), and maltose binding protein (MBP-E_322_δ). The GFP and YFP fusions are C-terminal, so production of the fluorescent protein ensures that suppression of the stop codon was successful. Similar to the GFP screening transfection, *Mb* AcdRS 82 and DN-eRF1 plasmids were cotransfected into HEK293T/17 cells along with a vector encoding the protein of interest with a TAG codon at a site previously shown to permit bulky amino acids.^57-59^ After washout of unincorporated Acd, we imaged Acd-containing insulin IR-K_676_δ-GFP, spHCN-W_355_δ-YFP, and MBP-E_322_δ in live cells with confocal microscopy.

The localization of these proteins is known to be membrane bound in the case of IR-GFP and spHCN-YFP, and cytosolic for MBP. Confocal imaging for IR-K_676_δ-GFP and spHCN-W_355_δ-YFP revealed that Acd fluorescence was localized to the membrane and colocalized with YFP or GFP fluorescence, respectively (Fig. 5). This demonstrates Acd’s ability to be used as a fluorescent probe for protein localization. We additionally imaged cells containing the protein construct and Acd in the absence of the pAcBac1 tR4-MbAcdRS82 plasmid (Fig. 5, no RS conditions). Confocal imaging revealed little or no fluorescence in the GFP or YFP channels, which indicates that no substantial readthrough of the TAG codon occurred. We observed only minor fluorescence in the Acd channel. This indicates that our washout protocols are effective and the remaining fluorescence may be due to endogenous fluorophores such as NADH or tRNA loaded with Acd, which cannot be washed out of the cell (wider fields of view shown in ESI, Fig. S27). Note that the rounded morphology of the cells is not a result of Acd toxicity, but of the extensive washing to remove Acd, as demonstrated by applying the same protocol to cells in the absence of Acd (ESI, Fig. S27). From these studies, we conclude that Acd incorporation is specific to our protein of interest and requires *Mb* AcdRS 82, as expected.

**Fig. 5.**
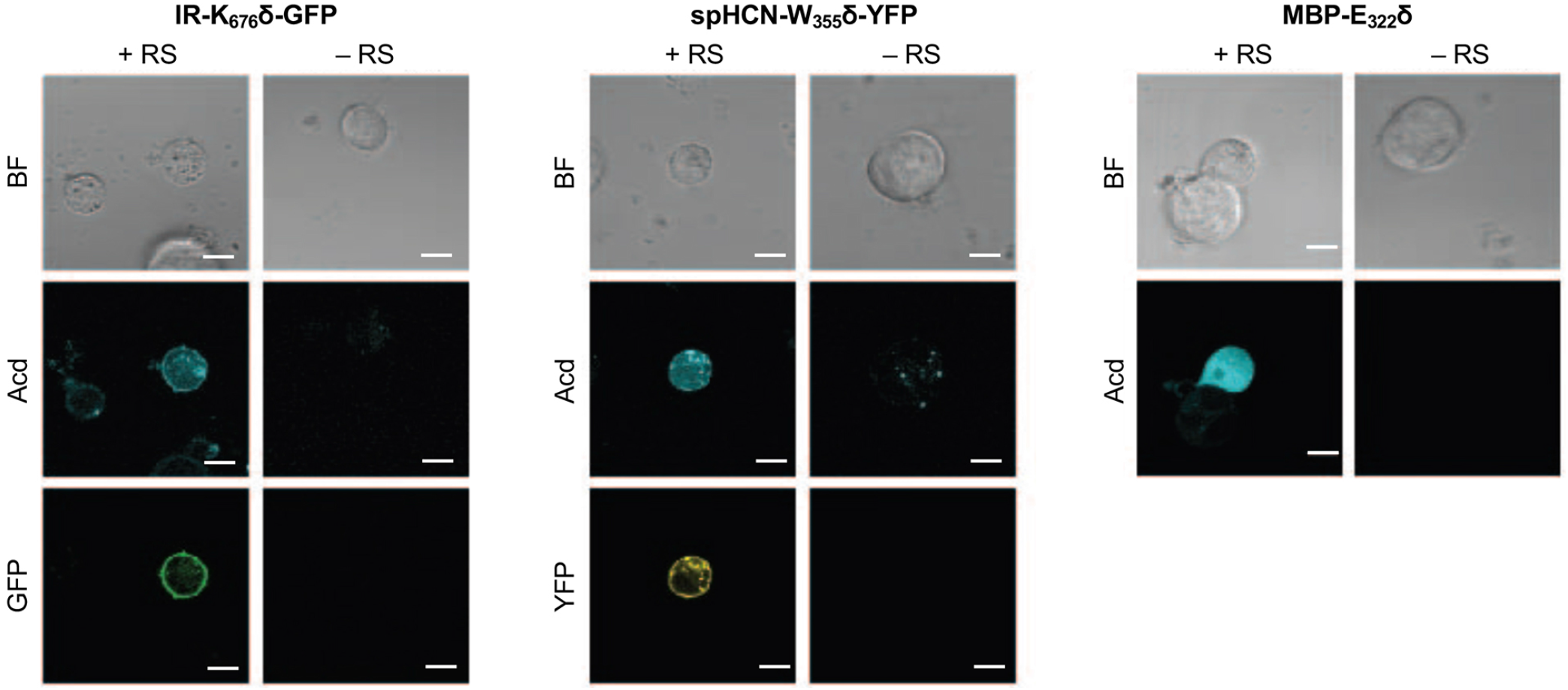
Acd Incorporation into Proteins in Live HEK Cells. Cells were transfected with the indicated plasmid for the protein of interest with a TAG codon and a plasmid encoding a dominant negative release factor to suppress protein truncation, with (+ RS) or without (– RS) a plasmid encoding *Mb* AcdRS 82 and tRNA_CUA_, Confocal microscopy images show brightfield (BF), the Acd fluorescence channel (405 nm excitation; 409-480 nm emission), and the GFP (488 nm excitation; 500-621 nm emission) or YFP (514 nm excitation; 519-621 nm emission) channels. Scale bar = 20 μm.

### Mammalian cell fluorescence lifetime imaging

Since Acd is particularly useful as a FLIM probe, as a proof of concept, we monitored the localization of Acd labeled insulin receptor (IR-K_676_δ-GFP) in HEK293T/17 cells using FLIM. FLIM spatially resolves the fluorescence decay rate of a sample to produce an image based on its fluorescence lifetime. Endogenous fluorophores typically found in a cell have a fluorescence lifetime in the range of 0.3 – 7.5 ns.^28^ Since Acd has a fluorescence lifetime of 15-16 ns, our imaging method is free of background from endogenous cellular fluorophores. For this study, we transfected vectors containing *Mb* AcdRS 82, DN-eRF1, and IR-K_676_δ-GFP into HEK293T/17 cells. Imaging the cells using intensity alone while exciting at 375 nm (Fig. 6) shows Acd fluorescence as well as autofluorescence, likely from NAD and NADH. Highlighting the pixels in which the majority of the intensity was due to Acd (lifetime >15 ns) in red allows the membrane localization of IR-K_676_δ-GFP to be discerned and also eliminates the autofluorescence signal. Highlighting the pixels with a short lifetime (<4 ns) in blue reveals the autofluorescence in all of the cells. The long lifetime pixels (red) compare well with emission from the GFP tag (excitation at 488 nm). The pronounced autofluorescence signal in Fig. 6 compared to Fig. 5 is due to differences in the excitation wavelength used in each experiment. We note that the images in Fig. 6 seem to capture IR-K_676_δ-GFP in different stages of maturation where it is observed in what appears to be ER and/or Golgi in addition to the plasma membrane. Additional FLIM data with spHCN-W_355_δ-YFP highlighting long lifetime Acd emission are shown in Fig. S28 in ESI.

**Fig. 6.**
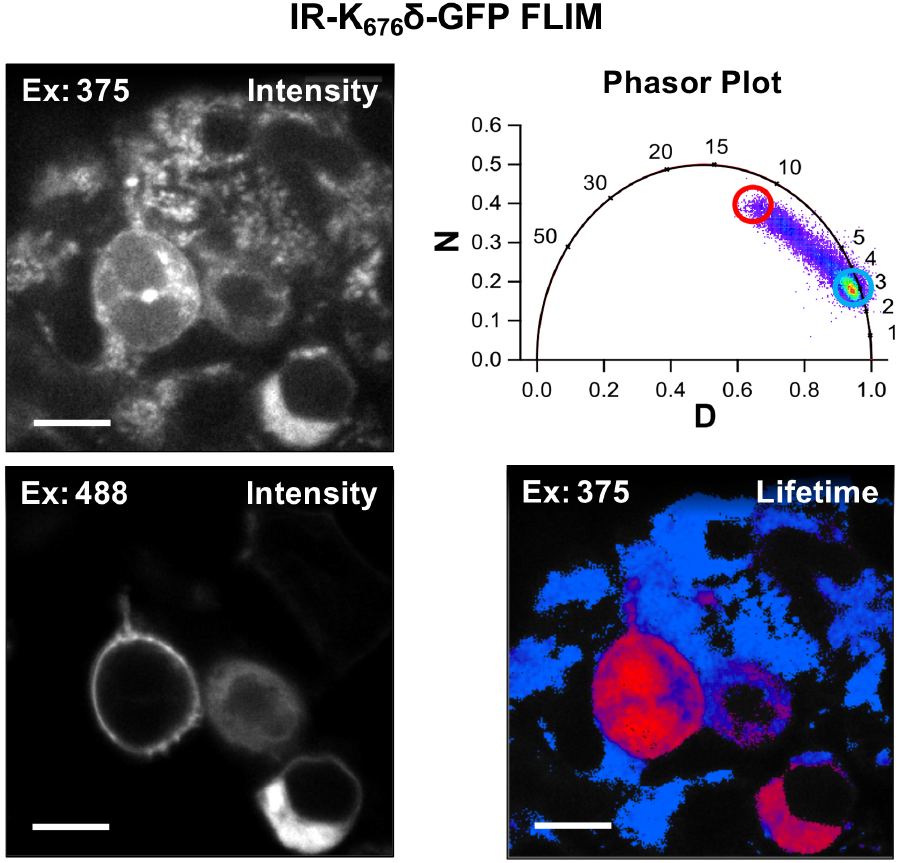
Acd Fluorescence Lifetime Imaging (FLIM). FLIM allows Acd fluorescence to be separated from autofluorescence in living cells. IR-K_676_δ-GFP was expressed with *Mb* AcdRS 82 in HEK cells. Intensity plots (left) show emission after excitation at 375 or 488 nm. Phasor plot (top right) shows phase vectors D and N (see ESI for details) plotted for excitation at 375 nM with 10 MHz modulation and points representing lifetimes of pixels. Circles indicate pixels with a large fraction of lifetime ≥ 15 ns (red) and pixels with a large fraction of lifetimes <4 ns (blue) in the lifetime image (bottom right).

## Discussion

The development of a PylRS-based system for incorporating Acd is valuable for a number of reasons, including the applications in confocal microscopy and FLIM described here, as well as the potential for double-labelling of proteins using mutually orthogonal RSs deriving from PylRS and *Mj* TyrRS. Acd has a higher quantum yield and greater photostability in buffer than any of the other genetically encodable ncAAs (ESI, Figs. S1-S9 and Tables S1-S6, quantum yield is comparable to Hco). Thus, Acd is in many ways superior for imaging, although this depends on the application. For example, the high level of environmental sensitivity depending on the protonation state of Hco is valuable for probing changes in local environment, but may confound other types of studies like FRET. Alternatively, the large Stokes shift of DanAla is useful for eliminating background scatter in spite of DanAla’s very low quantum yield in water (0.10, see ESI, Table S2). Acd is particularly well-suited to FLIM studies because of its 15-16 ns lifetime in water, as most endogenous fluorophores have lifetimes ≤7 ns allowing one to exclude their fluorescence by gating lifetime data.^60^ This makes Acd a superior FLIM probe compared to GFP variants, none of which have comparably long lifetimes (<5 ns).^61, 62^ The other genetically-encodable ncAAs all have shorter lifetimes, except DanAla in hydrophobic environments. However, the variation of the DanAla lifetime from 3 ns in water to 10-12 ns in MeOH, DMSO, and THF (ESI, Table S5) is a less useful range than that of Acd, which varies from ~15 ns in water to ~8 ns in organic solvent, only dropping to ~4 ns in the very hydrophobic environment of THF (ESI., Table S3). Although we did not demonstrate it here, Acd’s lifetime also makes it useful for anisotropy imaging, where the size range distinguishable by a fluorophore increases with fluorophore lifetime.^63^

Beyond FLIM applications, *Mb* AcdRS 82 can potentially be used to incorporate a photo-activatable version of Acd developed by Xiao and coworkers.^64^ *Mb* AcdRS 82 could also be paired with an *E. coli* TyrRS engineered to encode *p*-azidophenylalanine in mammalian cells.^65^ This would allow *in cellulo* fluorescent labelling of a second site with “clickable dyes” opening access to site-specific controlled addition of FRET partners for Acd. Several strategies for encoding two amino acids have been published, including the use of ochre or opal codons, as well as four base codons.^65-67^ Chin and coworkers have also shown that one can generate mutually orthogonal RS/tRNA pairs from PylRS variants.^68^ We will pursue these strategies where a coumarin derivative (Acd FRET donor) or BODIPY derivative (Acd FRET acceptor) could be attached.^13^ Finally, while Acd’s properties are valuable, its low extinction coefficient and blue wavelength emission are not ideal for some microscopy applications, and we will attempt to obtain *Mb* AcdRS 82 variants that can encode brighter and/or red-shifted Acd analogs such as aminoacridonylalanine.^21^ We expect that the PylRS may be better for this than the *Mj* TyrRS as it has been shown to accommodate many large amino acids.^53^

## Conclusion

Acd has been previously demonstrated to be useful as a fluorescent marker and as a biophysical probe to monitor protein conformations without perturbing protein structure. Prior to this study, Acd labelling was limited to proteins expressed in *E. coli* and these proteins were used for *in vitro* biophysical experiments. Here, we have demonstrated the capability of two PylRS variants to efficiently and selectively incorporate Acd into several proteins in *E. coli* and mammalian cells. Notably, the experiments were conducted in multiple laboratories and consistent results were observed. *Mb* AcdRS 82 afforded higher levels of protein expression in both cell types with excellent fidelity, as confirmed by MALDI and ESI MS, and was used for proof-of-concept fluorescence microscopy and FLIM studies. FLIM has advantages over confocal microscopy because it provides an additional dimension with which to discriminate changes in intensity due to the number of proteins from intensity changes due to solvatochromic effects resulting from conformational changes or interactions with other proteins. To our knowledge, the work described here is the first example of the direct use of a fluorescent ncAA in FLIM (the lone previous example used FRET with GFP).^69^ The long lifetime of Acd provides a large dynamic range for FLIM measurements, making it superior to GFP variants or the other genetically encodable ncAAs. This, in addition to Acd’s high quantum yield and photostability, make *Mb* AcdRS 82 a valuable new tool for the protein imaging community, enabling facile labelling of proteins in mammalian cells with a small, non-perturbing fluorophore that has unique properties.

## Experimental

### Aminoacyl tRNA synthetase (RS) screening

Acd RS selection was performed on a library of *Mb* PylRS mutants a life/death selection as previously described.^70^ Briefly, RS library members in a pBK screening plasmid were transformed into *E. coli* cells with a positive selection plasmid, pREP-pylT, containing an amber codon-disrupted chloramphenicol acetyltransferase gene and pylRS tRNA (pylT). Cells containing the pBK-RS library and pREP-pylT were cultured in the presence of chloramphenicol and Acd. Surviving pBK RS library members were isolated and the pBK plasmid was transformed into cells containing a negative selection plasmid, pYOBB2-pylT, encoding an amber codon-disrupted barnase gene under control of an arabinose promoter as well as pylT. Cells containing pBK-RS library and pYOBB2-pylT were cultured in the presence of arabinose, but in the absence of Acd. The remaining pBK-RS library from the surviving colonies was transformed into cells containing a GFP reporter plasmid, pALS-GFP-TAG_150_. Cells containing pBK-RS library and pALS-GFP-TAG_150_ plasmids were grown in 96 well plates in autoinducing media (see ESI) containing 1 mM Acd or Npf, or autoinducing media with no ncAA. GFP fluorescence was measured after 48 h. Additional selection analysis and sequence information are provided in ESI.

### Computational modeling of Acd RS 82 active site

*M. mazei* homologues of Acd RS 82 and 41 were identified by ClustalOmega alignment of *Mb*PylRS and *Mm*PylRS. A crystal structure of *Mm*PylRS bound to *N*_ε_-Boc-L-Lysine and an adenosine triphosphate (ATP) analogue (PDB 2ZIN) was used as the basis for Rosetta modeling.^34^ Firstly, the ATP analogue was removed, and the amino acid ligand was replaced with Ala. Subsequently, missing loops were modeled using Rosetta Remodel and the structure was relaxed using the beta_nov16 scorefunction.^71^ Upon active site mutations, Acd was initially positioned using DARC and several relaxations of the structure were performed.^72^ The final structure was analyzed in PyMol.^73^

### Protein expression and characterization of Acd incorporation

Plasmids for each RS (*Mj* AcdRS A9, *Mb* AcdRS 41, or *Mb* AcdRS 82) in the previously described pDULE2 vector,^35^ containing tRNA_CUA_, were separately transformed into BL21 cells that also contained the pTXB1_αS-TAG_114__Mxe-H_6_ plasmid. After growth in auto-inducing media (see ESI for details), cells were harvested by centrifugation. The supernatant was discarded and the pellet was resuspended, sonicated and pelleted, then the supernatant was collected and the protein purified with a Ni^2+^-NTA column. After elution, the protein was subjected to intein cleavage. Protein production and cleavage yields were analyzed by SDS PAGE. Each construct was further purified by fast protein liquid chromatography and analyzed by MALDI MS. The protein concentrations were analyzed by DC assay and UV-Vis absorbance. See ESI for CaM expression and purification details.

### Mammalian cell expression (EGFP screening)

*Mb* AcdRS 41 and *Mb* AcdRS 82 were separately cloned into the pAcBac1 tR4-MbPyl vector to form pAcBac1 tR4-MbAcdRS41 and pAcBac1 tR4-MbAcdRS82.^54^ These plasmids were transfected into HEK293T/17 cells using Lipofectamine 2000, along with DN-eRF1 (peRF1-E55D.pcDNA5-FRT) and pEGFP-Y40TAG-C1 to evaluate suppression efficiency. Transfections were also performed in the absence of pAcBac1 tR4-MbAcdRS41 or pAcBac1 tR4-MbAcdRS82 as negative (-RS) controls. Transfections with DN-eRF1 and pEGFP-C1 (WT EGFP) were used as positive controls. Transfections were performed with and without DN-eRF1 as well as with varying Acd concentrations, incubation times, and Lipofectamine 2000/DNA ratios to identify optimal expression conditions. Transfections were evaluated by imaging using an Olympus CKX53 microscope and by measuring the EGFP fluorescence of cell lysates using a Tecan M1000 plate reader.

### Mammalian cell expression (confocal microscopy)

pAcBac1-tR4-MbAcdRS82 and DN-eRF1 plasmids were co-transfected into HEK293T/17 cells along with either pIR-TAG676-GFP, pcDNA3.1HE-spHCN-TAG355-YFP, or pcDNA3-FLAG-MBP1-E322TAG (FLAG-MBP1-E322TAG-W340W.pcDNA3-k) plasmids. Confocal imaging .was performed on a Zeiss LSM 710 microscope in and analysed using Fiji software.^74^

### Fluorescence lifetime imaging microscopy (FLIM)

FLIM experiments were performed at room temperature in DMEM supplemented with 10% FBS, penicillin, and streptomycin using an ISS Q2 laser scanner and ISS A320 FastFLIM system mounted on a Nikon TE2000U microscope. Acd was excited using a 375 nm laser and GFP and YFP were excited using bandpass filters to select lines of a YSL SC-PRO 7 supercontinuum laser. FLIM data were acquired and analysed using ISS VistaVision software.

### Plasmid and Acd Availability

Sequences of the pDULE2-MbAcdRS82, for expression in *E. coli*, and pAcBac1-tR4-MbAcdRS82, for expression in mammalian cells, are given in the ESI. The plasmids will be made available via Addgene. Acd is available upon request to Prof. Petersson and is commercially available from Watanabe Chemicals.

## Acknowledgements

This work was supported by funding from the National Science Foundation (NSF CHE-1150351 to E.J.P, MCB-1518265 to R.A.M.) and the National Institutes of Health (NIH R01-NS103873 to E.J.P.; R01-GM131168 to R.A.M.; R01-GM125351 and RO1-EY017564 to S.E.G.; R01-EY010329 and R01-GM127325 to W.N.Z.). Instruments supported by the NSF and NIH include: MALDI MS (NSF MRI-0820996) ESI MS (NIH S10-RR025628), FLIM microscope upgrade (NIH R01-GM125351-S1), and confocal microscope (NIH S10-RR025429). C.M.J. thanks the NIH for funding through the Structural Biology and Molecular Biophysics Training Program (T32 GM-008275). S.E.G. and W.N.Z. thank Edward Lemke for pIR-tAG-GFP cDNA, Peter Schultz for pEGFP-Y40TAG cDNA, and Jason Chin for peRF1-E55D.pcDNA5-FRT cDNA as well as Nathaniel Peters for assistance with confocal imaging. C321.ΔA.exp was a gift from George Church (Addgene plasmid # 49018). We thank Abhishek Chatterjee for providing the pUltra-MbPyl-AcKRS and pET22b-T5 sfGFP plasmids. We thank Jeffrey Morré at the Mass Spectrometry Center at Oregon State University (OSUMSC) for ESI MS assistance.

