## Supplementary Information for "Genetic Encoding of a Highly Photostable, Long Lifetime Fluorescent Amino Acid for Imaging in Mammalian Cells"

*Electronic Supporting Information for*

**Contents:**

*General Information* ........ **S2**

*Fluorescent Amino Acid Properties* ........ **S4**

*Screening of Aminoacyl tRNA Synthetases (RSs) in E. coli* ........ **S14**

*Computational Modeling* ........ **S16**

*Expression of Proteins in E. coli* ........ **S19**

*Comparison of Protein Expression and Imaging in HEK Cells* ........ **S32**

*References* ........ **S49**

*General Information*

**Materials.** *E. coli* BL21 DE3 cells were purchased from Agilent (Santa Clara, CA, USA). Amicon Ultra centrifugal filter units (3 kDa MWCO) were purchased from EMD Millipore. The non-canonical amino acids (ncAAs) acridon-2-ylalanine (Acd), hydroxycoumarin ethylglycine (Hco), and dansylalanine (DanAla) were synthesiszed using previously reported methods.[1-3] 3-[(6-acetyl-2-naphthalenyl)amino]-L-alanine (Anap) was purchased from Cayman Chemical (Ann Arbor, MI, USA). *N*6-((2-(7-hydroxycoumarin-4-yl)ethoxy)carbonyl)-L-lysine (CouLys) was not tested, but shares the 7-hydroxycoumarin chromophore with Hco and is expected to have identical photophyiscal properties.[4] DNA sequencing was performed at the University of Pennsylvania sequencing facility unless otherwise noted. The DC protein assay kit was purchased from BioRad (Hercules, CA, USA). Primers were purchased from Integrated DNA Technologies (Coralville, IA, USA). dNTPs and Agarose were purchased from Invitrogen (ThermoFisher, Waltham, MA, USA). Q5 High-Fidelity DNA polymerase, HiFi DNA Assembly Master Mix, competent High Efficiency E. coli cells (NEB 10-beta), Monarch Gel Extraction Kit were purchased from New England Biolabs (Ipswich, MA, USA). Gel Green DNA gel stain was purchased from Biotium (Freemont, CA, USA). DNA Clean & Concentrator Kit and Plasmid Miniprep Kit were purchased from Zymo Research (Irvine, CA, USA).

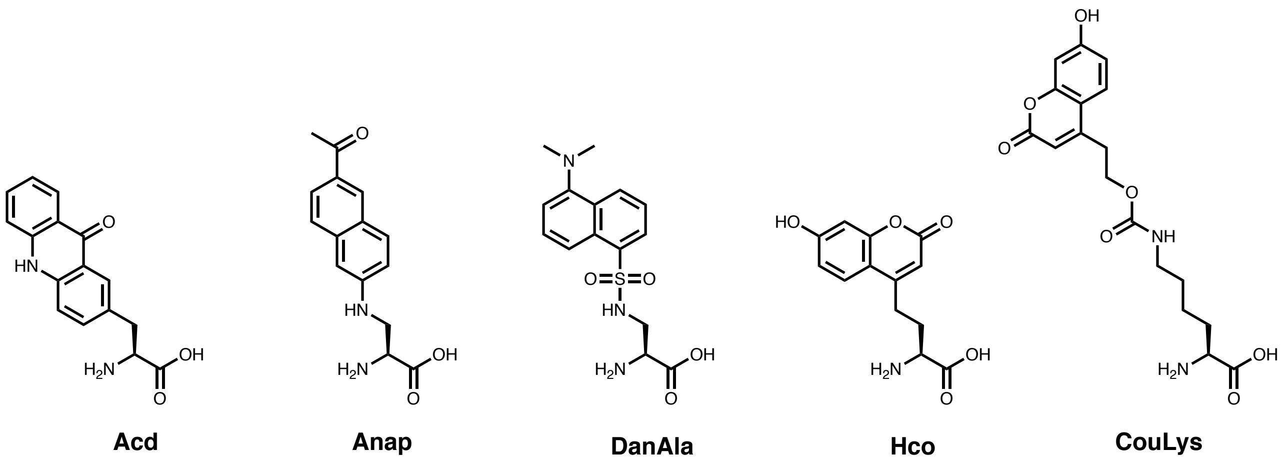

**Instruments.** Matrix assisted laser desorption ionization (MALDI) mass spectrometry (MS) data were acquired on a Bruker Ultraflex III instrument. Electrospray ionization mass spectra (LRMS) were obtained on a Waters Acquity Ultra Performance liquid chromatography (LC) connected to a single quadrupole detector (SQD) mass spectrometer (Milford, MA, USA). Protein quantification was performed on a ThermoScientific Genesys 150 UV-Vis spectrometer (Waltham, MA, USA). Absorbance readings for the DC assay were made on a Tecan M1000 plate reader (Mannedorf, Switzerland). Photostability degradation was performed with a Penn PhD Photoreactor M2 (Pennsburg, PA, USA). Acd fluorescence was visualized by illuminating the gels in the dark with an AnalytikJena 6 Watt lamp, exciting at 365 nm (Beverly, MA, USA). Protein fluorescence lifetime measurements were made using a Photon Technology International (PTI) QuantaMaster™ 40 fluorescence spectrometer (Birmingham, NJ, USA). Quantum yield (QY) measurements were performed using a Jasco FP-8300 Fluorimeter with ILF-835 integrating sphere attachment (Easton, MD, USA). Fluorescence measurements for sfGFP expressions in *E. coli* were performed on BioTek Synergy 2 Microplate Reader with Gen5 All-in-one Microplate Reader Software v.2.00 (Winooski, VT, USA). HEK cell expression tests were imaged on an Olympus (Waltham, MA, USA) CKX53 microscope and lysate measurements were made on a Tecan M1000 plate reader (Mannedorf, Switzerland). HEK cell expression for analysis of Acd incorporation fidelity was imaged using a Keyence BZ-X710 microscope (Itasca, IL, USA). Electrospray ionization (ESI) MS was performed with a Waters Synapt G2 instrument. Confocal imaging was performed on a Zeiss LSM 710 microscope (White Plains, NY, USA). FLIM experiments were performed using an ISS Q2 laser scanner and ISS A320 FastFLIM system (Champaign, IL, USA) mounted on a Nikon TE2000U microscope (Melville, NY, USA).

*Fluorescent Amino Acid Properties*

**Photostability measurements.** Each ncAA was diluted to an absorbance of 0.05 absorbance units (AUs) at 365 nm in MilliQ water. 3 mL of this solution was irradiated in a Penn PhD Photoreactor M2 using a 365 nm light source set to 2% of its maximal intensity (1.5 W of delivered flux at 100% intensity). From this irradiated solution, 105 µL was taken out and kept in a light tight container at the following timepoints: 0, 2, 4, 8, 12, 15, 20, 30, 45, 60, 90 and 120 minutes. At the end of the irradiation, the fluorescence was measured exciting at each ncAA's excitation maximum ($\lambda_{ex}$) and recording at the emission maximum ($\lambda_{ex}$) using a Tecan M1000 plate reader (Table S1). Readings were normalized to their 0 minute fluorescence values. All photobleaching experiments were performed in triplicate and reported in Fig. S1 as an average. The half-life ($t_{1/2}$) for each ncAA was calculated using the exponential model in GraphPad Prism 8. DanAla fluorescence was so low in MilliQ water, that 30 µL of sample was diluted in 80 µL of DMSO before reading.

**Table S1.** Measured photostabilities for various fluorescent ncAAs.

| | $t_{1/2}$ (min) | $\lambda_{ex}$ (nm) | $\lambda_{em}$ (nm) |
|---|---|---|---|
| Acd | 124 (90-197)* | 386 | 420 |
| Anap | 14 (12-15)* | 350 | 495 |
| Hco | 1.7 (1.6-1.8)* | 340 | 460 |
| DanAla | 44 (36-55)* | 340 | 550 |

*95% confidence interval given in parentheses

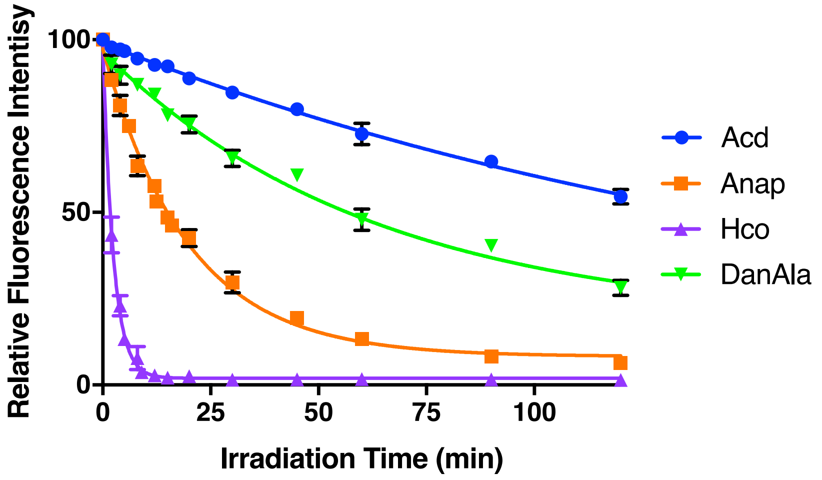


**Fig. S1.** Photostability for each fluorescent ncAA. Measurements performed over three replicates and fits to exponential decays are shown as a solid line.

**Quantum Yield (QY) Measurements.** For each QY replicate, the incident excitation light spectrum was collected in the presence of 2 mL of solvent. After measuring the incident light intensity, 5, 10 or 15 μL of a 5 mM dye stock was added to the solvent and the new fluorescence spectrum (fluorescence intensity and new incident light intensity) was collected. Using the JASCO Quantum Yield Software, the dye quantum yield was calculated by dividing the dye fluorescence intensity by the difference in incident light intensity in the presence and absence of dye. The minimum excitation wavelength for a full excitation incident spectrum using this setup is 363 nm. In order to separate the excitation and emission curves, some dyes were not excited at their maximum excitation wavelength.

**Table S2.** Calculated QYs in varying solvents averaged three replicates.*

| | THF | MeOH | EtOH | DMSO | Water | Buffer |
|---|---|---|---|---|---|---|
| Acd | 0.15 | 0.61 | 0.60 | 0.70 | 0.93 | 0.98 |
| Anap | 0.46 | 0.73 | 0.88 | 0.96 | 0.45 | 0.45 |
| Hco** | n/a | 0.85 | 0.89 | 0.57 | 0.98 | 0.97 |
| DanAla | 0.62 | 0.41 | 0.37 | 0.69 | 0.10 | 0.08 |

* All errors for QY measurements were less than 0.01.

**Due to restictions with the excitation wavelength of our system, NaOH was added to deprotonate the phenolic oxygen of Hco and data could only be recorded for solvents in which the anionic form is soluble.

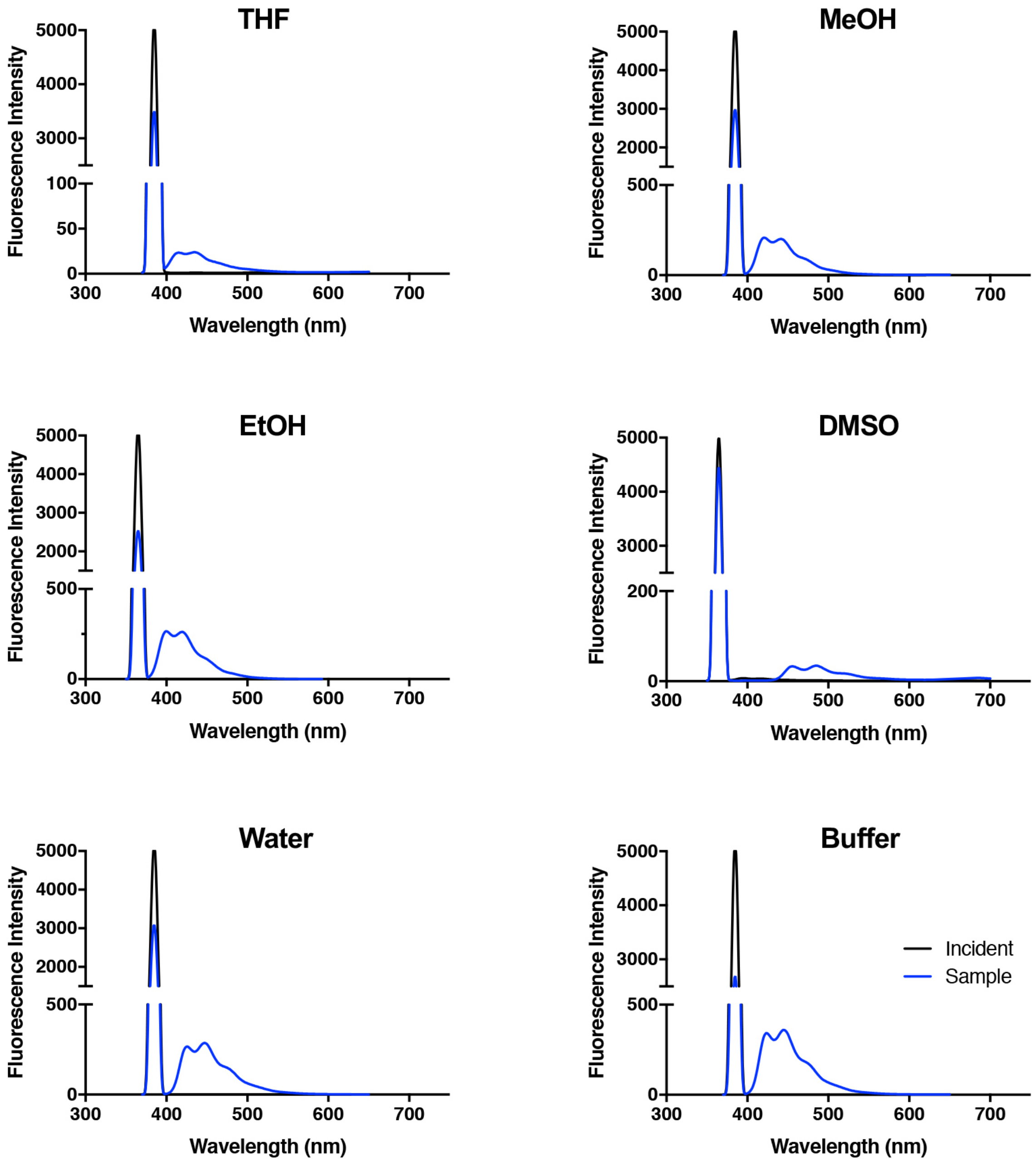


**Fig. S2.** Representitive QY acquisitions for Acd in varing solvent conditions. All had consistent excitation (385 nm, sw 10) and spectral colection (370-650 nm) parameters.

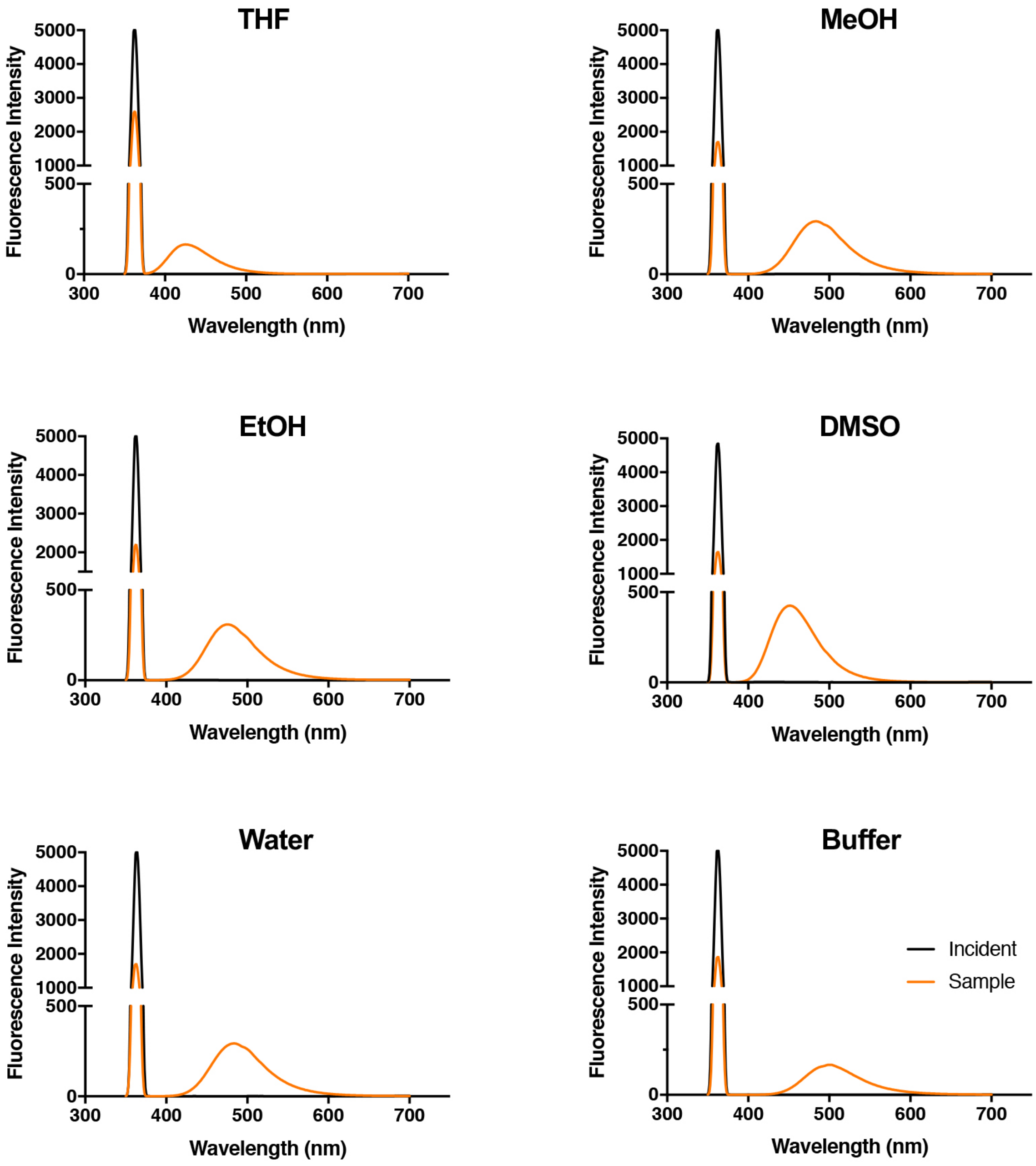


**Fig. S3.** Representitive QY acquisitions for Anap in varing solvent conditions. All had consistent excitation (363 nm, sw 10) and spectral colection (350-600 nm) parameters.

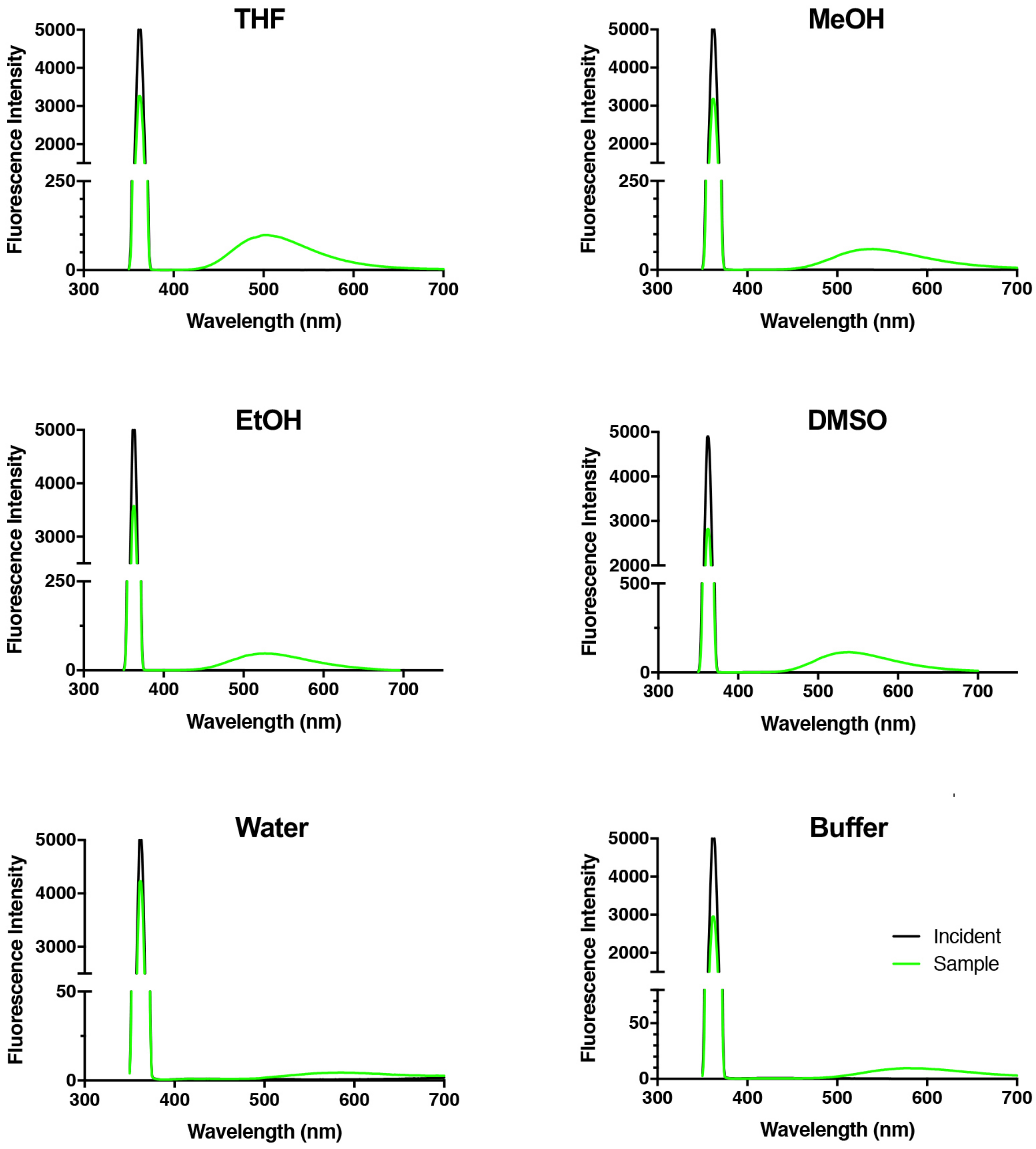


**Figure S4:** Representitive QY acquisitions for DanAla in varing solvent conditions. All had consistent excitation (363 nm, sw 10) and spectral colection (350-800 nm) parameters.

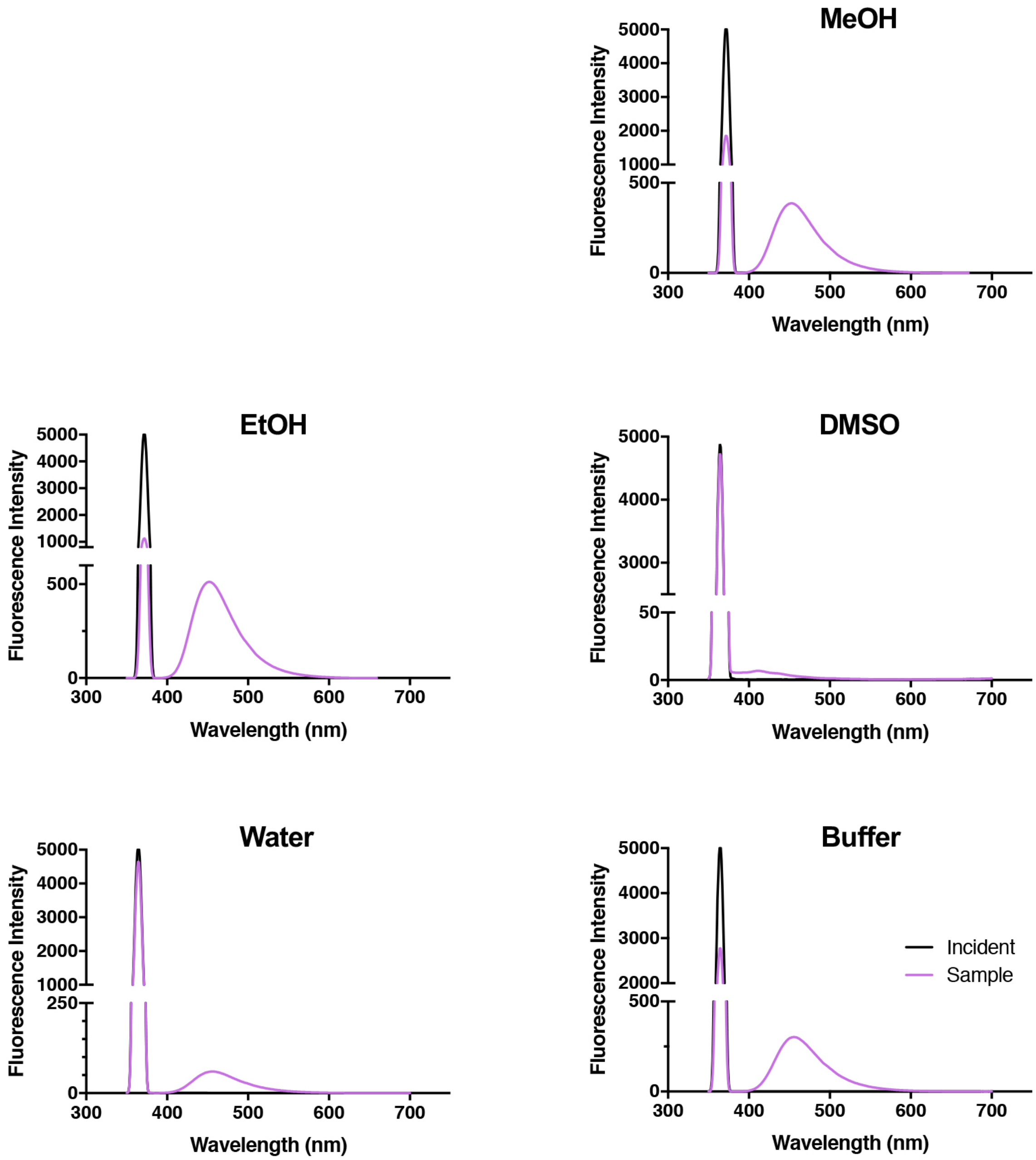


**Fig. S5.** Representitive QY acquisitions for Hco in varing solvent conditions. All had consistent excitation (363 nm, sw 10) and spectral colection (350-800 nm) parameters.

**Fluorescence lifetime measurements.** Time correlated single photon counting (TCSPC) measurements of fluorescence lifetime decays for 5 μM samples of Acd, Anap, DanAla, or Acd were collected with the PTI Quantamaster™ 40 using a pulsed LED with a maximum emission at 340 nm. Fluorescence emission was collected at the indicated wavelength for each ncAA with 20 nm slit widths. The instrument response function (IRF) was collected under identical conditions. Data analysis was performed with FluoFit software (PicoQuant GmbH; Berlin, Germany) using an exponential decay model.

**Table S3.** Acd fluorescence lifetime measured in various solvents and fit $\chi^2$ values.

| Solvent | Lifetime (ns) | $\chi^2$ |
|---|---|---|
| Buffer | 13.52 ± 0.02 | 1.003 |
| Water | 14.04 ± 0.02 | 1.642 |
| MeOH | 8.92 ± 0.03 | 1.144 |
| DMSO | 7.57 ± 0.22 | 1.058 |
| THF | 3.98 ± 0.01 | 1.240 |

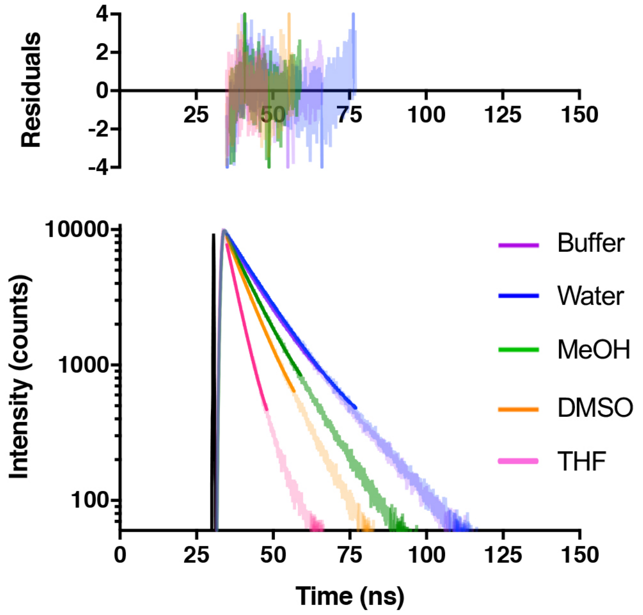


**Fig. S6.** Acd fluorescence lifetime measured in various solvents. Lifetime measurements with exponential fits shown as solid lines (bottom) and residuals after fitting (top). Acd TCSPC data were collected at 420 nm emission.

**Table S4.** Anap fluorescence lifetime measured in various solvents and fit $\chi^2$ values.

| Solvent | Lifetime (ns) | $\chi^2$ |
|---|---|---|
| Buffer | 2.27 ± 0.01 | 1.942 |
| Water | 2.25 ± 0.01 | 1.859 |
| MeOH | 2.62 ± 0.02 | 1.424 |
| DMSO | 2.39 ± 0.01 | 1.919 |
| THF | 2.63 ± 0.01 | 1.878 |

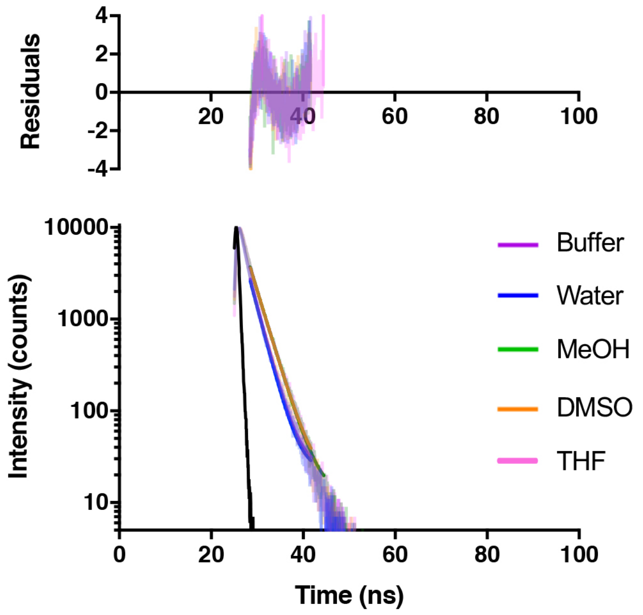


**Fig. S7.** Anap fluorescence lifetime measured in various solvents. Lifetime measurements with exponential fits shown as solid lines (bottom) and residuals after fitting (top). Anap TCSPC data were collected at 495 nm emission.

**Table S5.** DanAla fluorescence lifetime measured in various solvents and fit $\chi^2$ values.

| Solvent | Lifetime (ns) | $\chi^2$ |
|---|---|---|
| Buffer | 3.32 ± 0.01 | 0.990 |
| Water | 3.30 ± 0.01 | 1.088 |
| MeOH | 10.33 ± 0.02 | 1.648 |
| DMSO | 12.16 ± 0.03 | 1.562 |
| THF | 10.16 ± 0.06 | 1.074 |

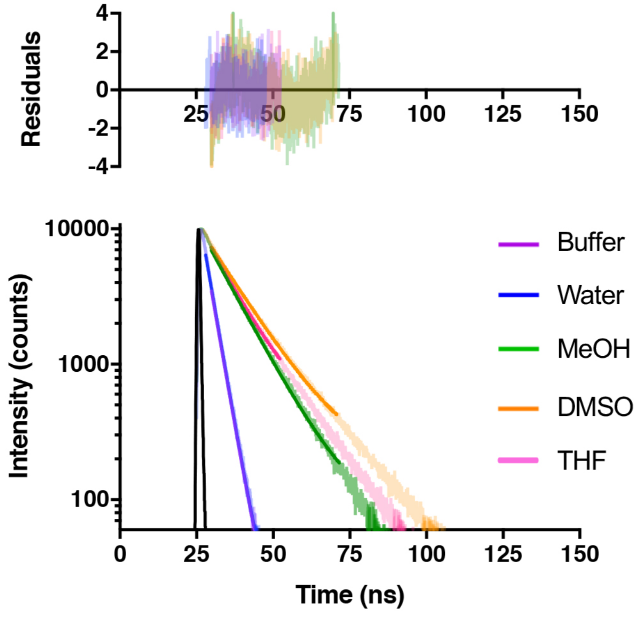


**Fig. S8.** DanAla fluorescence lifetime measured in various solvents. Lifetime measurements with exponential fits shown as solid lines (bottom) and residuals after fitting (top). DanAla TCSPC data were collected at 540 nm emission.

**Table S6.** Hco fluorescence lifetime measured in various solvents and fit $\chi^2$ values.

| Solvent | Lifetime (ns) | $\chi^2$ |
|---|---|---|
| Buffer | 1.68 ± 0.01 | 1.043 |
| Water | 1.64 ± 0.01 | 1.279 |
| MeOH | 4.74 ± 0.02 | 1.01 |
| DMSO | 1.28 ± 0.01 | 1.511 |
| THF | 2.05 ± 0.02 | 1.566 |

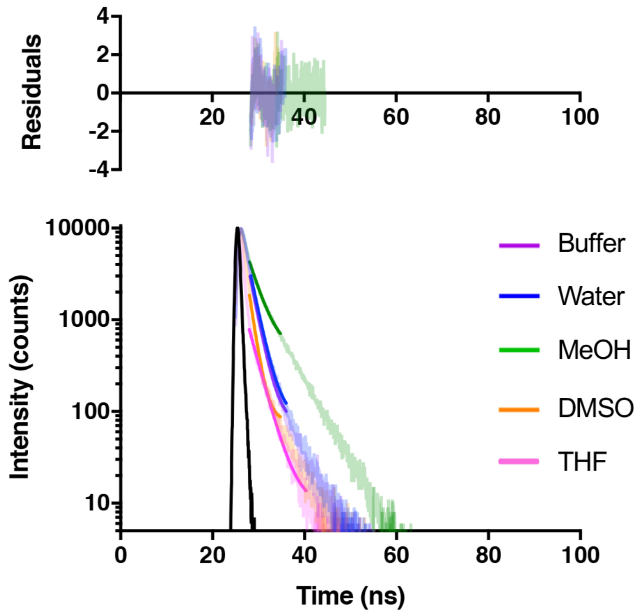


**Fig. S9.** Hco fluorescence lifetime measured in various solvents. Lifetime measurements with exponential fits shown as solid lines (bottom) and residuals after fitting (top). Hco TCSPC data were collected at 455 nm emission.

*Screening of Aminoacyl tRNA Synthetases (RSs) in E. coli*

**Acd aaRS Library Generation and Selection.** To identify a *Methanosarcina barkeri* pyrrolysyl (*Mb* Pyl) RS and $tRNA_{CUA}$ pair capable of site-specifically incorporating Acd into proteins in response to an amber stop codon (TAG) we screened a library of *Mb* PylRS variants in which 5 active-site residues (N311, C313, V366, W382, G387) were randomized to all 20 amino acids and screened using a standard life/death selection.[5-7] Following selection, 96 colonies were assessed for their ability to suppress a TAG codon interrupted superfolder green fluorescent protein gene (sfGFP-$TAG_{150}$) in the presence or absence of Acd as well as in the presence of *N*-phenyl-*p*-amino phenylalanine (Npf). The remaining pBK plasmid library was transformed into 100 µL of pALS-containing DH10B cells. The cells were rescued for 1 hour in 1 mL of SOC (37 °C, 250 rpm) then plated on autoinducing agar plates[7, 8] with 25 µg/mL kanamycin and 50 µg/mL tetracycline. The plates were further divided by the presence or absence of amino acid (1 mM Acd). Plates were grown at 37 °C for 24 hours and then grown on the bench top, at room temperature, for an additional 24 hours. Green colonies were used to inoculate a 96-well plate containing 0.5 mL per well non-inducing media (NIM)[8] containing 50 µg/mL kanamycin and 25 µg/mL tetracycline. After 24 hours of growth (37 °C, 250 rpm), 50 µL of these non-inducing samples were used to inoculate three 96-well plates with 0.5 mL autoinducing media (AIM)[8] containing 50 µg/mL kanamycin, 25 µg/mL tetracycline. One 96-well plate was created with no ncAA, one plate with 1 mM Acd, and one with 1 mM Npf. Fluorescence measurements of the cultures were collected 48 hours after inoculation using a BIOTEK® Synergy 2 Microplate Reader. The emission from 528 nm (20 nm bandwidth) was summed with excitation at 485 nm (20 nm bandwidth). Samples were prepared by diluting suspended cells directly from culture 4-fold with sterile water. Data after 48 hours of incubation are shown in Fig. S10 for the colonies with the highest fluorescence in the presence of Acd. The top 25 performing clones were sequenced and 13 unique RS clones were identified (Table S7).

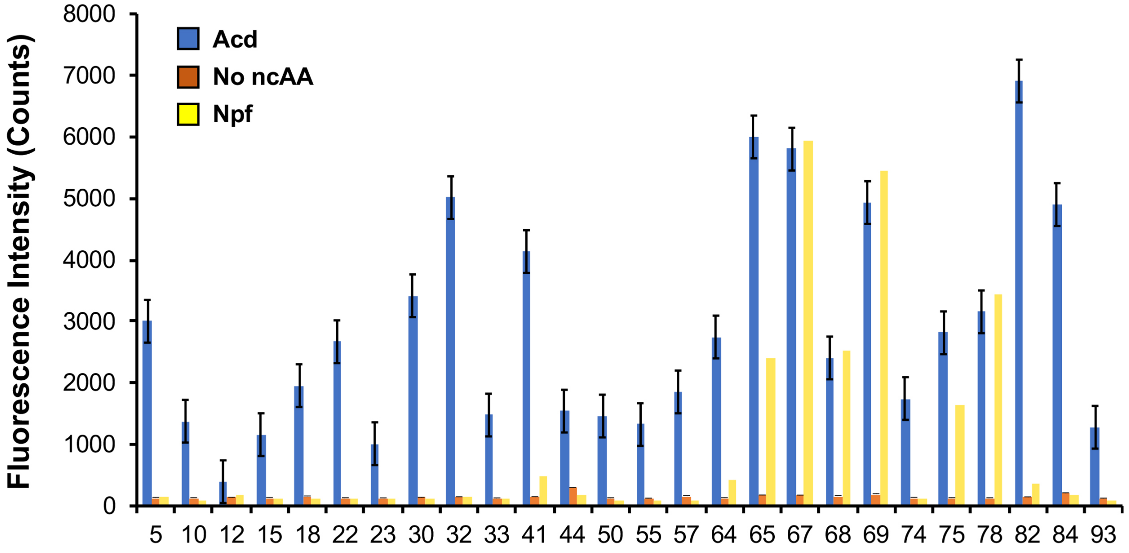


**Fig. S10**. Fluorescence measurements of RSs with sfGFP reporter after 48 hours. Orange, blue, and yellow represent fluorescence from colonies induced in media containing no ncAA, 1 mM Acd**,** or 1 mM Npf, respectively. Data for clones 5, 10, 12, 22, 32, 41, 50, 57, 64, 67, 68, 78, 82 are repeated from main text Fig. 2 to enable comparison to all clones for which sequencing is shown in Table S7.

**Table S7.** Sequences of PylRS mutants selected for highest GFP fluorescence in GFP-TAG$_{150}$ assay

| ***Mb* Pyl RS Variant** | **N311** | **C313** | **V366** | **W382** | **G386** | **Other Mutations** | |
|---|---|---|---|---|---|---|---|
| 5 | S | G | C | V | G | D299G | C319S |
| 22 | S | G | A | V | G | G300S | |
| 30, 64 | A | A | A | V | G | | |
| 32 | S | G | A | V | G | | |
| 50, 10, 15, 18, 23, 33, 93 | S | A | A | V | G | K172R | |
| 65, 67, 69 | A | A | C | T | G | | |
| 68 | A | A | T | T | G | | |
| 57, 74, 84 | S | A | A | V | G | | |
| 82 | S | G | A | T | G | L155V | |
| 12 | S | A | A | T | G | | |
| 41, 44 | S | C | A | T | G | | |
| 75 | S | G | A | T | G | | |
| 78 | A | G | A | T | G | K251R | K286R |

*Computational Modeling*

**Preparation of starting PylRS structure.** PDB 2ZIN was obtained from the PDB. The Boc-Lys ligand was manually changed to Ala in PyMol using the Mutagenesis Wizard. The structure was prepared for Rosetta modeling. To model missing loops into the prepared structure, a blueprint file was generated. The blueprint was modified such the rotamer, identity, and secondary of all residues solved in the structure would remain fixed. The identity of missing residues was defined and the secondary structure of them and those residues immediately up and downstream was defined as loop. After finalizing the blueprint file, a RosettaScripts xml file was written containing the RemodelMover, which was then used to build in missing segments as follows. The structure was then renumbered for further modeling. Finally, this structure was relaxed into the beta_nov16 scorefunction. The resulting structure was then used for modeling of enzyme mutations.

**Modeling of PylRS variants 41 and 82**. PylRS variants were modeled in PyRosetta4 (Mac version 203 for Python 3.6). A script was written which first mutated all relevant residues to their mutant identities. The pose was then FastRelaxed with coordinates constrained as follows. The relax_taskfactory included all residues with the IncludeCurrent(), RestrictToRepacking(), and InitializeFromCommandLine() task operations. The relax_movemapfactory allowed all chi angles, backbone, bond angles, and jumps to move and prohibited bond lengths from moving. Structures were then saved to PDB files for both AcdRS 41 and AcdRS 82. Then, the Ala ligand for each was deleted and ligand-free structures were output to PDB files.

**Docking of Acd into PylRS variants 41and 82.** To initially position the Acd ligand in the binding pocket, DARC was used.[9] Ray files were generated for the pocket surrounding the polar mutations at positions 311 and 382 (positions 159 and 230 in the model). Rays were cast using the Ala ligand from the original structure as the origin. A params file and rotamer library for Acd were generated using a previously published method.[10] The params file was edited such that Acd was named LIG and the PROTEIN and L_AA properties were removed from the properties list and DARC was performed. The LIG residue from the pdb output of the enzyme/ligand complex was replaced with ACD and the merged pdb was prepared for further modeling.

The position of Acd in the pocket was optimized in PyRosetta using the beta_nov16 scorefunction as follows. Firstly, the ligand position was adjusted. A FastRelax of the ligand constrained to starting coordinates without ramping-down was performed in DualSpace, allowing chi angles, backbone, bond angles, and jumps to minimize only at the ligand residue (i.e., the enzyme remained unmodified). Next, the ligand and all enzyme residues within 10 Å of it were packed using the PackRotamersMover and the beta_nov16_soft scorefunction. The full structure was then minimized in Cartesian space using the beta_nov16_cart scorefunction and the MinMover. Finally, the full structure was relaxed as above via FastRelax two times sequentially. The first time it was constrained to starting coordinates with ramping down of constraints. The second time all constraints were removed.

Due to steric constriction of the pocket, Acd could not be initially positioned well in the pocket of AcdRS 41 using DARC. Therefore, to model Acd bound to AcdRS 41, all steps of the docking protocol following DARC were performed after mutation to $C_{313}$ in the fully docked AcdRS 82 structure. Continued steric clashes caused a flip of the Acd ring and prevented adopting more optimal hydrogen bonding interactions as in AcdRS 82. The AcdRS 41 structure and comparison of these distances for AcdRS 41 and 82 is shown in Fig. S11.

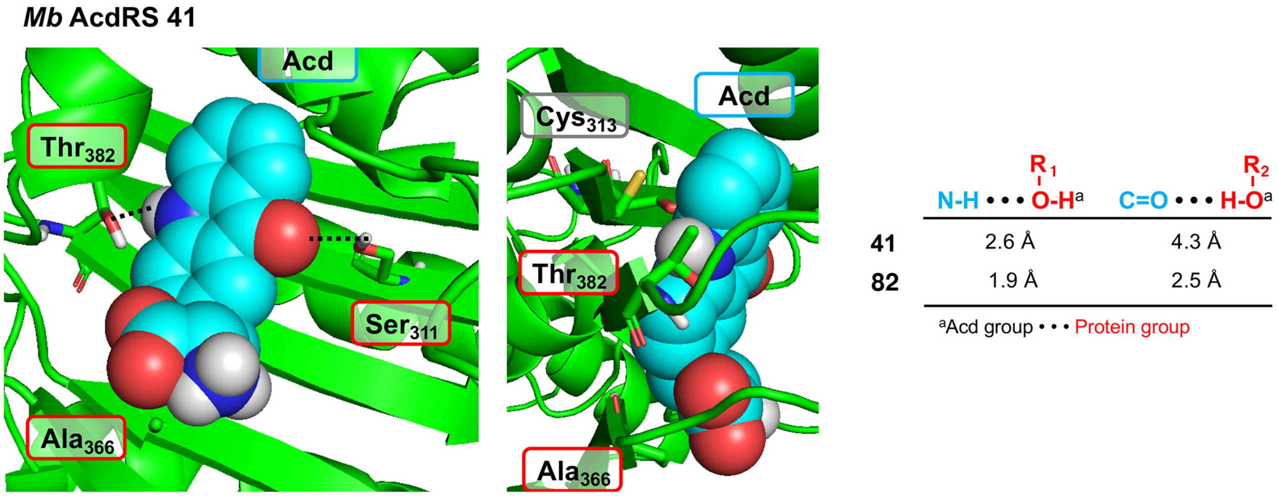


**Fig. S11.** A homology model of *Mb* AcdRS 41 based on the x-ray crystal structure of *Mm* PylRS (PDB ID: 2ZIN)[11] with Acd docked in the active site using Rosetta. Key points of interaction with Acd are shown from two angles. Sites of mutation relative to parent *Mb* PylRS are labeled in red boxes, residues from the parent *Mb* PylRS are shown in grey boxes. Steric clashes with $Cys_{313}$ prevent adopting optimal hydrogen bonding interactions with $Thre_{382}$ and Ser $_{311}$. Interatom distances from the AcdRS 41 and 82 structures are shown. Due to a flip of the Acd ring, $R_1$-O-H corresponds to $Thr_{382}$ for AcdRS 41 and to $Ser_{311}$ for AcdRS 82. $R_2$-O-H corresponds to $Ser_{311}$ for AcdRS 41 and to $Thr_{382}$ for AcdRS 82.

```
MmPylRS                    MDKKPLNTLISATGLWMSRTGTIHKIKHHEVSRSKIYIEMACGDHLVVNNSRSSRTARAL  60
MmPylRS_Rosetta_Model      ------------------------------------------------------------  0
MbAcdRS 41                 MDKKPLDVLISATGLWMSRTGTLHKIKHHEVSRSKIYIEMACGDHLVVNNSRSCRTARAF  60
MbAcdRS 82                 MDKKPLDVLISATGLWMSRTGTLHKIKHHEVSRSKIYIEMACGDHLVVNNSRSCRTARAF  60
MbPylRS                    MDKKPLDVLISATGLWMSRTGTLHKIKHHEVSRSKIYIEMACGDHLVVNNSRSCRTARAF  60


MmPylRS                    RHHKYRKTCKRCRVSDEDLNKFLTKANEDQTSVKVKVVSAPTRTKKAMPKSVARAPKPLE 120
MmPylRS_Rosetta_Model      ------------------------------------------------------------  0
MbAcdRS 41                 RHHKYRKTCKRCRVSDEDINNFLTRSTESKNSVKVRVVSAPK-VKKAMPKSVSRAPKPLE 119
MbAcdRS 82                 RHHKYRKTCKRCRVSDEDINNFLTRSTESKNSVKVRVVSAPK-VKKAMPKSVSRAPKPLE 119
MbPylRS                    RHHKYRKTCKRCRVSDEDINNFLTRSTESKNSVKVRVVSAPK-VKKAMPKSVSRAPKPLE 119


MmPylRS                    NTEAAQAQPSGSKFSPAIPVSTQESVSVPASVSTSISSISTGATASALVKGNTNPITSMS 180
MmPylRS_Rosetta_Model      ------------------------------------------------------------  0
MbAcdRS 41                 NSVSAKASTNTSRSVPSP----------------------------------AKSTPNSS 145
MbAcdRS 82                 NSVSAKASTNTSRSVPSP----------------------------------AKSTPNSS 145
MbPylRS                    NSVSAKASTNTSRSVPSP----------------------------------AKSTPNSS 145


MmPylRS                    APVQASAPALTKSQTDRLEVLLNPKDEISLNSGKPFRELESELLSRRKKDLQQIYAEERE 240
MmPylRS_Rosetta_Model      -------PALTKSQTDRLEVLLNPKDEISLNSGKPFRELESELLSRRKKDLQQIYAEERE  53
MbAcdRS 41                 VPASAPAPSLTRSQLDRVEALLSPEDKISLNMAKPFRELEPELVTRRKNDFQRLYTNDRE 205
MbAcdRS 82                 VPASAPAPSVTRSQLDRVEALLSPEDKISLNMAKPFRELEPELVTRRKNDFQRLYTNDRE 205
MbPylRS                    VPASAPAPSLTRSQLDRVEALLSPEDKISLNMAKPFRELEPELVTRRKNDFQRLYTNDRE 205
                                  *::*:** **:*.**.*:*:**** .******* **::***:*:*::*:::**

MmPylRS                    NYLGKLEREITRFFVDRGFLEIKSPILIPLEYIERMGIDNDTELSKQIFRVDKNFCLRPM 300
MmPylRS_Rosetta_Model      NYLGKLEREITRFFVDRGFLEIKSPILIPLEYIERMGIDNDTELSKQIFRVDKNFCLRPM 113
MbAcdRS 41                 DYLGKLERDITKFFVDRGFLEIKSPILIPAEYVERMGINNDTELSKQIFRVDKNLCLRPM 265
MbAcdRS 82                 DYLGKLERDITKFFVDRGFLEIKSPILIPAEYVERMGINNDTELSKQIFRVDKNLCLRPM 265
MbPylRS                    DYLGKLERDITKFFVDRGFLEIKSPILIPAEYVERMGINNDTELSKQIFRVDKNLCLRPM 265
                           :*******:**:***************** **:*****:***************:*****

MmPylRS                    LAPNLYNYLRKLDRALPDPIKIFEIGPCYRKESDGKEHLEEFTMLNFCQMGSGCTRENLE 360
MmPylRS_Rosetta_Model      LAPNLYNYLRKLDRALPDPIKIFEIGPCYRKESDGKEHLEEFTMLNFCQMGSGCTRENLE 173
MbAcdRS 82                 LAPTLYNYLRKLDRILPGPIKIFEVGPCYRKESDGKEHLEEFTMVSFCQMGSGCTRENLE 325
MbAcdRS 82                 LAPTLYNYLRKLDRILPGPIKIFEVGPCYRKESDGKEHLEEFTMVSFGQMGSGCTRENLE 325
MbPylRS                    LAPTLYNYLRKLDRILPGPIKIFEVGPCYRKESDGKEHLEEFTMVNFCQMGSGCTRENLE 325
                           ***.********** **.******:*******************:.* ************

MmPylRS                    SIITDFLNHLGIDFKIVGDSCMVYGDTLDVMHGDLELSSAVVGPIPLDREWGIDKPWIGA 420
MmPylRS_Rosetta_Model      SIITDFLNHLGIDFKIVGDSCMVYGDTLDVMHGDLELSSAVVGPIPLDREWGIDKPWIGA 233
MbAcdRS 41                 ALIKEFLDYLEIDFEIVGDSCMVYGDTLDIMHGDLELSSAAVGPVSLDREWGIDKPTIGA 385
MbAcdRS 82                 ALIKEFLDYLEIDFEIVGDSCMVYGDTLDIMHGDLELSSAAVGPVSLDREWGIDKPTIGA 385
MbPylRS                    ALIKEFLDYLEIDFEIVGDSCMVYGDTLDIMHGDLELSSAVVGPVSLDREWGIDKPWIGA 385
                           ::*.:**::* ***:**************:**********.***: ********** ***

MmPylRS                    GFGLERLLKVKHDFKNIKRAARSESYYNGISTNL 454
MmPylRS_Rosetta_Model      GFGLERLLKVKHDFKNIKRAARSESYYNGISTNL 267
MbAcdRS 41                 GFGLERLLKVMHGFKNIKRASRSESYYNGISTNL 419
MbAcdRS 82                 GFGLERLLKVMHGFKNIKRASRSESYYNGISTNL 419
MbPylRS                    GFGLERLLKVMHGFKNIKRASRSESYYNGISTNL 419
                           ********** *.*******:*************
```

**Fig. S12.** Multiple sequence alignment of PylRS proteins used in model construction. Alignment performed with CLUSTAL O(1.2.4).[12] * = sequence identity, : = high conservation, . = low conservation, yellow highlights indicate site of mutation in AcdRS 41 or 82.

*Expression of Proteins in E. coli*

**Previous Cloning.** The gene containing full length calmodulin (CaM) was previously cloned into the pTXB1 vector containing a C-terminal MxeGyrA intein, followed by a $His_6$ purification tag.[13] The TAG codon for Acd incorporation at position 113 via amber stop codon suppression was previously inserted using QuikChange® PCR, yielding the CaM-$TAG_{113}$-GyrA-$H_6$ plasmid.[14] The construction of the wild-type (WT) CaM-GyrA-$H_6$ plasmid was previously reported in Walters *et al.*[15] Similarly, the gene encoding full length α-synuclein (αS) was previously cloned into a pTXB1 vector containing a C-terminal MxeGyrA intein, followed by a $His_6$ purification tag.[16] The TAG codon used for Acd incorporation was previously inserted at position 114, yielding the pTXB1_αS-$TAG_{114}$_Mxe-$H_6$ plasmid.[17] The construction of the WT αS-GyrA-$H_6$ plasmid was reported in Haney *et al.*[16]

**General information about cloning.** All PCR reactions were carried out on a T100 thermocycler from Bio-Rad (Hercules, CA, USA). DNA concentrations were determined with a TECAN Nanoquant plate on an Infinite M1000Pro plate reader. Agarose gels were visualized on a Typhoon Imager (GE Healthcare, Marlborough, MA, USA) or a SmartBlue transilluminator (Southern Labware, Cumming, GA USA).

**Cloning of aminoacyl tRNA synthetases (RSs) into pDule2.** The following pDule2 forward and reverse primers were used to amplify AcdRS 41 and 82 genes from the pBK plasmid.

pDule2 Forward primer:

`5′-GAGTTTACGCTTTGAGGAATCCCCCATGGATAAAAAACCGCTGGATG-3′`

pDule2 Reverse primer:

`5′-CCTCTTCTGAGATGAGTTTTTGTTCTTACAGGTTCGTGCTAATGC-3′`

The amplified fragment was gel purified and cloned into the pDule2 plasmid using a SLICE reaction as described for other RSs in Jang *et al.*[6]

**Expression of αS-$E_{114}$δ with various AcdRSs.** pDule2 plasmids for each RS (*Mj* AcdRS A9, *Mb* AcdRS 41, or *Mb* AcdRS 82) were separately transformed into BL21 cells that also contained the pTXB1_αS-$TAG_{114}$_Mxe-$H_6$ plasmid. These transformations were plated on an LB-agar plate containing streptomycin (Strep, 50 μg/mL) and amplicillin (Amp, 100 μg/mL). Single colonies were picked and grown in non-inducing media (NIM, 3x5 mL each, containing 50 μg/mL Strep and 100 μg/mL Amp, media prepared as described previously[8]) at 37 °C, shaking at 250 RPM until saturation. 12.5 mL of primary culture was added to a 250 mL secondary culture of auto-inducing media (AIM, media prepared as described previously[8]) containing 1 mM Acd. Secondary cultures were grown at 37 °C with shaking at 250 RPM for four hours at which time the temperature and shaking speed were decreased to 30 °C and 200 RPM. Secondary cultures were left to incubate for 18-22 hours.

**Expression of CaM-$L_{113}\delta$ with various AcdRSs.** Expression of Acd-labeled CaM constructs was performed similarly to αS-$E_{114}\delta$ expression, where the plasmids encoding each RS were separately transformed into BL21 cells that also contained the CaM-$TAG_{112}$-GyrA-$H_6$ plasmid.

**Expression of wild-type (WT) protein constructs**. pTXB1 plasmids encoding WT αS or WT CaM were separately transformed into BL21 cells. These transformations were plated on an LB-agar plate containing Amp (100 μg/mL). Single colonies were picked and grown in NIM (3x5 mL each, containing 100 μg/mL Amp) at 37 °C, shaking at 250 RPM until saturation. 12.5 mL of primary culture was added to a 250 mL secondary culture of AIM. Secondary cultures were grown at 37 °C with shaking at 250 RPM for four hours, at which time the temperature and shaking speed were decreased to 30 °C and 200 RPM. Secondary cultures were left to incubate for 18-22 hours.

**Purification of αS constructs.** Cells were harvested by centrifugation at 4000 RPM using a GS3 rotor and a Sorvall RC-5 centrifuge for 30 minutes at 4 °C. The supernatant was discarded and the pellet was resuspended in 10 mL of lysis buffer (40 mM Tris, 5 mM EDTA, pH 8.0) containing one cOmplete mini EDTA-free protease inhibitor cocktail (Sigma-Aldrich; St. Louis, MO, USA). Resuspended cells were lysed via sonication (30 amps power, 1 second pulse, 2 seconds rest, 3.5 minutes total sonication time. The sonicated resuspension was then pelleted at 14,000 RPM in an SS-34 rotor (Sorvall RC-5 centrifuge) for 35 minutes at 4 °C. The supernatant was collected and incubated with 3 mL of $Ni^{2+}$-NTA resin (Goldbio; St. Louis, MO, USA) for 1 hour with shaking at 4 °C. The slurry was added to a fritted column and the liquid was allowed to flow through. The resin was then washed with 2 x 5 mL of Wash 1 buffer (50 mM HEPES, pH 7.5) and 3 x 5 mL of Wash 2 buffer (50 mM HEPES, 10 mM imidazole, pH 7.5). Constructs were eluted off the resin with 3x5 mL of Elution buffer (50 mM HEPES, 300 mM imidazole, pH 7.5). The eluent was combined and immediately subjected to intein cleavage conditions by adding β-mercaptoethanol (βME) to a final concentration of 200 mM and incubated on a rotisserie at room temperature for 18 hours. The solution was then dialyzed against 20 mM Tris, pH 8.0 buffer overnight. The dialyzed solution was incubated with 3 mL of $Ni^{2+}$-NTA resin (product number and company) for 1 hour with shaking at 4 °C. The slurry was added to a fritted column and the flow-through was collected. As a precaution, any uncleaved intein products were eluted off the resin with 3x5 mL Elution buffer. The flow through containing cleaved products was dialyzed against 20 mM Tris pH, 8.0 overnight. Protein production and cleavage yields were analyzed by SDS-page (10% acrylamide/glycine gel). Each construct was purified by ion-exchange chromatography using a HiTrap Q HP column (5 mL) on an ÄKTA FPLC using a 70 minute NaCl gradient (0 to 500 mM NaCl in 20 mM Tris, pH 8.0). The fractions containing the product were identified by MALDI MS. The constructs were concentrated using Amicon Ultra centrifugal filter units (3 kDa MWCO), and their concentration analyzed by DC assay and UV-Vis absorbance using the molar extinction coefficient for Acd (5,700 $M^{-1}cm^{-1}$ at 386 nm) and for αS based on tyrosine absorbance (5,120 $M^{-1}cm^{-1}$ at 280 nm).

**Purification of CaM constructs.** Cells were harvested, lysed, and protein isolated using an identical procedure to that for αS constructs. Protein production and cleavage yields were analyzed by SDS-page (10% acrylamide/glycine gel). Each construct was purified by ion-exchange chromatography using a HiTrap Q HP column (5 mL) on an ÄKTA FPLC using a 100 minute NaCl gradient (0 to 500 mM NaCl in 20 mM Tris, pH 8.0). The fractions containing the product were identified by MALDI MS. The constructs were concentrated using Amicon Ultra centrifugal filter units (3 kDa MWCO), and their concentration analyzed by DC assay and UV-Vis absorbance (WT CaM = 3006 $M^{-1}cm^{-1}$ at 276 nm). Once aliquoted, the samples were flash frozen in liquid $N_2$ and lyophilized to a powder.

**Table S8.** Protein yields for Acd incorporation in αS*

| | Pure protein (mg) | Protein/1 L culture (mg) |
|---|---|---|
| WT** | 18.08 | 72.3 |
| *Mj* AcdRS A9 | 11.19 | 44.8 |
| *Mb* AcdRS 41 | 1.00 | 4.0 |
| *Mb* AcdRS 82 | 8.12 | 32.5 |

*Based on molar absorptivity and confirmed by DC assay. WT amount scaled to account for some protein lost in second Ni column cleanup procedure based on gel band intensities in Ni Column 2 FT and El lanes in Fig. S13.

**Table S9.** Protein yields for Acd incorporation in CaM*

| | Pure protein (mg) | Protein/1 L culture (mg) |
|---|---|---|
| WT | 5.40 | 21.6 |
| *Mj* AcdRS A9 | 4.08 | 16.3 |
| *Mb* AcdRS 41 | 0.35 | 1.4 |
| *Mb* AcdRS 82 | 1.63 | 6.5 |

*Based on molar absorptivity and confirmed by DC assay.

**Gel Analysis.** Samples were taken during cell lysis, nickel column purification, intein cleavage, and the second nickel column purification. To confirm expression and monitor protein loss, these samples were analyzed by SDS-page (10% acrylamide tris-glycine gel, electrical current ranging from 60-140 volts over 1.5 hours). Before staining or imaging, the gel was fixed in destaining solution (40% (v/v) methanol, 10% (v/v) (v/v) glacial acetic acid in Milli-Q water) to remove free Acd which was observed to be non-covalently bound to proteins in the cell lysates. Acd fluorescence was visualized by illuminating the gels in the dark with an AnalytikJena 6-Watt handheld lamp at 365 nm. The gels were then stained with Coomasie brilliant blue solution (0.25% (w/v) brilliant Blue-G (ThermoFisher), 40% (v/v) methanol, 10% (v/v) glacial acetic acid) and destained by shaking in destaining solution to remove excess dye. A representative gel from expression of each protein construct is shown below.

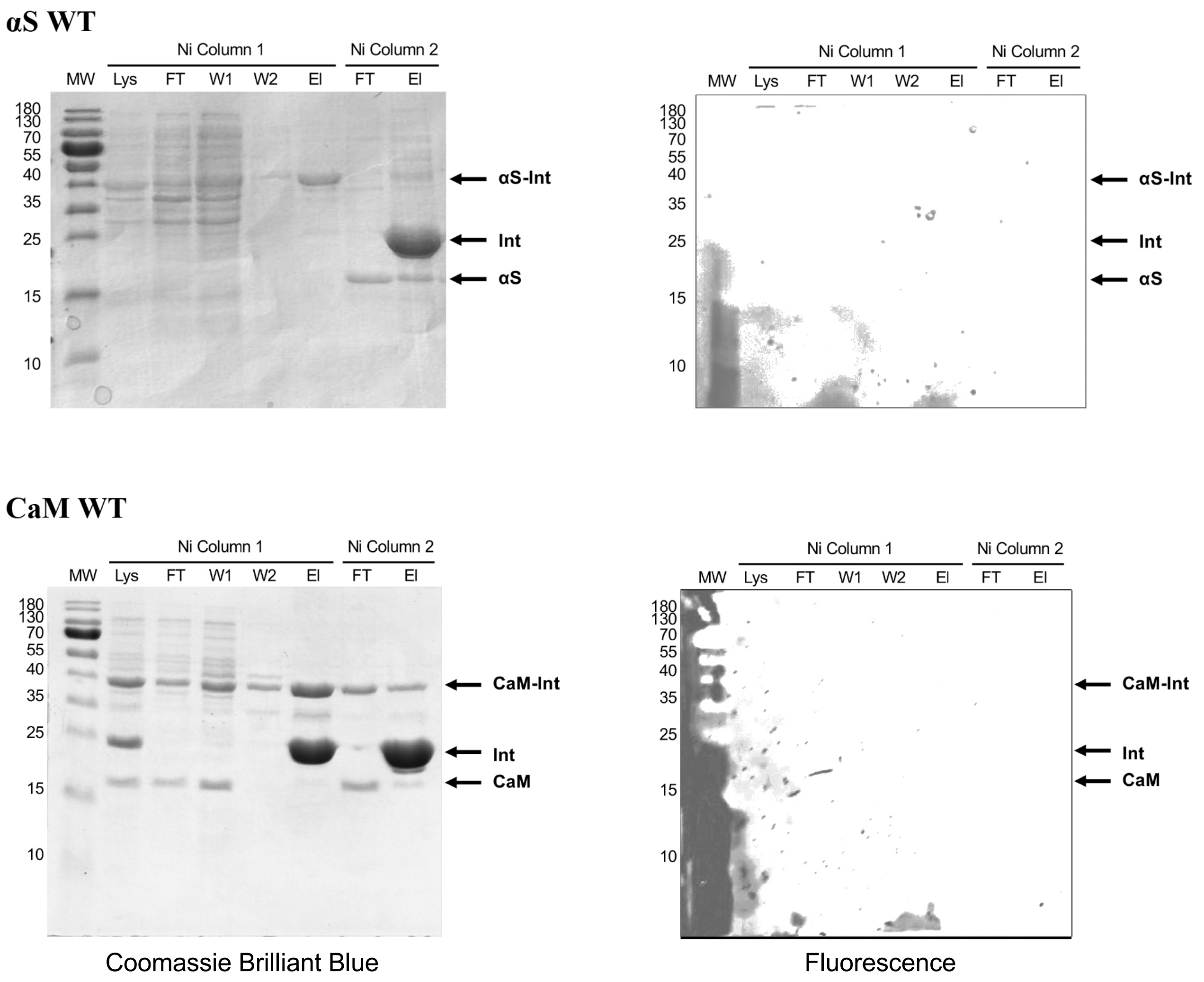


**Fig. S13.** Gel analysis of expression of WT αS and CaM. MW = molecular weight in kDa. Lys = lysate, FT = flow through, W1 = wash 1, W2 = wash 2, El = elution, Int = intein.

**αS-E$_{114}$δ *Mj* AcdRS A9**

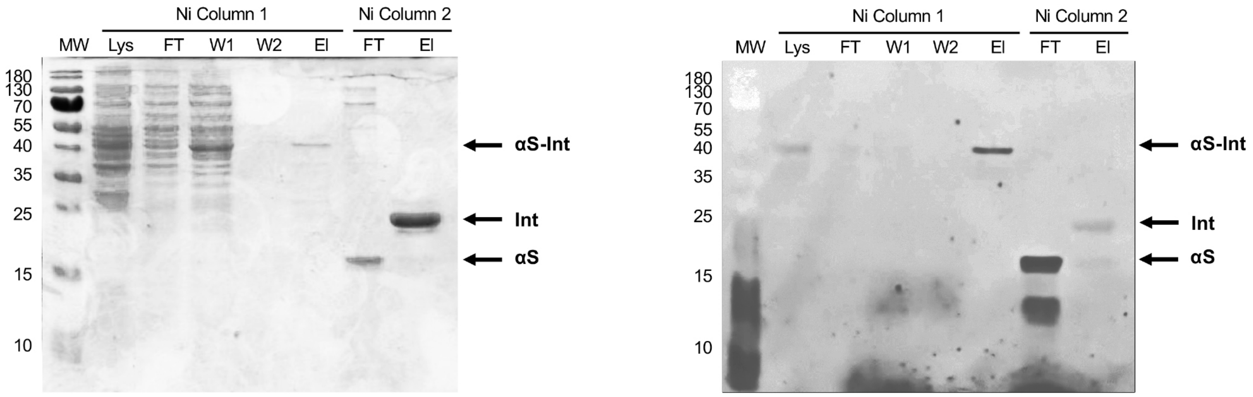


**αS-E$_{114}$δ *Mb* AcdRS 41**

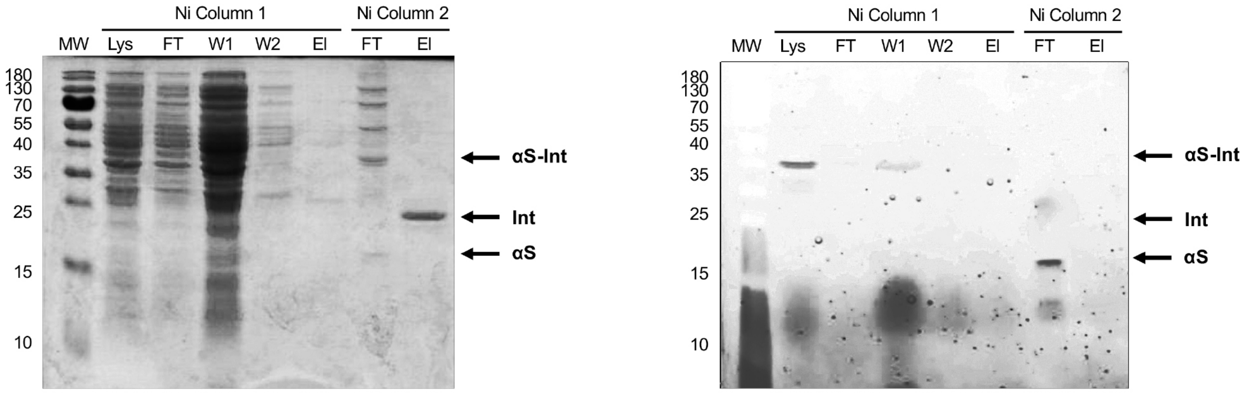


**αS-E$_{114}$δ *Mb* AcdRS 82**

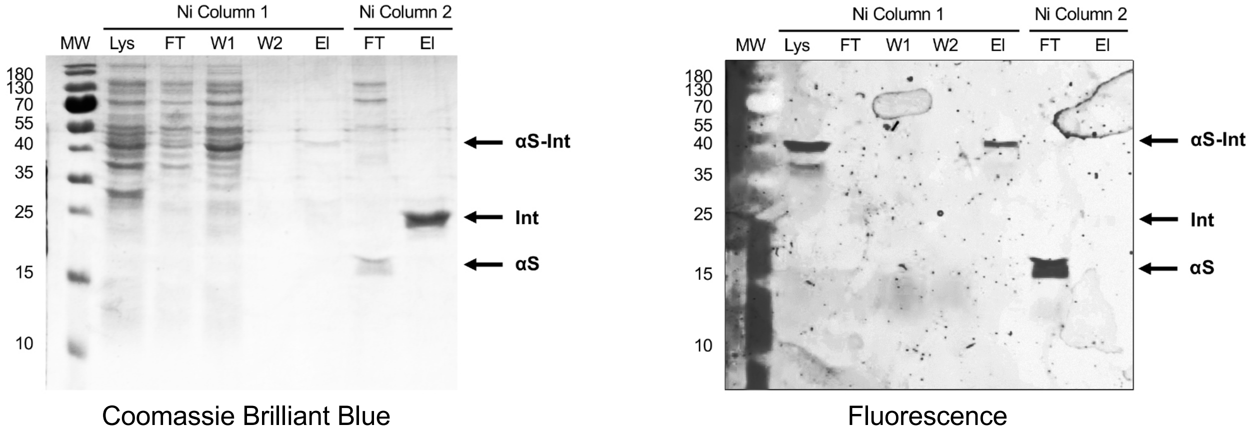


**Fig. S14.** Gel analysis of Acd incorporation in αS using various RSs. MW = molecular weight in kDa. Lys = lysate, FT = flow through, W1 = wash 1, W2 = wash 2, El = elution, Int = intein.

### CaM-L$_{113}$δ *Mj* AcdRS A9

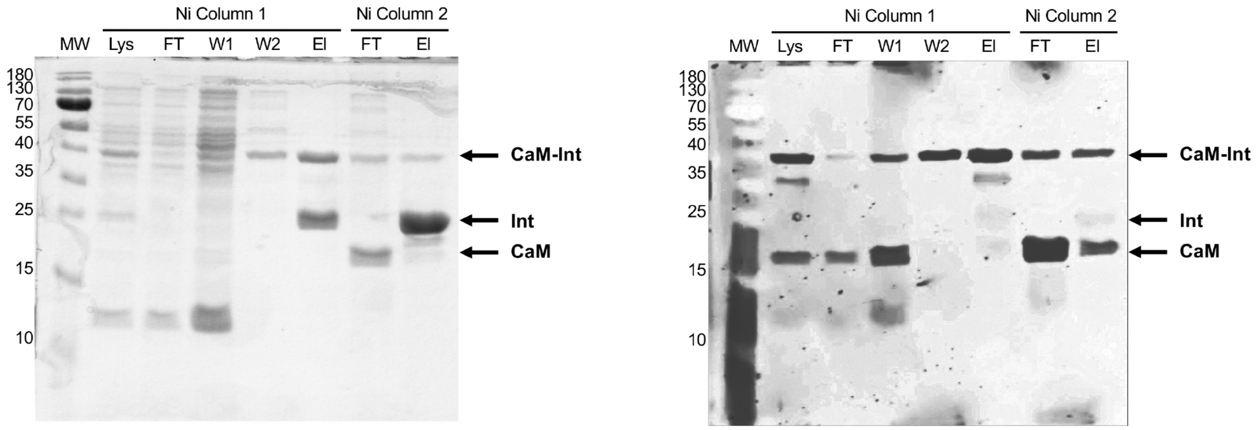


### CaM-L$_{113}$δ *Mb* AcdRS 41

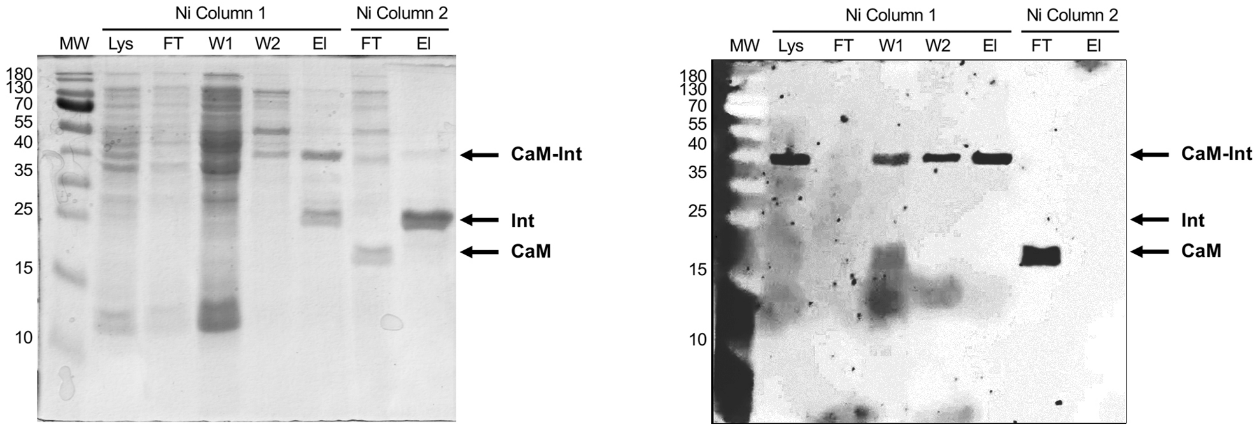


### CaM-L$_{113}$δ *Mb* AcdRS 82

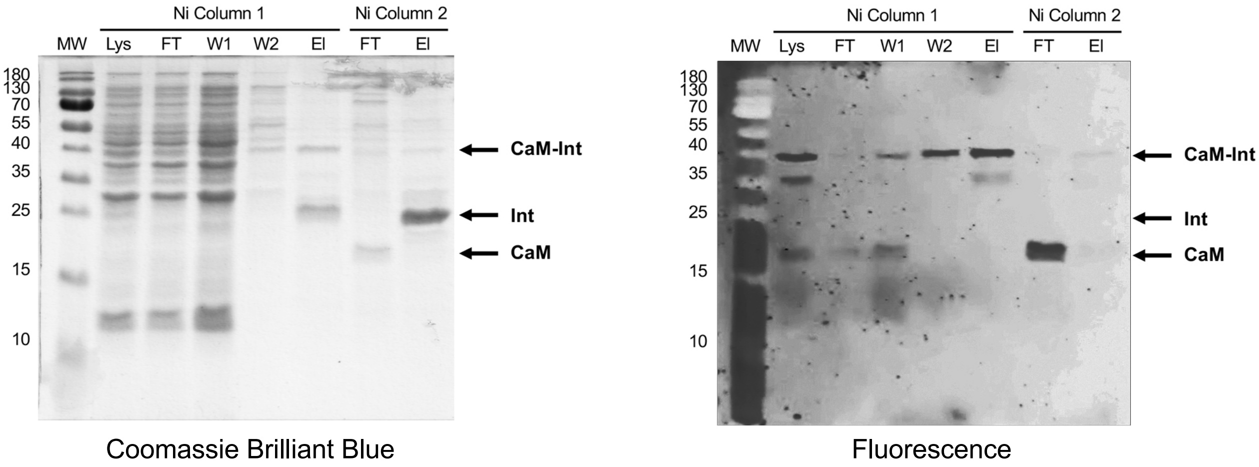


**Fig. S15.** Gel analysis of Acd incorporation in CaM using various RSs. MW = molecular weight in kDa. Lys = lysate, FT = flow through, W1 = wash 1, W2 = wash 2, El = elution , Int = intein.

**Purified αS-$E_{114}$δ**

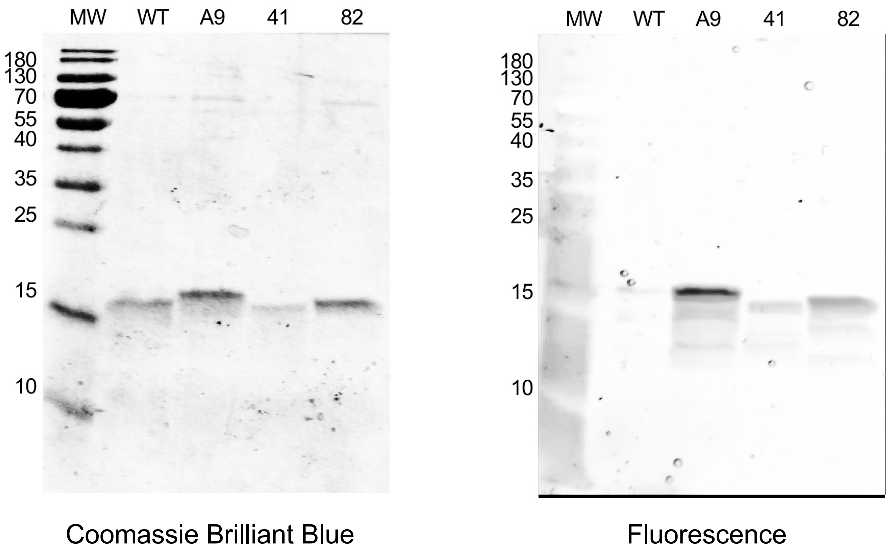


**Purified CaM-$L_{113}$δ**

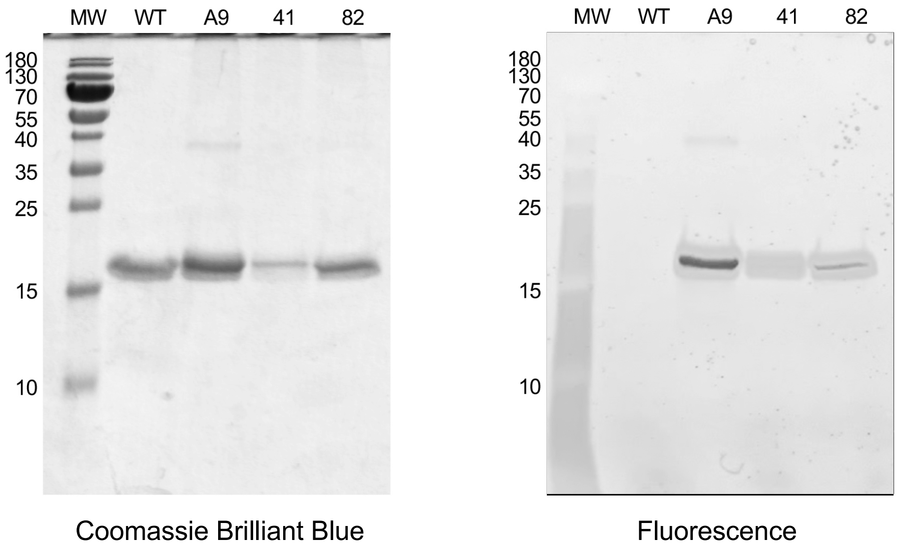


**Fig. S16.** Gel analysis of purified Acd and CaM proteins expressed using various RSs. MW = molecular weight in kDa.

### MALDI Spectra

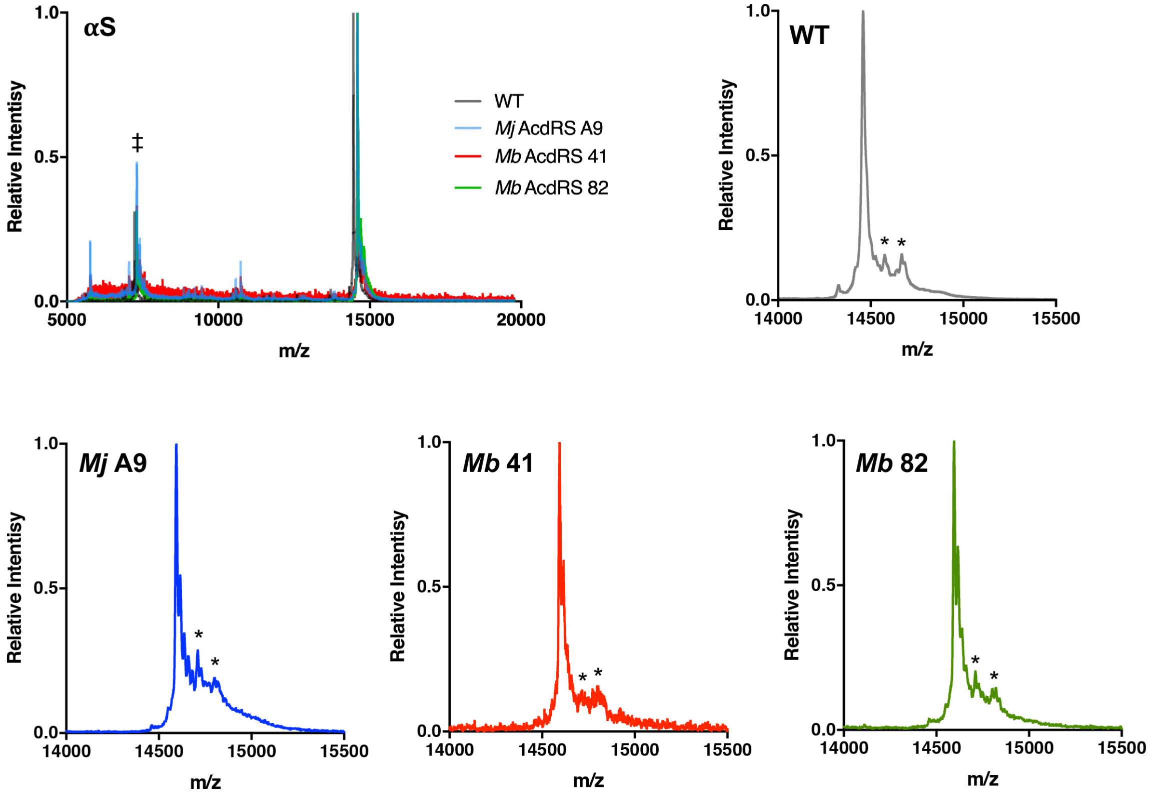


**Fig. S17.** MALDI MS Analysis of Acd Incorporation in αS. Expressions of αS-$E_{114}$δ using *Mj* AcdRS A9, *Mb* AcdRS 41, or *Mb* AcdRS 82 were performed in parallel. WT αS was also expressed for comparison. MALDI MS data showed masses corresponding to Acd incorporation with no peaks corresponding to mis-incorporation of a canonical amino acid. ‡indicates the doubly charged species. * indicates a known matrix adduct.[18]

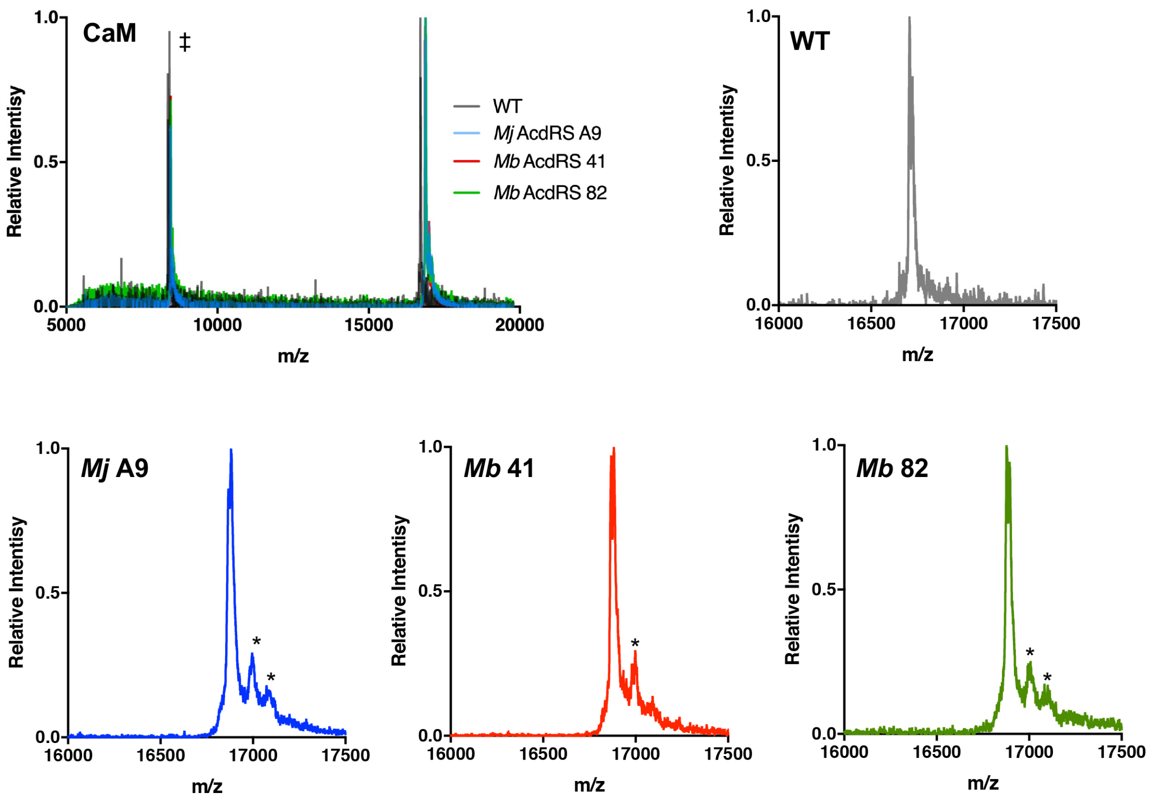


**Fig. S18.** MALDI MS Analysis of Acd Incorporation in CaM. Expressions of CaM- $L_{113}\delta$ using *Mj* AcdRS A9, *Mb* AcdRS 41, or *Mb* AcdRS 82 were performed in parallel. WT CaM was also expressed for comparison. MALDI MS data showed masses corresponding to Acd incorporation with no peaks corresponding to mis-incorporation of a canonical amino acid. ‡indicates the doubly charged species. * indicates a known matrix adduct.[18]

**Cloning of AcdRS into pUltra vector.** Primers were dissolved in Milli-Q grade water to be at a concentration of 50 μM. The amount of plasmid used ranged from 0.1 – 1 ng total DNA. Primers, plasmid and buffer components were mixed according to manufacturer recommendations. Vector (pUltra Mb-AcKRS, a gift from Abhishek Chatterjee) and insert (pDule AcdRS41 or pDule AcdRS82) were amplified simultaneously. PCR was carried out for 30 cycles with an annealing temperature of 67 °C, as calculated with the New England Biolabs Tm calculator tool (https://tmcalculator.neb.com). PCR product was purified using a 1% Agarose gel containing GelGreen Dye. Gel was visualized using a Blue light transillluminator and bands of the expected size were excised. DNA was extracted was extracted using a DNA gel extraction kit according to manufacturer's protocol and isolated DNA was quantified. DNA assembly was performed with 100 ng of vector DNA and 2 equiv of insert DNA using a HiFi DNA Assembly Mix according to manufacturer's protocol. 2 μL of the ligated product were transformed in 50 μL competent NEB10β cells according to manufacturer's instructions and plated on an LB-Agar plate containing Spectinomycin. Individual colonies were picked, grown up in 5 mL LB-medium and their DNA was extracted using a Plasmid Miniprep Kit according to manufacturer's protocol. Quantified DNA was submitted for sanger sequence analysis to verify correct sequence.

Forward primer for vector:
5′–TACTATAACGGCATTAGCACGAACCT–3′
Reverse complement primer for vector:
5′–GCAGGTTTTGCGGTATTTATGATGAC–3′
Forward primer for insert:
5′–GCGTGCGTTTCGTCATCATAAATAC–3′
Reverse complement primer for insert:
5′–CAGGTTCGTGCTAATGCCGTT–3′

**Cloning of αS-Mxe-$H_6$ and αS-$TAG_{94}$_Mxe-$H_6$ into pET22b vector.** Primers were dissolved in Milli-Q grade water to be at a concentration of 50 μM. The amount of plasmid used ranged from 0.1 – 1 ng total DNA. Primers, plasmid and buffer components were mixed according to manufacturer recommendations. Vector (pET22b sfGFP, a gift from Abhishek Chatterjee) and insert (pTXB1 αS-Mxe-$H_6$ or pTXB1 αS-$TAG_{94}$_Mxe-$H_6$) were amplified simultaneously using a touchdown protocol. PCR was carried out for 10 cycles with an initial annealing temperature of 65 °C in the first cycle and was lowered by 0.5 °C in each subsequent cycle. PCR was continued with an annealing temperature of 60 °C for another 25 cycles. PCR product was purified using a 1% Agarose gel containing

GelGreen Dye. Gel was visualized using a Blue light transilluminator and bands of the expected size were excised. DNA was extracted was extracted using a DNA gel extraction kit according to manufacturer's protocol and isolated DNA was quantified. DNA assembly was performed with 100 ng of vector DNA and 2 equiv of insert DNA using a HiFi DNA Assembly Mix according to manufacturer's protocol. 2 μL of the ligated product were transformed in 50 μL competent NEB10β cells according to manufacturer's instructions and plated on an LB-Agar plate containing Ampicillin. Individual colonies were picked, grown up in 5 mL LB-medium and their DNA was extracted using a Plasmid Miniprep Kit according to manufacturer's protocol. Quantified DNA was submitted for sanger sequence analysis to verify correct sequence.

Forward primer for vector:
5′—CATCATCATTAAAAGCTTAATTAGCTG—3′
Reverse complement primer for vector:
5′—GAATACATCCATATGTAATTTCTCCTCTTT—3′

Forward primer for insert:
5′—AAAGAGGAGAAATTACATATGGATGTATTCATGAAA—3′
Reverse complement primer for insert:
5′—ATTAAGCTTTTAATGATGATGATGATGATG—3′

**Expression Using C321 Cells.** pET22b_αS-Mxe-$H_6$ was transformed into chemically competent C321 cells (Addgene # 49018) via chemical transformation and plated on LB-Agar plates containing Ampicillin. pET22b_αS-$TAG_{94}$-Mxe-$H_6$ and pUltra_AcdRS41/82 were transformed simultaneously into electrocompetent C321 cells via electroporation and plated on LB-Agar plates containing Ampicillin and Spectinomycin. Individual colonies from each plate picked and grown overnight at 37 °C in 3 mL LB media containing appropriate antibiotics. The next day, 25 mL 2xYT media in 125 mL Erlenmeyer flasks with appropriate antibiotics was inoculated and grown in an incushaker at 37 °C and 250 RPM. Once cultures reached an $OD_{600}$ of 0.6-0.9, flasks were transferred to incushaker set to 18 °C and 250 RPM. Acd dissolved in water with addition of 5M NaOH and added to cultures at a final concentration of 1 mM. After 15 minutes, IPTG added at a final concentration of 1 mM and culture grown at 18°C overnight. Next day cells pelleted by centrifugation. Lysis was performed through resuspension in Per-B lysis solution supplemented with 1 mM EDTA according to manufacturer's protocol. Cell lysate pelleted in microcentrifuge tubes by centrifuging at 13.1k RPM at 4 °C for 1 hour. For each culture, 0.5 mL Ni-NTA resin washed with 8 mL equilibration buffer (50 mM HEPES, pH 7.5). Cell lysate added to resin and incubated on ice for 1 hour, after which the liquid was drained. Resin was washed with 5 mL equilibration buffer, 10 mL of wash buffer (50 mM HEPES, 10 mM imidazole, pH 7.5) and finally eluted with 1 mL elution buffer (50 mM HEPES, 300 mM imidazole, pH 7.5). 15 µL βME were added to induce intein cleavage and reaction was incubated overnight at room temperature. Next day, samples were dialyzed against 20 mM Tris, pH 8.0 to remove excess βME and imidazole. Samples were analyzed by SDS-PAGE and MALDI-TOF MS (Fig. S19).

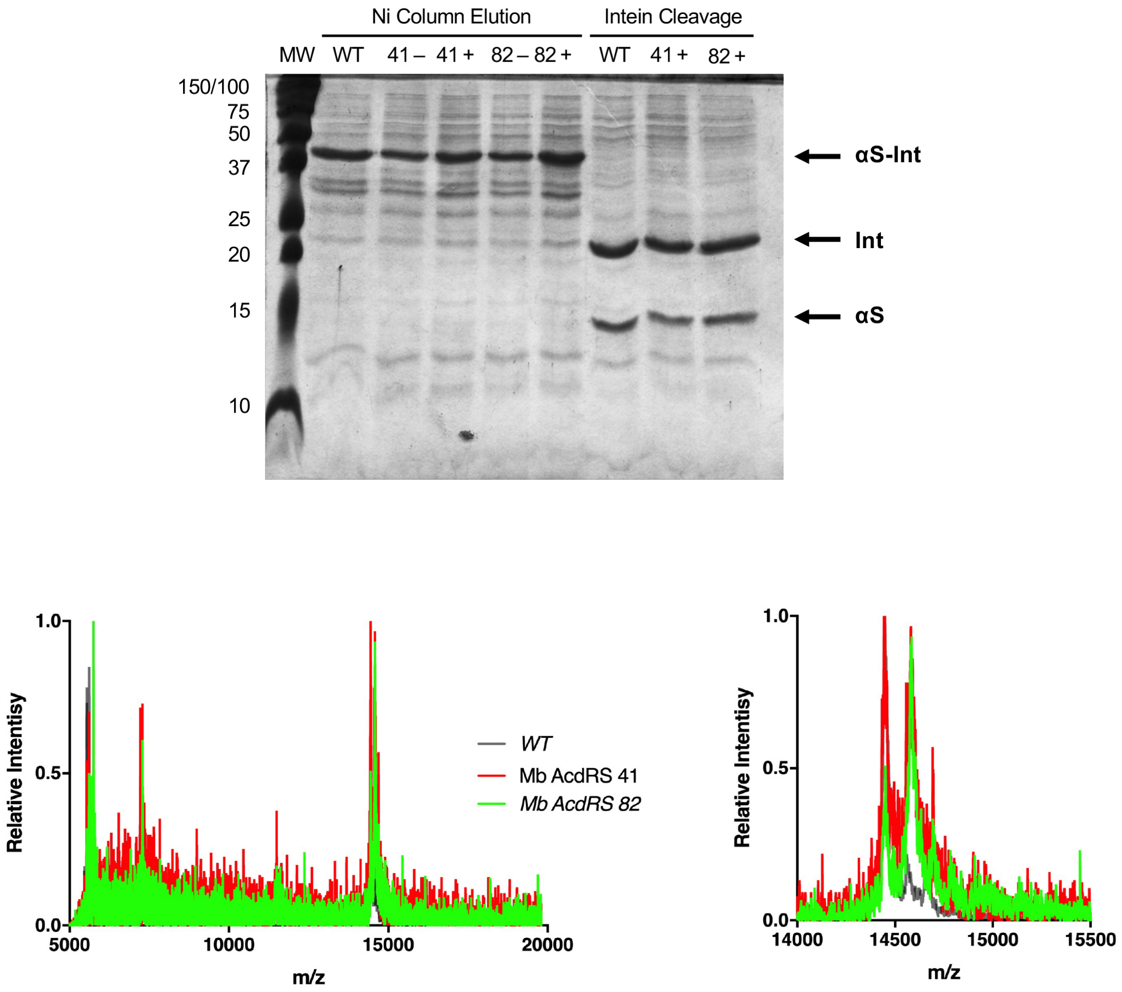


**Fig. S19.** Gel and MALDI MS analysis of Acd incorporation in αS using various aaRSs with pUltra plasmid and C321 cells. Top: Elutions are shown for expressions using *Mb* AcdRS 41 and 82 with (+) or without (–) Acd added. Expressions with Acd underwent intein cleavage and MALDI analysis. MW = molecular weight in kDa. Bottom: MALDI analysis indicates significant mis-incorporation for both RS. Expected masses 14,460 m/z (WT) and 14,577 m/z (Acd incorporation).

*Comparison of Protein Expression and Imaging in HEK Cells*

**Cloning RSs into pAcBac1.tR4-MbPyl.** Mammalian codon optimized genes for AcdRS 41 and AcdRS 82 were synthesized individually by BioBasic (Ontario, Canada) in a pUC57 plasmid. During gene synthesis, the first methionine from both RSs was removed and a nuclear export signal (NES) was added at the 5' end of the open reading frame for both RSs. The pUC plasmids containing both the AcdRS 41 and AcdRS 82 with NES (included to increase the amount of functional, cytoplasmic RS)[19] were digested with the restriction enzymes EcoRI and NheI. The same enzymes were used to excise the *Mb* PylRS gene from and pAcBac1.tR4-MbPyl (Addgene; Watertown, MA, USA). The desired RS gene fragments and the vector fragment were gel purified and ligated to give pAcBac1.tR4-AcdRS82 and pAcBac1.tR4-AcdRS41. After transforming to NEB5alpha competent cells (New England BioLabs; Ipswich, MA, USA) screening for colonies containing the RS gene in pAcBac vector were performed by restriction analysis. Identity of ligation products was confirmed by sequencing (Genewiz; Seattle, WA, USA and Eurofins; Louisville, KY, USA).

**HEK293T/17 Cell Maintenance**. HEK293T/17 cells were purchased from ATCC (CRL-11268, Manassas, VA, USA). Growth medium was prepared with DMEM with pyruvate and high glucose supplement with 10% FBS and Penicillin/Streptomycin (Pen/Strep, 50U/mL) (Invitrogen). Cells were maintained at 37 ºC with 5% $CO_2$ and passed at 1:10 or 1:20 dilution every 3 to 4 days.

**Acd Toxicity Screen**. HEK293T/17 cells were plated in a white Nuclon Delta-treated flat bottom 96-well plate (Thermo, Waltham, MA) at approximately 10% confluency in 100 µL of DMEM 10% FBS media containing Penn/Strep. Prior to incubation, a 0.8 M Acd stock solution was made by adding 1 M sodium hydroxide 50 µL at a time until dissolved, the solution was the diluted with sterile water to the desired final volume. This starting stock was diluted to 3.3 µM-331 mM with sterile water and heated to 90 °C for 15 minutes to fully sterilize the stock. Cells in 100 µL of media were incubated for 48 h at 37 °C under 5% $CO_2$ with 2.5 µL of room temperature Acd stock neutralized with 2.5 µL of 1 M HEPES buffer (pH 7.4). Additionally, cells were incubated with and without 2.5 µL of HEPES buffer for vehicle and untreated controls respectively. Cell viability was measured using a CellTiter Glo assay kit (Promega; Madison, WI, USA) according to the manufacturer's instructions. Briefly, 100 µl of CellTiter Glo reagent was added to each well and incubated for 10 minutes at room temperature. Then, luminescence emission was measured using a Tecan Spark M10 (Männedorf, Switzerland). The data was normalized to vehicle control and fitted to an [Inhibitor vs response – Variable slope] function by nonlinear regression using GraphPad Prism 5. Mild Acd toxicity was observed at concentrations greater than 600 μM. Two biological replicates and 5 technical replicates were performed.

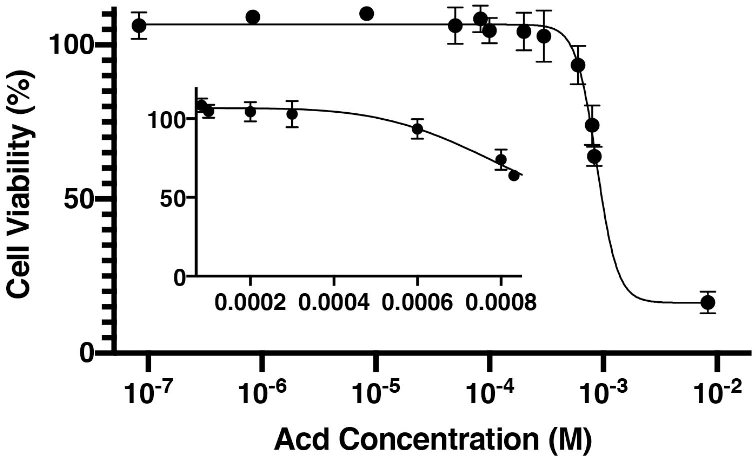


**Fig. S20** The dependence of cell viability on Acd concentration. Cell viability was measured using a CellTiter Glo assay kit and the data was normalized to untreated and vehicle control cells (n=5). The data were fit by an [Inhibitor vs response – Variable slope] function. Inset shows 0.1-1mM range.

**General Transfection protocol.** HEK293T/17 cells were plated in a 6-well plate at 20-30% confluency so that they reached 40-60% confluency at the time of transfection. Prior to transfection, the cell media was changed to antibiotic free DMEM containing 10% FBS. Cells were transfected with a total of 1.6 μg of DNA and 10 μL Lipofectamine 2000 (Invitrogen; Carlsbad, CA, USA) in 300 uL Opti-MEM (Invitrogen, Carlsbad, CA) per well using the manufacturer's protocol with minor modifications. Briefly, for each condition, 0.9 μg of pEGFP-C1-Y40TAG or pEGFP-C1 (WT), 0.3 μg of pAcBac1.tR4-AcdRS82 or pAcBac1.tR4-AcdRS41, and 0.4 μg of pDN-eRF1 were dissolved in 150 μL of Opti-MEM. In a separate tube, 10 μl of (1 mg/mL) Lipofectamine 2000 reagent was diluted in 150 μL of Opti-MEM. Both solutions were incubated for 5 minutes at room temperature, then combined into the Lipofectamine 2000 tube. After vigorous shaking, the solution was incubated at room temperature for a minimum of 20 minutes. The contents of each tube were then added to one well of the six-well plate. After 4-6 hours of incubation, the medium was changed to 2 mL of DMEM containing 10% FBS and Pen/Strep. Acd was dissolved in sterile water to a final concentration of 20 mM with dropwise addition of NaOH to aid dissolution. This solution was neutralized by combining 30 µL of 20 mM Acd with 30 µL of 1 M sterile HEPES buffer (pH 7.4) and added to one well of the 6-well plate. To wash excess Acd, the media was changed (DMEM containing 10% FBS and Penn/Strep) at 24- and 36-hours post-transfection. This protocol was used for most HEK cell expression experiments unless otherwise noted.

**Cell Imaging and Processing for Transfection Optimization Assays.** Cells were imaged at desired time points using an Olympus CKX53 microscope. Bright field images were obtained using brightfield (4000K color temperature LED light source). Fluorescence images were obtained by excitation using a 130W metal halide light source and a blue excitation filter (460-495 nm) with a 500 nm dichroic mirror and 510 nm long pass barrier filter. All images were obtained using 10X magnification. The green channel integrated fluorescence density was measured limited by a global threshold of 44-500 using Fiji software.[20]

**Cell harvesting, sonication, and lysate fluorescence quantification.** After transfection and protein expression in the presence or absence of Acd, cells were washed twice with DPBS containing calcium and magnesium. They were resuspended in 2 mL of DPBS without calcium and magnesium and detached via scraping. The cells were harvested by centrifugation at 1500 RPM for 10 minutes in an accuSPin 8C clinical centrifuge (Fisher Scientific; Pittsburgh, PA). The supernatant was removed and the pellet was frozen at -80 °C. The frozen pellet was thawed in 130 mM KCl, 30 mM Tris, pH 7.4 with cOmplete mini EDTA-free protease inhibitor cocktail and lysed using a SFX250 Sonifier (Branson Ultrasonics; Brookfield, Connecticut) set at 50% duty cycle, power = 4 one second on, two seconds off for ten pulses. The sonicated resuspension was pelleted at 14,000 RPM for 30 minutes on a 5415 R centrifuge (Eppendorf AG, Hamburg, Germany). EGFP fluorescence was measured on 100 µL of the supernatant using a Tecan M1000 plate reader (Mannedorf, Switzerland). Excitation was set at 470 nm, and emission from 485 to 600 nm was collected. Slit widths of 5 nm were used for both excitation and emission.

**Comparison of AcdRS incorporation efficiency.** Cells were plated in 6-well dishes. *Mb* AcdRS 41 and *Mb* AcdRS 82 were separately transfected into HEK293T/17 cells together with DN-eRF1 and EGFP-$Y_{40}\delta$ using the general transfection protocol. As a negative control, cells were transfected with DN-eRF1 and EGFP-$Y_{40}\delta$ in the absence of an AcdRS plasmid. As a positive control, they were transfected with WT EGFP and DN-eRF1. After 6 hours, cells were incubated in the presence or absence of 300 µM Acd. To wash away excess Acd, the media was changed with DMEM containing 10% FBS and Penn/Strep at 24- and 36-hours post-transfection. After 24 and 36 hours, each replicate was imaged on an Olympus CKX53 microscope, then cells were lysed and EGFP fluorescence was measured using the protocols described above. Microscopy images are shown in Fig. S21 and quantified data from lysates are shown in main text Fig. 4. Incorporation efficiency was determined by subtracting the EGFP emission observed under no RS conditions with Acd from EGFP emission for WT, AcdRS 41, and AcdRS 82 expressions with Acd and 41 and 82, then normalizing to corrected WT emission. This gave efficiencies of 29 ± 4% for AcdRS 82 and 5 ± 1% for AcdRS 41.

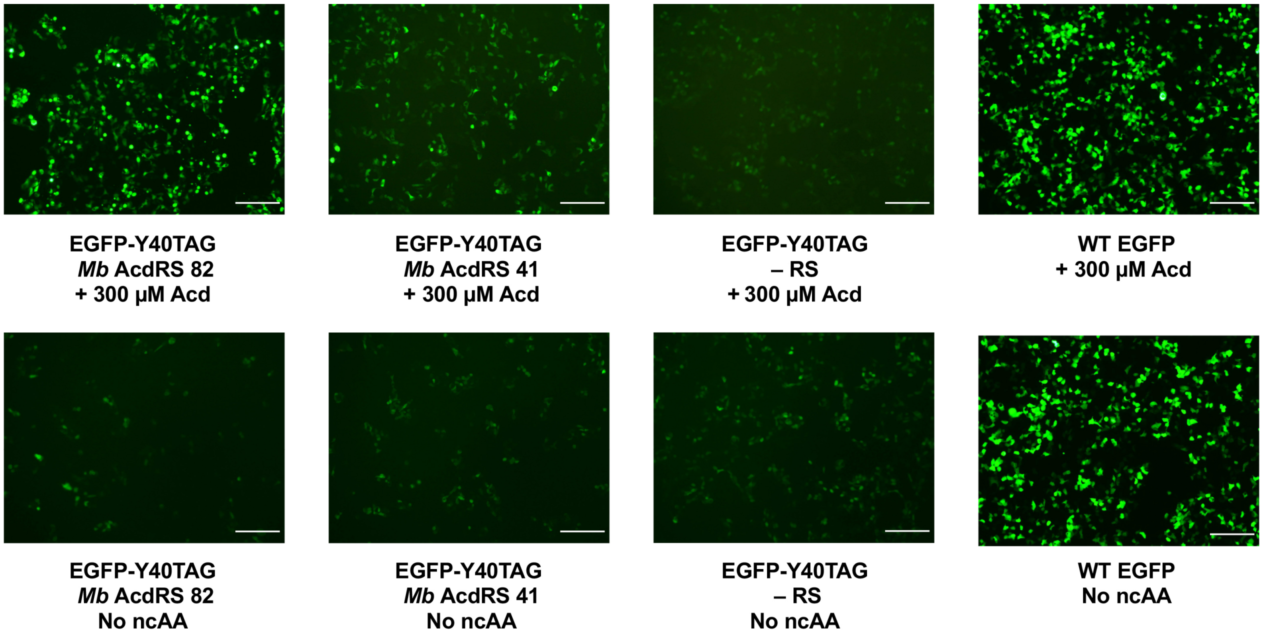


**Fig. S21.** Images 36 hours post-transfection of EGFP fluorescence from suppression using EGFP-Y40TAG plasmid and *Mb* AcdRS plasmids, a negative control lacking an AcdRS plasmid, and WT EGFP in the presence or absence of 300 µM Acd. The DN-eRF1 plasmid was also present in all cases. Scale bar = 200 µm.

**Effect of eRF1 on transfected protein production.** Cells were plated in a 24-well plate and followed the general transfection protocol, but used the following plasmid combinations: 0.18 µg of pEGFP-C1-Y40TAG or pEGFP-C1 DNA, and 0.06 µg *Mb* AcdRS 82 DNA, with or without 0.08 µg DN-eRF1 DNA. Each DNA combination was tested in the presence or absence of 300 µM Acd. At 36 hours post-transfection, the cells were imaged using the Olympus CKX53 microscope as described above. Measured EGFP fluorescence indicated that DN-eRF1 had little to no effect of WT EGFP production, but increased EGFP-$Y_{40}\delta$ protein production by roughly 5-fold. Additionally, DN-eRF1 showed a 2.8-fold increase in read-through of EGFP-$Y_{40}$TAG in the absence of Acd.

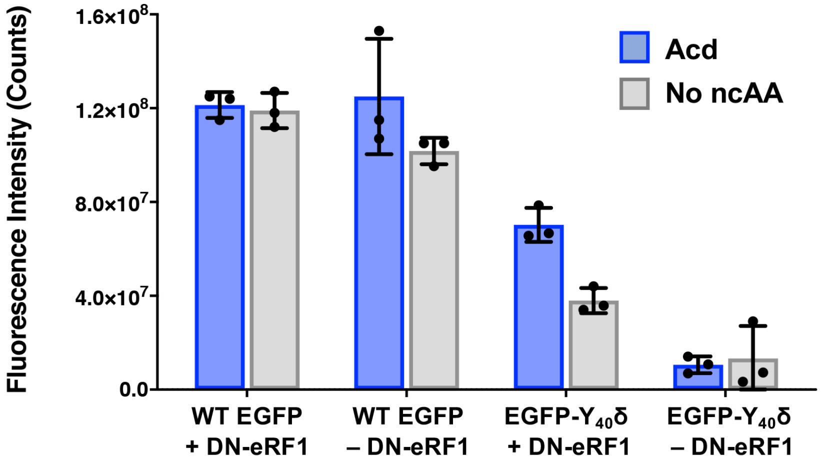


**Fig. S22.** Effect of DN-eRF1 on EGFP expression. EGFP fluorescence intensity measured from images using quantification in Fiji 36 hours post-transfection using EGFP-Y40TAG plasmid and *Mb* AcdRS 82 plasmids or WT EGFP plasmid in the presence or absence of 300 µM Acd. The DN-eRF1 plasmid was present or absent as indicated. n = 3.

**Optimizing expression conditions.** HEK293T/17 cells were plated in 6-well tissue culture trays and were transfected with 1.6 µg of total DNA using the general transfection protocol. Four hours post-transfection, the media was changed to DMEM 10% FBS containing Pen/Strep. Additionally, 60 µL of 50:50 1 M HEPES/Acd (40 mM-1.66 mM) was added to the medium resulting in final Acd concentrations of 25, 50, 100, 150, 200, 300, or 600 µM. The medium was changed at 24 hours post transfection and again at 36 hours post transfection. Each replicate (n=3) was imaged using the Olympus CKX53 microscope in bright field and fluorescence modes as described above. Then, the cells were harvested, sonicated, and EGFP fluorescence intensity was measured using the Tecan

M1000 plate reader as described above. Microscopy images are shown in Fig. S23 and quantified data from lysates are shown in main text Fig. 4.

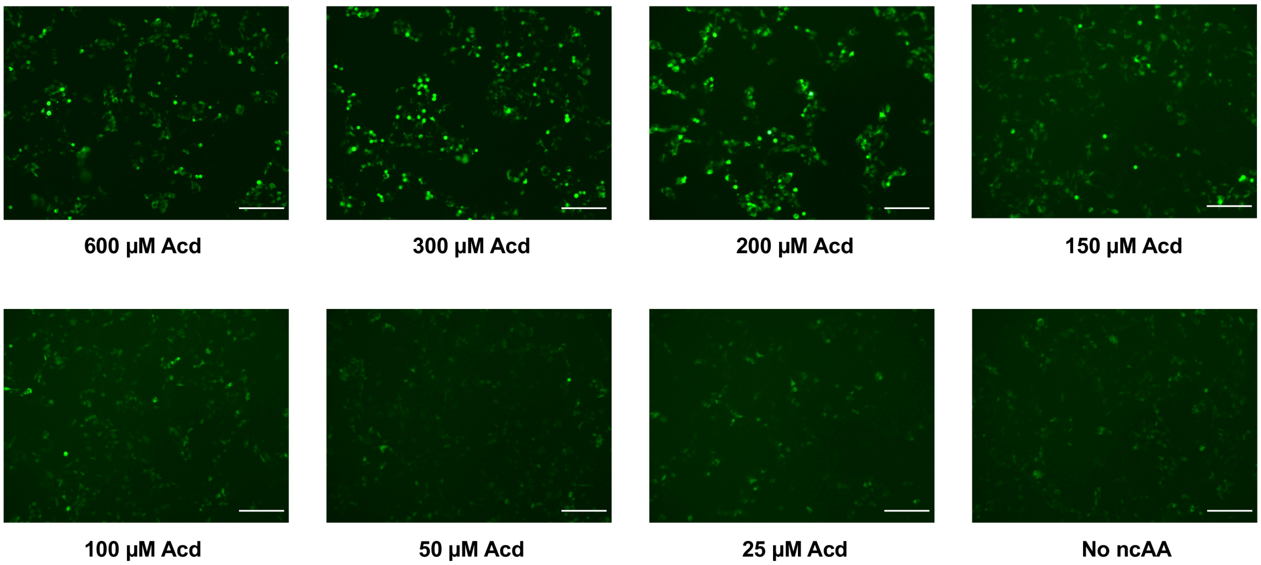


**Fig. S23.** Images of EGFP fluorescence from suppression using EGFP-Y40TAG, *Mb* AcdRS 82, and DN-eRF1 plasmids in the presence of varying concentrations of Acd. Scale bar = 200 μm.

To investigate the optimal DNA/Lipofectamine 2000 concentration, cells were transfected in a 24-well plate as in the general transfection protocol in the presence or absence of 300 μM Acd, but varying total DNA amounts (in the same ratios given above for the general transfection protocol, but scaled down for the 24 well plate) and DNA/Lipofectamine ratios as follows: 0.32 μg DNA (1:0.5, 1:1, 1:3, or 1:5 ratio), 0.64 μg DNA (1:1, 1:3, or 1:5 ratio), 0.32 μg WT EGFP DNA (1:5 ratio) , or no DNA. At 36 hours post-transfection, the cells were imaged and fluorescence quantified in Fiji as described. We observed that the 1:5 DNA/Lipofectamine ratio was optimal (we did not test higher ratios as the manufacturer recommends against this). We also observed that doubling the total amount of DNA led to a 2.3-fold increase in EGFP fluorescence, but also a 5.4-fold increase in EGFP fluorescence in the absence of Acd. Therefore, we chose to use a 1:5 DNA/Lipofectamine ratio for subsequent experiments with 0.32 μg total DNA (the equivalent of 1.6 μg DNA in a six well plate) since we were more concerned with eliminating read through of the TAG codon than maximizing the number of positive cells.

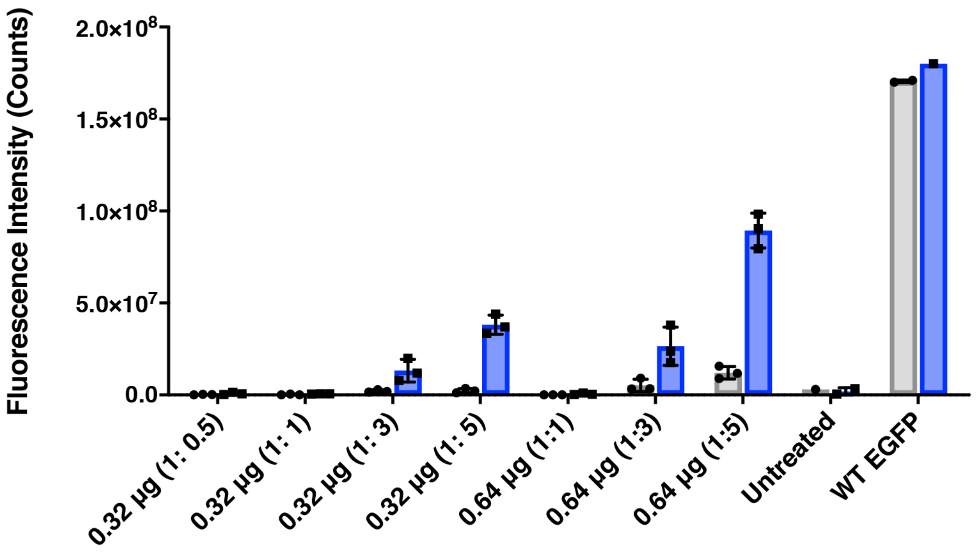


**Fig. S24.** EGFP fluorescence intensity measured from images using quantification in Fiji 36 hours post-transfection using EGFP-Y40TAG plasmid and *Mb* AcdRS 41 or 82 plasmids or WT EGFP plasmid in the presence of 300 µM Acd. The DN-eRF1 plasmid was present in all cases. DNA amounts and DNA/Lipofectamine ratios were varied as indicated. n = 3.

**Acd expression protocol effects on cell morphology.** HEK cells were transfected with DN-ERF1 and either pEGFP-C1 (WT) or pEGFP-C1-Y40TAG with pAcBac1.tR4-AcdRS82 in 6-well plates, using 1.6 µg DNA, Lipofectamine 2000 (1:5 DNA/Lipofectamine) and 300 µM Acd. At 36 hours post-transfection and ~80% confluency, the cells were detached using trypsin-EDTA and seeded onto 35 mm glass bottom Petri dishes (MatTek; El Segundo, CA) and incubated overnight to allow for cell adherence. At 48 hours post-transfection, the cells were washed twice with DPBS (containing calcium and magnesium) and were resuspended in DMEM (10% FBS, Pen/Strep) without Phenol Red. Cells were immediately imaged using a Nikon Eclipse Ti2 inverted microscope with ANDOR iXon Ultra 897, Yokogawa CSU-X1 spinning disk confocal with 488 nm excitation at 6% laser power, and the FITC emission filter (525-536 nm). The objective lens used was a CFI60 Apochromat Lambda S 60X Oil immersion lens, N.A. 1.4, W.D. 0.14 mm, F.O.V. 22 mm, DIC, Spring Loaded. A Z-stack with 0.5 micron spacing (7 images total) was acquired. The max intensity slice is shown for both brightfield and GFP fluorescence channels in Figure S25. All image brightness/contrasts were set to the same scale in Fiji.[20] We see that cell morphology is not significantly affected by Acd treatment or AcdRS plasmid transfection.

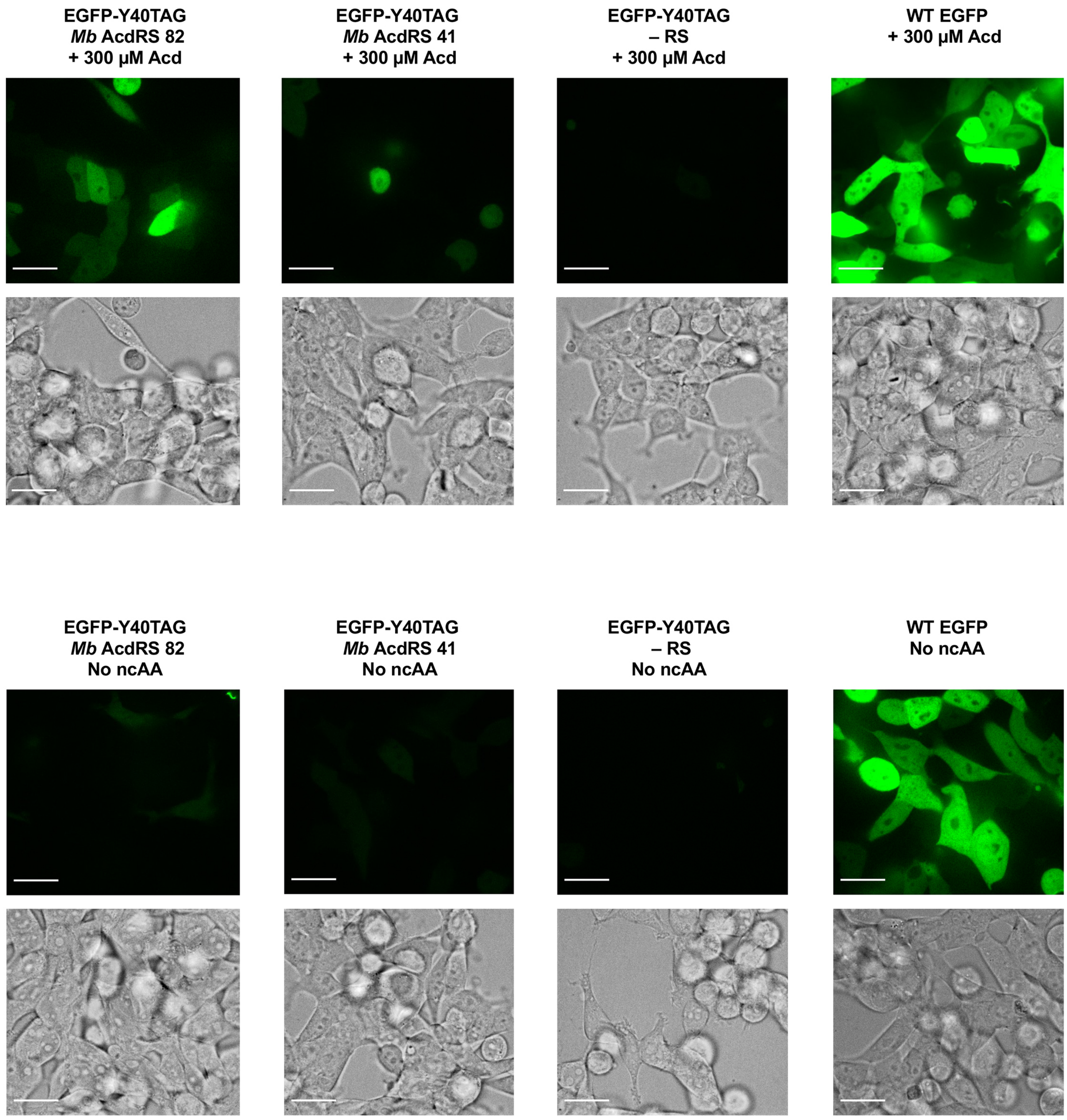


**Fig. S25.** Images 48 hours post-transfection of EGFP fluorescence from suppression using EGFP-Y40TAG plasmid and *Mb* AcdRS plasmids, a negative control lacking an AcdRS plasmid (– RS), and WT EGFP in the presence (top) or absence (bottom) of 300 µM Acd. The DN-eRF1 plasmid was also present in all cases. Cells imaged in DMEM without Phenol Red using a Nikon Eclipse Ti2 inverted microscope with 488 nm excitation. Scale bar = 25 µm.

**Analysis of AcdRS fidelity in mammalian cells.** Analysis of AcdRS fidelity was performed in HEK293T cells using purified sfGFP-$N_{150}\delta$ variants expressed with previously reported plasmid pAcBac1-sfGFP-N150TAG,[6, 21] with WT sfGFP expressed using pAcBac1-wtsfGFP. HEK293T cells were transfected using jetPRIME transfection reagent (Polyplus Transfection; Illkirch, France) with minor alterations from the manufacture's protocol. Confluent HEK293T cells were split into 100 mm plates for an initial confluency of 35% (DMEM + 10% FBS, 14.5 mL). At a confluency of 60-80%, the HEK293T cells were transfected using jetPRIME. For both AcdRSs, 1.45 mL jetPRIME buffer, 2.6 µg of pAcBac1.tR4-AcdRS41 or pAcBac1.tR4-AcdRS82, 3.4 µg of pDN-eRF1, 7.75 µg of pAcBac1-sfGFP-N150TAG, and 27.5 µL of jetPRIME transfection reagent were added. For WT sfGFP, 1.45 mL jetPRIME buffer, 3.4 µg of pDN-eRF1, 7.75 µg of pAcBac1-wtsfGFP and 22.3 µL of jetPRIME transfection reagent were added. Acd was then supplemented for a final concentration of 300 µM and cells were incubated for 24 hours and imaged on a Keyence BZ-X710 microscope to verify sfGFP expression. At 36 hours cells were imaged and pelleted for protein purification after three 10 mL DPBS washes.

Cell pellets were resuspended in 1 mL of lysis buffer (50 mM $Na_2PO_4$ pH 7.0, 500 mM NaCl, 5 mM imidazole) and subject to lysis via sonication using a Fisher Dismembrator Model 60 with microprobe. Cells were centrifuged in 1.7 mL microcentrifuge tubes for 30 minutes at 20,000 rcf, 4 °C and the soluble fractions were added to 100 µL TALON resin. The lysate was incubated with the resin and slowly rocked (1.5 hour, 4 °C). After binding, the resin and lysis were added to columns. The resin was washed with 30x the bed volume (3 mL lysis buffer) and then eluted into 300 µL elution buffer (50 mM $Na_2PO_4$, 500 mM NaCl, 250 mM imidazole). The sample was spin concentrated and desalted to a concentration of 50 µM in LC-MS grade water. Purified sfGFP samples were diluted to 10 µM, desalted on $C_4$ zip tips, and analyzed using the Waters Synapt G2 mass spectrometer at the Oregon State University Mass Spectrometry Facility in ESI mode. Spectra were deconvoluted using the Maximum Entropy deconvolution algorithm in BioConfirm. The masses of the proteins are given in Table S9 and the spectra are shown in Fig. S26.

Amino acid sequence of WT sfGFP (site of Acd incorporation at $N_{150}$ underlined)

```
MVSKGEELFTGVVPILVELDGDVNGHKFSVRGEGEGDATNGKLTLKFICTTGKLPVPWPTLVTTLTYGVQCF
SRYPDHMKRHDFFKSAMPEGYVQERTISFKDDGTYKTRAEVKFEGDTLVNRIELKGIDFKEDGNILGHKLEY
NFNSHNVYITADKQKNGIKANFKIRHNVEDGSVQLADHYQQNTPIGDGPVLLPDNHYLSTQSVLSKDPNEKR
DHMVLLEFVTAAGITHGMDELYKGSGKPIPNPLLGLDSTHHHHHH
```

**Table S9.** Masses of sfGFP variants expressed in HEK cells

| | Calc'd $[M+H]^+$ | Obsv'd $[M+H]^+$ |
|---|---|---|
| WT | 29142* | 29142 |
| *Mb* AcdRS 41 | 29292 | 29292 |
| *Mb* AcdRS 82 | 29292 | 29292 |

* Predicted average $[M+H]^+$ of unmodified WT sfGFP-$His_6$: 29251; GFP chromophore processing for fluorescence (-20): 29231; loss of N-terminal methionine (-131): 29100; N-terminal acetylation (+42): 29142; substitution of Acd for Asn150: 29292. All values in Da.

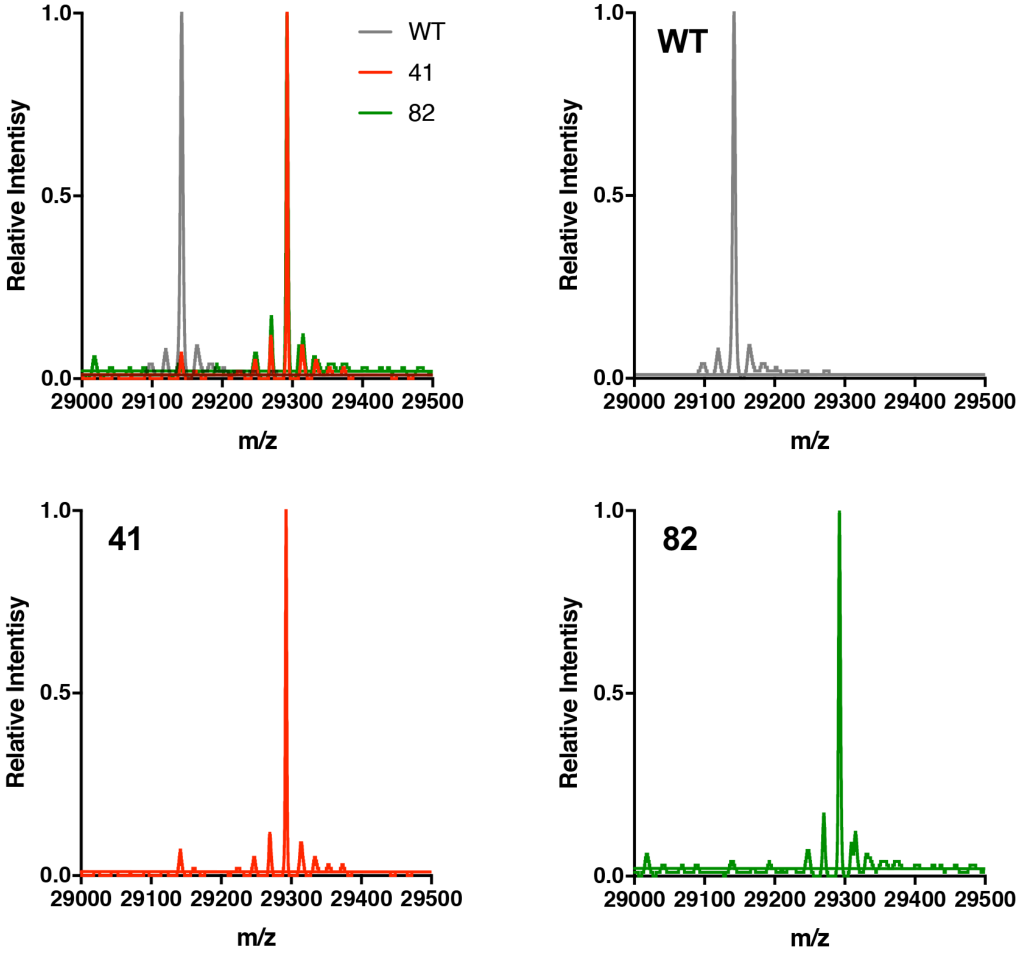


**Figure S26.** ESI MS data demonstrating Acd incorporation in sfGFP in HEK cells. Expressions of sfGFP-$N_{150}\delta$ using *Mb* AcdRS 41 or *Mb* AcdRS 82 were performed in parallel. WT sfGFP was also expressed for comparison. ESI MS data showed masses corresponding to Acd incorporation. Trace mis-incorporation of a canonical amino acid is observed for AcdRS 41. No mis-incorporation is observed for AcdRS 82.

**Confocal Microscopy and FLIM.** For confocal and FLIM experiments, the DN-eRF1 plasmid was used for all experiments, along with pIR-TAG676-GFP, pcDNA3.1HE-spHCN-TAG355-YFP, or pcDNA3-FLAG-MBP1-E322TAG plasmids. For + RS conditions, pAcBac1.tR4-AcdRS82 was included, for – RS conditions, this plasmid was absent but replaced with pcDNA3 to maintain the total DNA amount for transfection to be the same. For confocal imaging, HEK293T/17 cells were plated in 6-well tissue culture trays. Cells were transfected and incubated as in the EGFP conformal imaging experiment above, but at low confluency in order to ensure a cellular monolayer for imaging. The cells were replated to poly-K treated 8 well glass bottom plates (Ibidi) with growth medium containing ~450 μM Acd. Growth medium in each well was changed twice between transfection and imaging to dialyze out unincorporated Acd to lower background. Trays were wrapped with aluminum foil to block any light and incubated at 37 °C until use. Confocal microscopy images with protein expression are shown in Fig. 5 and expanded views of – RS conditions are shown in Fig. S27.

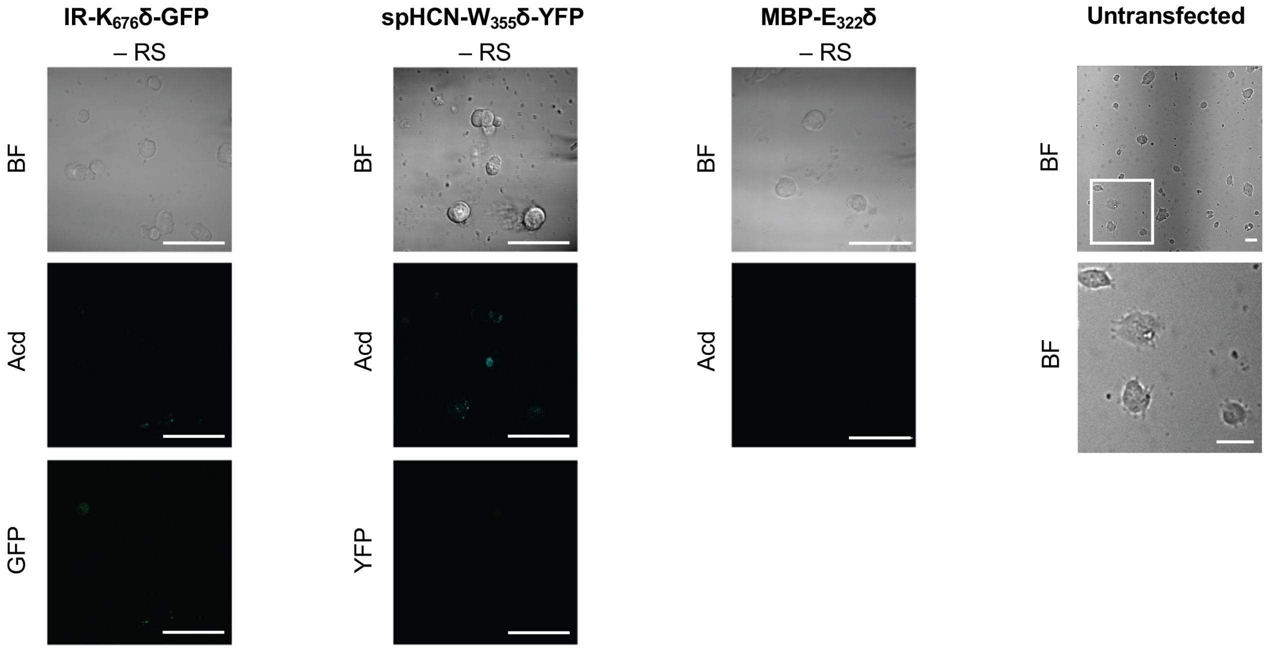


**Fig. S27.** Negative Controls for Acd Incorporation into Proteins in HEK Cells. Cells were transfected with the indicated plasmid for the protein of interest with a TAG codon and the DN-eRF1 plasmid, with or without (– RS) a plasmid encoding *Mb* AcdRS 82 and $tRNA_{CUA}$. Untransfected cells were treated with Acd and washed like transfected cells (shown at two levels of magnification). Confocal microscopy images show brightfield (BF), the Acd fluorescence channel (405 nm excitation; 409-480 nm emission), and the GFP (488 nm excitation; 500-621 nm emission) or YFP (514 nm excitation; 519-621 nm emission) channels. Scale bar = 50 μm.

For FLIM experiments, the gray scale images in Fig. 6 and Fig. S28 show the intensity of confocal emission at the indicated wavelengths of excitation. The phasor plot is a graphical representation of the raw fluorescence lifetime data in a vector space. The real (D) and imaginary (N) components of the lifetime, at 10 mHz, were calculated for each voxel via digital Fourier transform of its corresponding phase histogram, subject to the instrument response function (IRF) calibration using a standard sample of known lifetime. In these phasor plots, points corresponding to major lifetimes can be observed (circles), with the points in between representing a mixture of the species in different ratios. The red highlighted points in the image represents voxels from the data within the red circle on the phasor plot and the blue highlighted points represent voxels from the data within the blue circle.

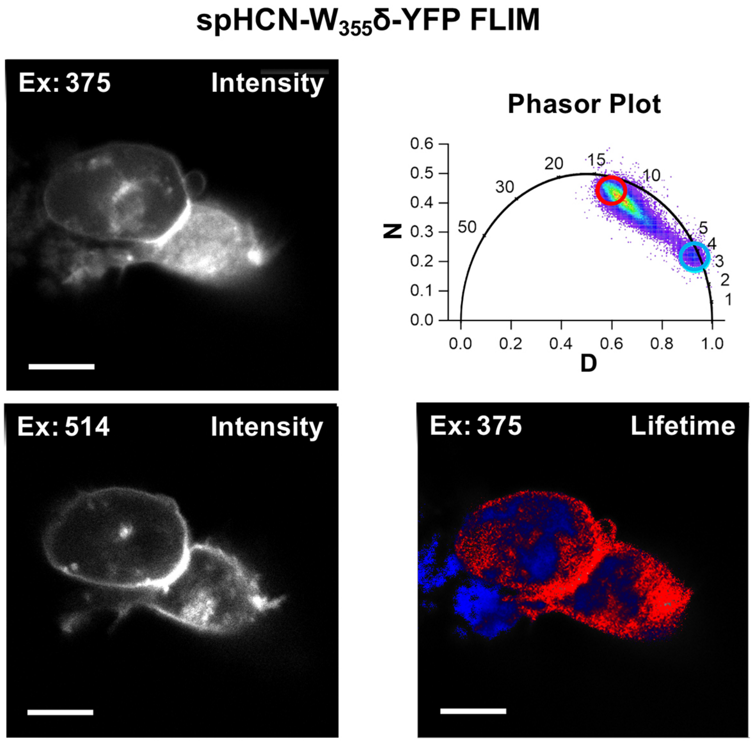


**Fig. S28.** Acd Fluorescence Lifetime Imaging (FLIM). FLIM allows Acd fluorescence to be separated from autofluorescence in living cells. spHCN-$W_{355}$δ-YFP was expressed with *Mb* AcdRS 82 in HEK cells. Intensity plots (left) show emission after excitation at 375 or 514 nm. Phasor plot (top right) shows phase vectors D and N with points representing lifetimes of pixels. Circles indicate the selected lifetime ranges shown in red (≥ 15 ns) and blue (<5 ns) in the lifetime image (bottom right).

*AcdRS Sequences*

**pDULE2_MbAcdRS82 (RS sequence underlined):**

gcgatgagcgaaatgtagtgcttacgttgtcccgcatttggtacagcgcagtaaccggcaaaatcgcgccga
aggatgtcgctgccgactgggcaatggagcgcctgccggcccagtatcagcccgtcatacttgaagctagac
aggcttatcttggacaagaagaagatcgcttggcctcgcgcgcagatcagttggaagaatttgtccactacg
tgaaaggcgagatcaccaaggtagtcggcaaataatgtctaacaatgtctaacaattcgttcaagccgaggg
gccgcaagatccggccacgatgacccggtcgtcggttcagggcagggtcgttaaatagccgcttatgtctat
tgctggtttaccggggatcctctacgccggacgcatcgtggccggcatcaccggcgccacaggtgcggttgc
tggcgcctatatcgccgacatcaccgatggggaagatcgggctcgccacttcgggctcatgagcgcttgttt
cggcgtgggtatggtggcaggccccgtggccgggggactgttgggcgccatctccttgcatgcaccattcct
tgcggcggcggtgctcaacggcctcaacctactactgggctgcttcctaatgcaggagtcgcataagggaga
gcgtcgaccgatgcccttgagagccttcaacccagtcagctccttccggtgggcgcggggcatgactatcgt
cgccgcacttatgactgtcttctttatcatgcaactcgtaggacaggtgccggcagcgctctgggtcatttt
cggcgaggaccgctttcgctggagcgcgacgatgatcggcctgtcgcttgcggtattcggaatcttgcacgc
cctcgctcaagccttcgtcactggtcccgccaccaaacgtttcggcgagaagcaggccattatcgccggcat
ggcggccgacgcgctgggctacgtcttgctggcgttcgcgacgcgaggctggatggccttccccattatgat
tcttctcgcttccggcggcatcgggatgcccgcgttgcaggccatgctgtccaggcaggtagatgacgacca
tcagggacagcttcaaggatcgctcgcggctcttaccagcctaacttcgatcattggaccgctgatcgtcac
ggcgatttatgccgcctcggcgagcacatggaacgggttggcatggattgtaggcgccgccctataccttgt
ctgcctccccgcgttgcgtcgcggtgcatggagccgggccacctcgacctgaatggaagccggcggcacctc
gctaacggattcaccactccaagaattggagccaatcaattcttgcggagaactgtgaatgcgcaaaccaac
ccttggcagaacatatccatcgcgtccgccatctccagcagccgcacgcggcgcatctcgggctccttgcat
gcaccattccttgcggcggcggtgctcaacggcctcaacctactactgggctgcttcctaatgcaggagtcg
cataagggagagcgtctggcgaaaggggggatgtgctgcaaggcgattaagttgggtaacgccagggttttcc
cagtcacgacgttgtaaaacgacggccagtgccaagcttaaaaaaaatccttagctttcgctaagatctgca
gtggcggaaacccccgggaatctaacccggctgaacggatttagagtccattcgatctacatgatcaggttcc
cgcggccgcgaattcagcgttacaagtattacacaaagtttttttatgttgagaatatttttttgatggggcg
ccacttatttttgatcgttcgctcaaagaagcggcgccagggttgtttttcttttcaccagtgagacgggca
acagaacgccatgagcggcctcatttcttattctgagttacaacagtccgcaccgctgccggtagctccttc
cggtgggcgcggggcatgactatcgtcgccgcacttatgactgtcttctttatcatgcaactcgtaggacag
gtgccggcagcgcccaacagtcccccggccacggggcctgccaccatacccacgccgaaacaagcgccctgc
accattatgttccggatctgcatcgcaggatgctgctggctaccctgtggaacacctacggtacctcgggtt
gtcagcctgtcccgcttataagatcatacgccgttatacgttgtttacgctttgaggaatcccatatggata
aaaaaccgctggatgtgctgattagcgcgaccggcctgtggatgagccgtaccggcaccctgcataaaatca
aacatcatgaagtgagccgcagcaaaatctatattgaaatggcgtgcggcgatcatctggtggtgaacaaca
gccgtagctgccgtaccgcgcgtgcgtttcgtcatcataaataccgcaaaacctgcaaacgttgccgtgtga
gcgatgaagatatcaacaactttctgacccgtagcaccgaaagcaaaaacagcgtgaaagtgcgtgtggtga
gcgcgccgaaagtgaaaaaagcgatgccgaaaagcgtgagccgtgcgccgaaaccgctggaaaatagcgtga
gcgcgaaagcgagcaccaacaccagccgtagcgttccgagcccggcgaaaagcaccccgaacagcagcgttc
cggcgtctgcgccggcaccgagcctgacccgcagccagctggatcgtgtggaagcgctgctgtctccggaag
ataaaattagcctgaacatggcgaaaccgtttcgtgaactggaaccggaactggtgacccgtcgtaaaaacg
attttcagcgcctgtataccaacgatcgtgaagattatctgggcaaactggaacgtgatatcaccaaatttt
ttgtggatcgcggctttctggaaattaaaagcccgattctgattccggcggaatatgtggaacgtatgggca
ttaacaacgacaccgaactgagcaaacaaattttccgcgtggataaaaacctgtgcctgcgtccgatgctgg
ccccgaccctgtataactatctgcgtaaactggatcgtattctgccgggtccgatcaaaattttgaagtgg
gcccgtgctatcgcaaagaaagcgatggcaaagaacacctggaagaattcaccatggtttcgtttgggcaaa
tgggcagcggctgcacccgtgaaaacctggaagcgctgatcaaagaattcctggattatctggaaatcgact
tcgaaattgtgggcgatagctgcatggtgtatggcgataccctggatattatgcatggcgatctggaactga
gcagcgcggctgtgggtccggttagcctggatcgtgaatggggcattgataaaccgactattggcgcgggtt
ttggcctggaacgtctgctgaaagtgatgcatggcttcaaaaacattaaacgtgcgagccgtagcgaaagct
actataacggcattagcacgaacctgtaactgcagtttcaaacgctaaattgcctgatgcgctacgcttatc

aggcctacatgatctctgcaatatattgagtttgcgtgcttttgtaggccggataaggcgttcacgccgcat
ccggcaagaaacagcaaacaatccaaaacgccgcgttcagcggcgtttttctgcttttcttcgcgaattaa
ttccgctctcgagcgaagcttgngcccgaacaaaaactcatctcanaagaggatctgantaggagagaagat
tttcagcctgatacagattaaatcagaacgcagaagcggtctgataaaacagaatttgcctggcggcagtag
cgcggtggtcccacctgaccccatgccgaactcagaagtgaaacgccgtagcgccgatggtagtgtggggtc
tccccatgcgagagtagggaactgccaggcatcaaataaaacgaaaggctcagtcgaaagactgggcctttc
gttttatctgttgtttgtcggtgaacgctctcctgagtaggacaaatccgccgggagctgtccctcctgttc
agctactgacggggtggtgcgtaacggcaaaagcaccgccggacatcagcgctagcggagtgtatactggct
tactatgttggcactgatgagggtgtcagtgaagtgcttcatgtggcaggagaaaaaaggctgcaccggtgc
gtcagcagaatatgtgatacaggatatattccgcttcctcgctcactgactcgctacgctcggtcgttcgac
tgcggcgagcggaaatggcttacgaacggggcggagatttcctggaagatgccaggaagatacttaacaggg
aagtgagagggccgcggcaaagccgtttttccataggctccgcccccctgacaagcatcacgaaatctgacg
ctcaaatcagtggtggcgaaacccgacaggactataaagataccaggcgtttccccctggcggctccctcgt
gcgctctcctgttcctgcctttcggtttaccggtgtcattccgctgttatggccgcgtttgtctcattccac
gcctgacactcagttccgggtaggcagttcgctccaagctggactgtatgcacgaaccccccgttcagtccg
accgctgcgccttatccggtaactatcgtcttgagtccaacccggaaagacatgcaaaagcaccactggcag
cagccactggtaattgatttagaggagttagtcttgaagtcatgcgccggttaaggctaaactgaaggaca
agttttggtgactgcgctcctccaagccagttacctcggttcaaagagttggtagctcagagaaccttcgaa
aaaccgccctgcaaggcggttttttcgttttcagagcaagagattacgcgcagaccaaaacgatctcaagaa
gatcatcttattaatcagataaaatatttctagatttcagtgcaatttatctcttcaaatgtagcacctgaa
gtcagccccatacgatataagttgtaattctcatgtttgacagcttatcatcgatgcgaccgagtgagctag
ctatttgtttatttttctaaatacattcaaatatgtatccgctcatgagacaataaccctgataaatgcttc
aataatattgaaaaaggaagagtatgagggaagcggtgatcgccgaagtatcgactcaactatcagaggtag
ttggcgtcatcgagcgccatctcgaaccgacgttgctggccgtacatttgtacggctccgcagtggatggcg
gcctgaagccacacagtgatattgatttgctggttacggtgaccgtaaggcttgatgaaacaacgcggcgag
ctttgatcaacgaccttttggaaacttcggcttcccctggagagagcgagattctccgcgctgtagaagtca
ccattgttgtgcacgacgacatcattccgtggcgttatccagctaagcgcgaactgcaatttggagaatggc
agcgcaatgacattcttgcaggtatcttcgagccagccacgatcgacattgatctggctatcttgctgacaa
aagcaagagaacatagcgttgccttggtaggtccagcggcggaggaactctttgatccggttcctgaacagg
atctatttgaggcgctaaatgaaaccttaacgctatggaactcgccgcccgactgggctg

**pAcBac1.tR4-AcdRS82 (RS sequence highlighted in yellow):**

tattgaagcatttatcagggttattgtctcatgagcggatacatatttgaatgtatttagaaaaataaacaa
ataggggttccgcgcacatttccccgaaaagtgccacctaaattgtaagcgttaatattttgttaaaattcg
cgttaaatttttgttaaatcagctcattttttaaccaataggccgaaatcggcaaaatcccttataaatcaa
aagaatagaccgagatagggttgagtgttgttccagtttggaacaagagtccactattaaagaacgtggactc
caacgtcaaagggcgaaaaaccgtctatcagggcgatggcccactacgtgaaccatcaccctaatcaagttt
tttggggtcgaggtgccgtaaagcactaaatcggaaccctaaagggagcccccgatttagagcttgacgggg
aaagccggcgaacgtggcgagaaaggaagggaagaaagcgaaaggagcgggcgctagggcgctggcaagtgt
agcggtcacgctgcgcgtaaccaccacacccgccgcgcttaatgcgccgctacagggcgcgtcccattcgcc
attcaggctgcaaataagcgttgatattcagtcaattacaaacattaataacgaagagatgacagaaaaatt
ttcattctgtgacagagaaaaagtagccgaagatgacggtttgtcacatggagttggcaggatgtttgatta
aaaacataacaggaagaaaaatgccccgctgtgggcggacaaaatagttgggaactgggaggggtggaaatg
gagtttttaaggattatttagggaagagtgacaaaatagatgggaactgggtgtagcgtcgtaagctaatac
gaaaattaaaaatgacaaaatagtttggaactagatttcacttatctggttcggatctcctagcaaaaaacg
gggggtgcgggagtttcacccgcctgcacggatttagagtccgtgtgatctattcgatccacccccggtgt
ctctatcactgatagggaacttataagtctctatcactgatagggatttcacgtttatggtgatttcccaga
acacatagcgacatgcaaatattaaaaaacggaaaccccgggaatctaacccggctgaacggatttagagtc
cgttcgatctacatgatcaggtttccggtgtttcgtcctttccacaagatatataaagccaagaaatcgaaa
tactttcaagttacggtaagcatatgatagtccattttaaaacataattttaaaactgcaaactacccaaga
aattattactttctacgtcacgtattttgtactaatatctttgtgtttacagtcaaattaattctaattatc
tctctaacagccttgtatcgtatatgcaaatatgaaggaatcatgggaaataggccctcttcctgcccgacc
taggctcaagcagtgatcagatccagacatgataagatacattgatgagtttggacaaaccacaactagaat
gcagtgaaaaaaatgctttatttgtgaaatttgtgatgctattgctttatttgtaaccattataagctgcaa
taaacaagttaacaacaacaattgcattcattttatgtttcaggttcagggggaggtgtgggaggttttta
aagcaagtaaaacctctacaaatgtggtatggctgattatgatcctctagtacttctcgacaagcttggccg
gcctggccttatcactttccaagtcggttcatctctatgtctgtataaatctgtcttttcttggtgtgcttt
aatttaatgcaaagatggataccaactcggagaaccaagaatagtccaatgattaaccctatgataaagaaa
aaagaggcaatagagcttttccaactactgaaccaaccttctacaagctcgattggattttggatagccca
gtatcaccaaaaaataaactctcatcatcaggaagttgcgaagcagcgtcttgaatgtgaggatgttcgaac
acctgagcctttgagctaagatgaagatcggagtccaacataccatgtccaatcatgtataaaggaaactta
tatcctgaactggtcctcagaactccattgggtccaatttccacgtcttcatatggtgcccagtcatcccac
agttccctttctgtggtagttccactgatcattccgaccattcttgagaggattggagcagcaatatcgact
ctgatgtatctggtctcaaagtattttagggtaccattgattatggtgaaagcaggaccggttcctgggttt
ttaggagcaagatagctgagatccactggagagattggaagacccgctctgattttgctccaggtttcttgg
cagagggaataatccaagatcctctcaacgtcctgaattagacttacatccactgaggtctgagatggagca
gagatacttgacccttctgggcattcagggaatctggctgcagcaaagagatccttatcagccatctcgaac
cagacacctgatgggagtctgactccccaatgcttgcagtattgcattttgcaggccttgcctccagtttca
taagcaaagtagttacttctgaaccctgtgccctcctttcccagggatgatagctctccgtcctctgagaag
aaggtgatgtccatggaaatgaggttagaatcacatagccctttgaccttatagtcagaatgccaggttgta
gagttatggacagtggggcatatgtaattgctgcattttccgttgatgaactgtgaatcaacccattctcct
gtgtattcatcaaccagcacatggtgaggagtcacctggacaatcactgcttcggcatccgtcacagttgca
tatccacaactttgaggagggaagcctggattcagccaagttccttgtttcgtttgttcaatgctttccttg
cattgttctacagatggagtgaaggatcggatggaatgtgttatatacttcggtccataccagcggaaatca
caagtagtgacccatttggaagcatgacacatccaaccgtctgcttgaatagccttgtgactcttgggcatt
ttgacttgtaaggctgtgcctattaagtcattatgccaatttaaatctgagcttgacggcaataatggtaa
ttagaaggaacattttccagtttcctttttggttgtgtggaaaaactatggtgaacttgcaattcacccca
atgaataaaaaggctaagtacaaaaggcacttcatggtggcggtttttttaggagttcgcgcccgatggtggg
acggtatgaataatccggaatatttataggtttttttattacaaaactgttacgaaaacagtaaaatactta
tttatttgcgagatggttatcattttaattatctccatgatctattaatattccggagtatacggaccgcgg
ccgcaaatacctgcaggatccgttttgcgctgcttcgcgatgtacgggccagatatacgcgttgacattgat
tattgactagttattaatagtaatcaattacggggtcattagttcatagcccatatatggagttccgcgtta
cataacttacggtaaatggcccgcctggctgaccgcccaacgacccccgcccattgacgtcaataatgacgt
atgttcccatagtaacgccaatagggactttccattgacgtcaatgggtggagtatttacggtaaactgccc

acttggcagtacatcaagtgtatcatatgccaagtacgccccctattgacgtcaatgacggtaaatggcccg
cctggcattatgcccagtacatgaccttatgggactttcctacttggcagtacatctacgtattagtcatcg
ctattaccatggtgatgcggttttggcagtacatcaatgggcgtggatagcggtttgactcacggggatttc
caagtctccaccccattgacgtcaatgggagtttgttttggcaccaaaatcaacgggactttccaaaatgtc
gtaacaactccgccccattgacgcaaatgggcggtaggcgtgtacggtgggaggtctatataagcagagctc
tctggctaactagagaacccactgcttactggcttatcgaaattaatacgactcactatagggagacccaag
ctggctagcgccaccatggcttgccccgtcccctgcagctcccccccctggagcggctgacactggatgac
aagaagcccctggacgtgctcatcagcgccaccggcctgtggatgagccggacaggcaccctgcacaagatt
aagcaccacgaggtgagccggagcaagatctacattgagatggcctgcggggatcacctcgtggtgaacaac
tctaggagctgccggacagccagggccttcaggcaccacaagtacaggaagacatgcaagcggtgcagggtg
tccgatgaggacatcaacaactttctgacccggagcacagagtccaagaacagcgtgaaggtgagggtggtg
agcgcccccaaggtgaagaaggccatgcccaagagcgtgtccaggcccctaagcctctggagaactccgtg
agcgccaaggccagcaccaacaccagccgctccgtgcccagccctgccaagtccaccccaaacagcagcgtg
cccgcctccgcccccgcccctccctgacccggagccagctcgatcgggtggaggccctgctgagccctgag
gataagatttccctgaacatggccaagcccttcagggagctggagcctgagctggtcacaaggcggaagaac
gatttccagaggctgtacacaaacgaccgggaggactacctggggaagctggagcgggacattaccaagttc
ttcgtcgataggggcttcctggagatcaagagccccatcctgatccctgccgagtacgtcgagcggatgggc
atcaacaacgacaccgagctgagcaagcagatctttcgggtggacaagaacctctgcctgaggcccatgctg
gccccaactctgtataactacctccggaagctggatcggattctgcccggccccatcaagatctttgaggtc
ggcccctgctacaggaaggagagcgatgggaaggagcacctggaggagtttaccatggtgagctttggccag
atggggtccgggtgcacaagggagaacctggaggccctgatcaaggagttcctggactacctggagattgac
ttcgagatcgtgggcgactcctgtatggtgtacggggacaccctggatatcatgcacggcgatctggagctc
agcagcgccgccgtggggcccgtgtccctggatagggagtggggcattgataagcccacaatcggggccggg
ttcggcctggagaggctgctgaaggtcatgcacggctttaagaacattaagcgggcctcgcggagcgagtcc
tactacaacggcattagcaccaacctgtaagaattcaacgcgttaagtcgacaatcaacctctggattacaa
aatttgtgaaagattgactggtattcttaactatgttgctccttttacgctatgtggatacgctgctttaat
gcctttgtatcatgctattgcttcccgtatggctttcattttctcctccttgtataaatcctggttgctgtc
tctttatgaggagttgtggcccgttgtcaggcaacgtggcgtggtgtgcactgtgtttgctgacgcaacccc
cactggttggggcattgccaccacctgtcagctcctttccgggactttcgctttccccctccctattgccac
ggcggaactcatcgccgcctgccttgcccgctgctggacaggggctcggctgttgggcactgacaattccgt
ggtgttgtcggggaaatcatcgtcctttccttggctgctcgcctgtgttgccacctggattctgcgcgggac
gtccttctgctacgtcccttcggccctcaatccagcggaccttccttcccgcggcctgctgccggctctgcg
gcctcttccgcgtcttcgccttcgccctcagacgagtcggatctccctttgggccgcctccccgcgtcgact
ttaactcgagtctagagggcccgtttaaacccgctgatcagcctcgactgtgccttctagttgccagccatc
tgttgtttgcccctcccccgtgccttccttgaccctggaaggtgccactcccactgtcctttcctaataaaa
tgaggaaattgcatcgcattgtctgagtaggtgtcattctattctggggggtggggtggggcaggacagcaa
gggggaggattgggaagacaatagcaggcatgctggggatgcggtgggctctatggcttctgaggcggaaag
aaccagatctcgggcaggaagagggcctatttcccatgattccttcatatttgcatatacgatacaaggctg
ttagagagataattagaattaatttgactgtaaacacaaagatattagtacaaaatacgtgacgtagaaagt
aataatttcttgggtagtttgcagttttaaaattatgttttaaaatggactatcatatgcttaccgtaactt
gaaagtatttcgatttcttggctttatatatcttgtggaaaggacgaaacaccggaaacctgatcatgtaga
tcgaacggactctaaatccgttcagccgggttagattcccggggtttccgtttttaatatttgcatgtcgc
tatgtgttctgggaaatcaccataaacgtgaaatccctatcagtgatagagacttataagttccctatcagt
gatagagacaccgggggtggatcgaatagatcacacggactctaaatccgtgcaggcgggtgaaactcccg
cacccccgtttttcagctggggctctaggggtatccccacgcgccctgtagcggcgcattaagcgcaag
ccctgccatagccactacgggtacgtagccaaccactagaactatagctagagtcctgggcgaacaaacgat
gctcgccttccagaaaccgaggatgcgaaccacttcatccggggtcagcaccaccggcaagcgccgcgacg
gccgaggtctaccgatctcctgaagcaagggcagatccgtgcacagcaccttgccgtagaagaacagcaagg
ccgccaatgcctgacgatgcgtggagaccgaaaccttgcgctcgttcgccagccaggacagaaatgcctcga
cttcgctgctgcccaaggttgccgggtgacgcacaccgtggaaacggatgaaggcacgaacccagttgacat
aagcctgttcggttcgtaaactgtaatgcaagtagcgtatgcgctcacgcaactggtccagaaccttgaccg
aacgcagcggtggtaacggcgcagtggcggttttcatggcttgttatgactgtttttttgtacagtctatgc
ctcgggcatccaagcagcaagcgcgttacgccgtgggtcgatgtttgatgttatggagcagcaacgatgtta

cgcagcagcaacgatgttacgcagcagggcagtcgccctaaaacaaagttaggtggctcaagtatgggcatc
attcgcacatgtaggctcggccctgaccaagtcaaatccatgcgggctgctcttgatcttttcggtcgtgag
ttcggagacgtagccacctactcccaacatcagccggactccgattacctcgggaacttgctccgtagtaag
acattcatcgcgcttgctgccttcgaccaagaagcggttgttggcgctctcgcggcttacgttctgcccaag
tttgagcagccgcgtagtgagatctatatctatgatctcgcagtctccggcgagcaccggaggcagggcatt
gccaccgcgctcatcaatctcctcaagcatgaggccaacgcgcttggtgcttatgtgatctacgtgcaagca
gattacggtgacgatcccgcagtggctctctatacaaagttgggcatacgggaagaagtgatgcactttgat
atcgacccaagtaccgccacctaacaattcgttcaagccgagatcggcttcccggccgcggagttgttcggt
aaattgtcacaacgccgcgaatatagtctttaccatgcccttggccacgcccctctttaatacgacgggcaa
tttgcacttcagaaaatgaagagtttgctttagccataacaaaagtccagtatgctttttcacagcataact
ggactgatttcagtttacaactattctgtctagtttaagactttattgtcatagtttagatctattttgttc
agtttaagactttattgtccgcccacacccgcttacgcagggcatccatttattactcaaccgtaaccgatt
ttgccaggttacgcggctggtctatgcggtgtgaaataccgcacagatgcgtaaggagaaaataccgcatca
ggcgctcttccgcttcctcgctcactgactcgctgcgctcggtcgttcggctgcggcgagcggtatcagctc
actcaaaggcggtaatacggttatccacagaatcaggggataacgcaggaaagaacatgtgagcaaaaggcc
agcaaaaggccaggaaccgtaaaaaggccgcgttgctggcgtttttccataggctccgcccccctgacgagc
atcacaaaaatcgacgctcaagtcagaggtggcgaaacccgacaggactataaagataccaggcgtttcccc
ctggaagctccctcgtgcgctctcctgttccgaccctgccgcttaccggatacctgtccgcctttctccctt
cgggaagcgtggcgctttctcatagctcacgctgtaggtatctcagttcggtgtaggtcgttcgctccaagc
tgggctgtgtgcacgaaccccccgttcagcccgaccgctgcgccttatccggtaactatcgtcttgagtcca
acccggtaagacacgacttatcgccactggcagcagccactggtaacaggattagcagagcgaggtatgtag
gcggtgctacagagttcttgaagtggtggcctaactacggctacactagaagaacagtatttggtatctgcg
ctctgctgaagccagttaccttcggaaaaagagttggtagctcttgatccggcaaacaaaccaccgctggta
gcggtggtttttttgtttgcaagcagcagattacgcgcagaaaaaaaggatctcaagaagatcctttgatct
tttctacggggtctgacgctcagtggaacgaaaactcacgttaagggattttggtcatgagattatcaaaaa
ggatcttcacctagatccttttaaattaaaaatgaagttttaaatcaatctaaagtatatatgagtaaactt
ggtctgacagttaccaatgcttaatcagtgaggcacctatctcagcgatctgtctatttcgttcatccatag
ttgcctgactccccgtcgtgtagataactacgatacgggagggcttaccatctggccccagtgctgcaatga
taccgcgagacccacgctcaccggctccagatttatcagcaataaaccagccagccggaagggccgagcgca
gaagtggtcctgcaactttatccgcctccatccagtctattaattgttgccgggaagctagagtaagtagtt
cgccagttaatagtttgcgcaacgttgttgccattgctacaggcatcgtggtgtcacgctcgtcgtttggta
tggcttcattcagctccggttcccaacgatcaaggcgagttacatgatcccccatgttgtgcaaaaaagcgg
ttagctccttcggtcctccgatcgttgtcagaagtaagttggccgcagtgttatcactcatggttatggcag
cactgcataattctcttactgtcatgccatccgtaagatgcttttctgtgactggtgagtactcaaccaagt
cattctgagaatagtgtatgcggcgaccgagttgctcttgcccggcgtcaatacgggataataccgcgccac
atagcagaactttaaaagtgctcatcattggaaaacgttcttcggggcgaaaactctcaaggatcttaccgc
tgttgagatccagttcgatgtaacccactcgtgcacccaactgatcttcagcatcttttactttcaccagcg
tttctgggtgagcaaaaacaggaaggcaaaatgccgcaaaaaagggaataagggcgacacggaaatgttgaa
tactcatactcttcctttttcaatat
